# Programmable artificial RNA condensates in mammalian cells

**DOI:** 10.64898/2026.01.28.702393

**Authors:** Shiyi Li, Yuna Kim, Kevin Wang, Eric John Payson, Anli A. Tang, Maria Villalba Nieto, Dino Osmanovic, Madison Yang, Diego Dilao, Alexandra Bermudez, Wen Xiao, Melody M.H. Li, Neil Y.C. Lin, Kathrin Plath, Douglas L. Black, Elisa Franco

## Abstract

Artificial biomolecular condensates have emerged as powerful tools to control cellular behaviors. Here we introduce a method to build artificial condensates within living mammalian cells through the design of modular RNA motifs formed by a single, short strand of RNA. These condensates emerge spontaneously, creating RNA-rich compartments that remain separated from their surrounding environment. The RNA sequences include stem-loop domains that fold as the RNA is transcribed, and then condense in the nucleus and cytoplasm through loop-loop interactions. These sequences can be optimized and diversified, enabling the generation of distinct, non-mixing condensate populations and the programmable control of their subcellular localization. The RNA motifs can also be modified to recruit small molecules, proteins, and RNA molecules in a sequence-specific manner to the RNA-rich phase. By introducing additional RNAs that link two distinct types of condensates, we can create droplets with multiple subcompartments, whose organization can be controlled by tuning the stoichiometry of different RNA sequences. These artificial condensates provide a versatile platform for studying and manipulating molecular functions inside living cells.

---

Biomolecular condensates are associated with a multitude of processes in living cells, including gene expression, metabolism, and diseases^1^. There is growing interest in methods for producing artificial condensates with controllable composition, viscoelastic properties, and subcellular location, as a means for manipulating cellular functions^2^. One approach to building artificial condensates is to establish weak interactions among engineered proteins that carry intrinsically disordered domains^3,4^. Similarly, RNA molecules designed to include multiple, short sequence repeats can form weak, easily reconfigurable bonds that yield artificial RNA condensates^5–8^. A notable disadvantage of these interactions is their promiscuity, which makes it difficult to build condensates with a compositional identity, and therefore with exclusive functions. An alternative approach is to take advantage of multivalent molecules with site-specific interactions^9,10^.

We and others recently described *in vitro* assembly of RNA condensates from short single strands of RNA (100-200 nucleotides-long) that operate as multivalent particles^11,12^. These structural motifs, termed single-stranded RNA (ssRNA) nanostars, consist of at least three tandem stem-loops that fold during transcription, functioning as “arms” (Fig. 1A). The arms interact through sequence-specific binding of their loop domains, known as kissing loops, which are typically palindromic and identical on each arm. Kissing loops can be designed to be non-interacting, so distinct nanostars can produce distinct condensates that do not mix^11,12^. The modular design of these nanostars allows for the inclusion of additional arms, enabling the addition of domains for the recruitment of small molecules and proteins to the dense phase, without compromising condensate formation^11,12^. Here we demonstrate that ssRNA nanostars can generate condensates within living mammalian cells with controlled mixing patterns, and that the interactions and localization of these condensates can be tuned by modifying the nanostar design.

**Figure 1.**
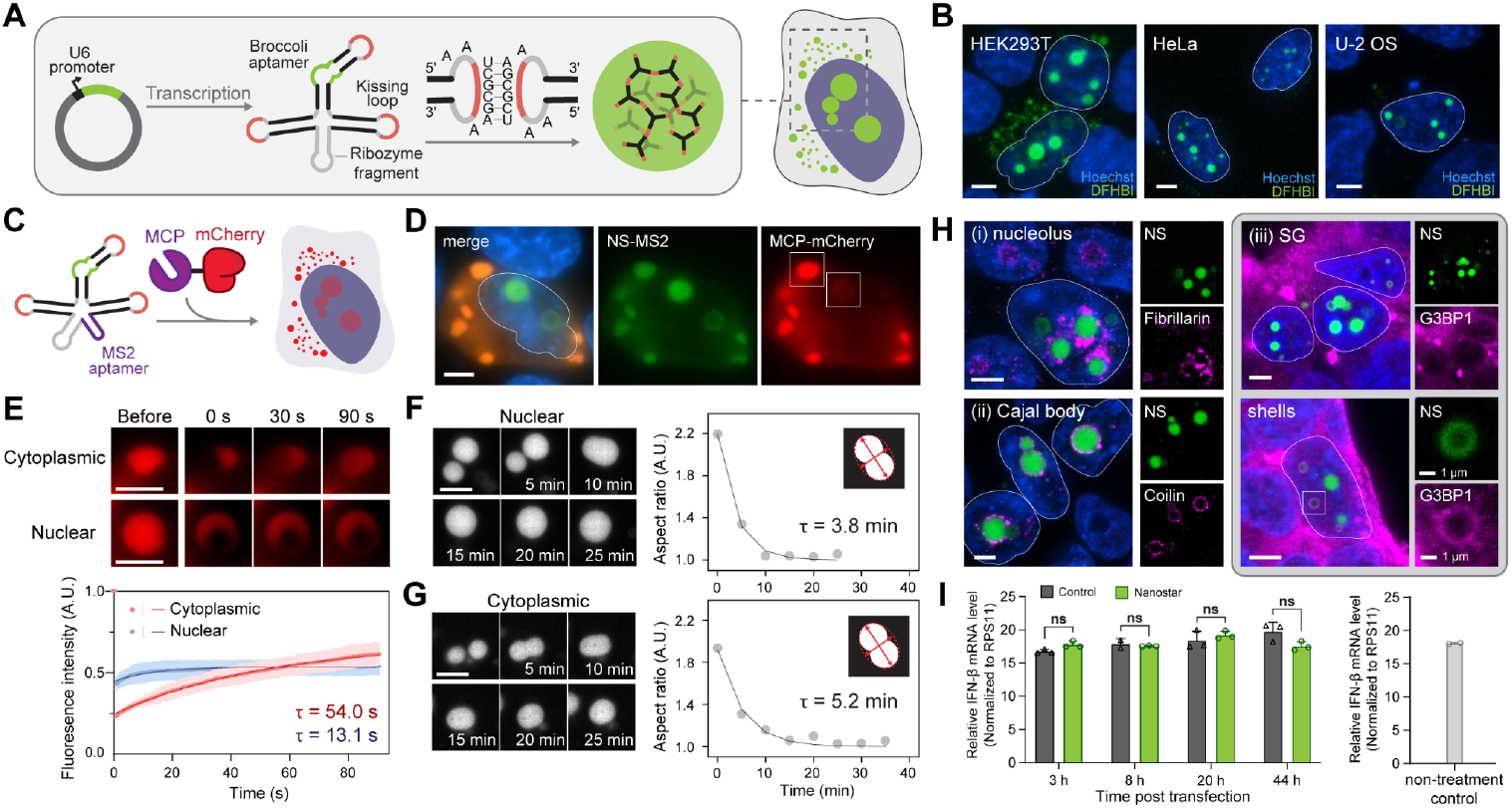
Circularized RNA nanostars generate condensates in different cell lines. **A**, We transfected ssRNA nanostar in mammalian cells using the Tornado plasmid, which results in a circularized motif. ssRNA nanostars form condensates through their kissing loop interactions. **B**, Representative confocal microscopy images of RNA condensate forming in HEK293T, HeLa, and U-2 OS cells. Nuclei of condensate-expressing cells are outlined. **C**, Schematic of RNA nanostars recruiting MCP-mCherry through the addition of an MS2 aptamer as an additional arm. **D**, Split channel fluorescent images of a HEK293T cell co-transfected with MCP-mCherry and MS2-nanostar stained with DFHBI. FRAP analysis was performed on condensates highlighted with white squares and shown in E. The nucleus is outlined in the merged channel image. **E**, Fluorescence images (top) and pixel intensity profile (bottom) at different time points after photobleaching. Condensates were imaged before, and every 200 ms for 90 s after bleaching. Blue and orange dots indicate the mean pixel intensity of the bleached area at the corresponding time point. Error bars indicate standard error. The dark blue and red lines indicate the fitted curve from equation 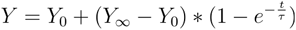 Mean and standard error were calculated from 3 fields of view each belonging to one replicate. **F, G**, Fluorescence images (left) and time-dependent change in aspect ratios (right) demonstrating coalescence of nuclear (F) or cytoplasmic (G) condensates formed in HEK293T cells transfected to express Broccoli-labeled nanostars. Aspect ratios of condensates (gray dots) are computed as the major and minor axes ratio from the best-fit-ellipses. The dashed line is a fit of the exponential function . 1 + Ae^(−t/τ)^. **H**, Co-localization of condensates with other cellular membraneless organelles in HEK293T cells. Cells were stained with Hoechst (blue), DFHBI (green), primary antibodies and Alexa Fluor™ 647 labeled secondary antibodies (magenta). **I**, Quantitative PCR (qPCR) results measuring the expression level of IFN-β in nanostar-expressing cells (left, green) and lipofectamine-treated cells (left, dark grey) at various time points post-transfection or in non-treated cells (right, light grey). Statistical significance determined by two-way ANOVA followed by multiple comparison tests show no significance. Nuclei of condensate expressing-cells are outlined in the merged channel image. All cellular expression results were captured 48 hours after transfection except for the time-lapse results. Micrographs in panel B are z-projections of stack images captured by a confocal microscope. Micrographs in panel F, G, and H are single slices of the stack images. Micrographs in panels D, and E are captured using an epifluorescence microscope. All experiments were replicated three times; images are representative examples. Scale bar, 5 μm.

## RNA nanostars produce condensates that recruit target molecules

We produced ssRNA nanostars in HEK293T cells using the TORNADO expression system developed by Litke et al.^13^, which uses the U6 promoter and includes domains for spontaneous RNA circularization to extend the RNA’s half-life (Supplementary Fig. 1 and 2, Supplementary Table 1). We started with a nanostar design that carries three 15-nucleotide (nt) long arms and each with a 6-nt long kissing loop UCGCGA (KL variant A), shown in Fig. 1A (design 15nt-3A-Br). We included the Broccoli aptamer in one of the arms, to visualize the expressed RNA in cells by adding its fluorogenic ligand 3,5-Difluoro-4-hydroxybenzylidene imidazolinone (DFHBI) to the media^14^. This nanostar design generates condensates during *in vitro* transcription at constant temperature (Supplementary Fig. 3), and we observed comparable structures forming in HEK293T, HeLa, and U-2 OS cells transfected with the nanostar-carrying plasmids (Fig. 1B, Supplementary Fig. 4 and 5). Transfection in HEK293T cells yields a higher expression level when compared with the other two cell types, because of the presence of the SV40 large T antigen that enhances gene expression^15,16^. Flow cytometry control experiments confirmed that expression of Broccoli alone with the Tornado system results in lower granularity when compared to nanostar-expressing cells (Supplementary Fig. 6); microscopy further confirmed diffuse fluorescence in the cytoplasm, with few puncta in the nucleus and shells that may correspond to circularized RNA colocalized with other cellular components^13,17^ (Supplementary Fig. 7A). Replacement of at least one kissing loop with a polyA sequence disrupts the formation of full droplets, although shells persist, confirming that condensation is driven by loop-loop interactions (Supplementary Fig. 7B).

The modular structure of RNA nanostars makes it possible to include additional, non-interacting arms for the recruitment of guest molecules through aptamer domains. To illustrate this, we included the MS2 aptamer^18^ as an additional arm, with the goal of recruiting to the condensates a reporter protein carrying an MS2 coat protein (MCP) domain. In our experiments we co-transfected two plasmids into cells, one encoding the modified RNA nanostar and the other encoding an MCP-mCherry reporter serving as guest molecule for the condensate (Fig. 1C, Supplementary Fig. 8 and 9), and we tested the colocalization of Broccoli and mCherry signals. Live-cell imaging confirmed successful mCherry recruitment into RNA condensates (Fig. 1D, Supplementary Fig. 10-12), though mCherry remained predominantly cytoplasmic due to the absence of a nuclear localization signal (NLS). Although the overall and relative expression level of MCP-mCherry and nanostar influence the shape and cellular localization of condensates, recruitment efficiency estimated from the partition coefficient remains similar (Supplementary Fig. 10B-E). In contrast, condensates formed from nanostars lacking the MS2 domain did not affect the spatial localization of mCherry (Supplementary Fig. 9).

## Diffusivity and viscosity of RNA condensates

Using fluorescence recovery after photobleaching (FRAP), we estimated the diffusion timescale for mCherry guest protein recruited to the dense phase. MCP-mCherry diffusion from cytoplasmic condensates exhibits recovery with a fitted time constant of 54 s, while nuclear condensates show faster but minimal recovery with a fitted time constant of 13.1 s (Fig. 1E and Supplementary Fig. 11). The lower recovery extent in the nucleus could be attributed to the lower abundance of mCherry (which lacks a NLS), and higher mCherry fluorescence density correlated with greater recovery (Supplementary Fig. 12). While mCherry does not participate directly in condensation interactions, it represents a cargo protein that may hinder RNA-RNA interactions, leading to increased reporter diffusivity and thus faster recovery.

FRAP analysis of nuclear and cytoplasmic condensates using Broccoli reveals faster recovery of this small molecule fluorogenic guest, respectively *τ*=7.8*s* and *τ* = 10.8*s*(Supplementary Fig. 13A-D). *In vitro* experiments with Broccoli, where condensates were produced during transcription in buffer at constant temperature, exhibited a slightly slower recovery profile when compared to that measured *in vivo*, with a fitted time constant of 18.2*s* (Supplementary Fig. 14A). Because small-molecule ligands bind their RNA aptamer non-covalently (Broccoli *K*_*d*_ ≈300*nM*), the observed recovery is likely dominated by dynamic ligand exchange rather than RNA diffusion. Nuclear condensates labeled with Pepper, another fluorogenic aptamer^17,19^ that produces red fluorescence upon the addition of (4-((2-hydroxyethyl)(methyl)amino)-benzylidene)-cyanophenylacetonitrile 620 (HBC620), showed distinct recovery with a larger fitted time constant of 79.5 s, consistent with a lower *K*_*d*_ ≈3.5*nM*^20^ when compared to Broccoli (Supplementary Fig. 13E-F). Incomplete recovery was observed with all fluorescent reporters, suggesting that RNA condensates might exhibit partially solid-like properties. Condensates formed *in vitro* also underwent fusion with slow relaxation dynamics, with one representative event exhibiting a relaxation timescale of τ ≈ 16*h*(Supplementary Fig. 15). In contrast, fusion dynamics in living cells are faster with τ = 3.8*min* for a nuclear event, and τ = 5.2 *min* for a cytoplasmic event (Fig. 1F and G, Supplementary Fig. S16-19). Nuclear events appeared faster than cytoplasmic events, potentially reflecting differences in the RNA-binding protein (RNP) environment between the nucleus and cytoplasm, which may influence condensate interactions. Although faster than in vitro, the fusion dynamics of RNA condensates are slower than those of some biomolecular condensates such as P granules and stress granules, which fuse on the timescale of seconds^21,22^, and are comparable to the dynamics of the nucleolus^23^. Collectively FRAP and fusion results indicate that RNA condensates are comparatively more viscous. Interestingly, we also observed a “splitting” event between nuclear condensates (Supplementary Fig. S19), suggesting that their behavior may be influenced by other biological components.

## RNA condensates interact with nuclear organelles but do not elicit an immune response

We examined the crosstalk between our RNA condensates with other membraneless organelles through immunostaining of relevant proteins (Fig. 1H, Supplementary Fig. 20-24). Immunostaining of fibrillarin indicates accumulation of nuclear RNA nanostars in the nucleolar region (Fig. 1H (i), Supplementary Fig. 20A). In these cells, fibrillarin was recruited to the condensate surface but it remained excluded from condensates, demonstrating selective partitioning of molecules. Some RNA nanostar variants might be favored for nucleolar localization (Supplementary Fig. 20B). Similarly, coilin also aggregates on the condensates surface, suggesting recruitment of Cajal bodies (Fig. 1H (ii), Supplementary Fig. 21). Cells producing high levels of RNA condensates showed an increased likelihood of stress granule (SG) formation when compared with cells expressing lower levels, as indicated by staining of G3BP1, a key SG component (Fig, 1H (iii), top, Supplementary Fig. 22). However, colocalization between G3BP1 and nanostars was only observed in a few cells with high cytoplasmic RNA condensate formation, indicating that total cytosolic RNA increase might result in G3BP1 condensing, possibly with different RNA molecules^24,25^ (Supplementary Fig. 22E). Additionally, in some nuclei, we found that G3BP1 colocalizes with shells, not with condensates, and wraps on the outside of these shells (Fig. 1H (iii), bottom, Supplementary Fig. 23). Formation of shells has also been reported in other systems producing circularized RNA, indicating that this structure is not specific to nanostar condensates^17^ (Supplementary Fig. 7 and 22). More work is needed to determine whether SGs in nanostar-expressing cells arise from direct interactions between RNPs such as G3BP1 and RNA nanostars, or whether they are triggered by metabolic stress caused by elevated cytoplasmic RNA levels^24–27^. We found no colocalization with P-bodies and nuclear speckles (Supplementary Fig. 24). No immunogenic response was observed, as indicated by comparable levels of interferon-β (IFN-β) and the IFN-stimulated genes ISG15 and IFIT1 in cells with and without condensates (Fig. 1I, Supplementary Fig. S25, Supplementary Table 4).

## RNA nanostar design determines the subcellular localization of condensates

Because RNA nanostars are produced in the nucleus and then exported to the cytoplasm, their local concentration is determined by the relative speed of transcription, export from the nucleus, and degradation. These processes crucially affect condensate formation, which occurs only if the nanostar concentration exceeds a critical value *C*^**\***28^. We illustrate the importance of production and export rates through a simple compartment model (Fig. 2A), which does not attempt to discriminate between active and passive transport, and neglects nuclear degradation. In the model our molecular subunits (RNA nanostars) are produced at a rate of inside the nucleus and then exported to the cytoplasm at a constant rate :

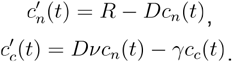

**Figure 2.**
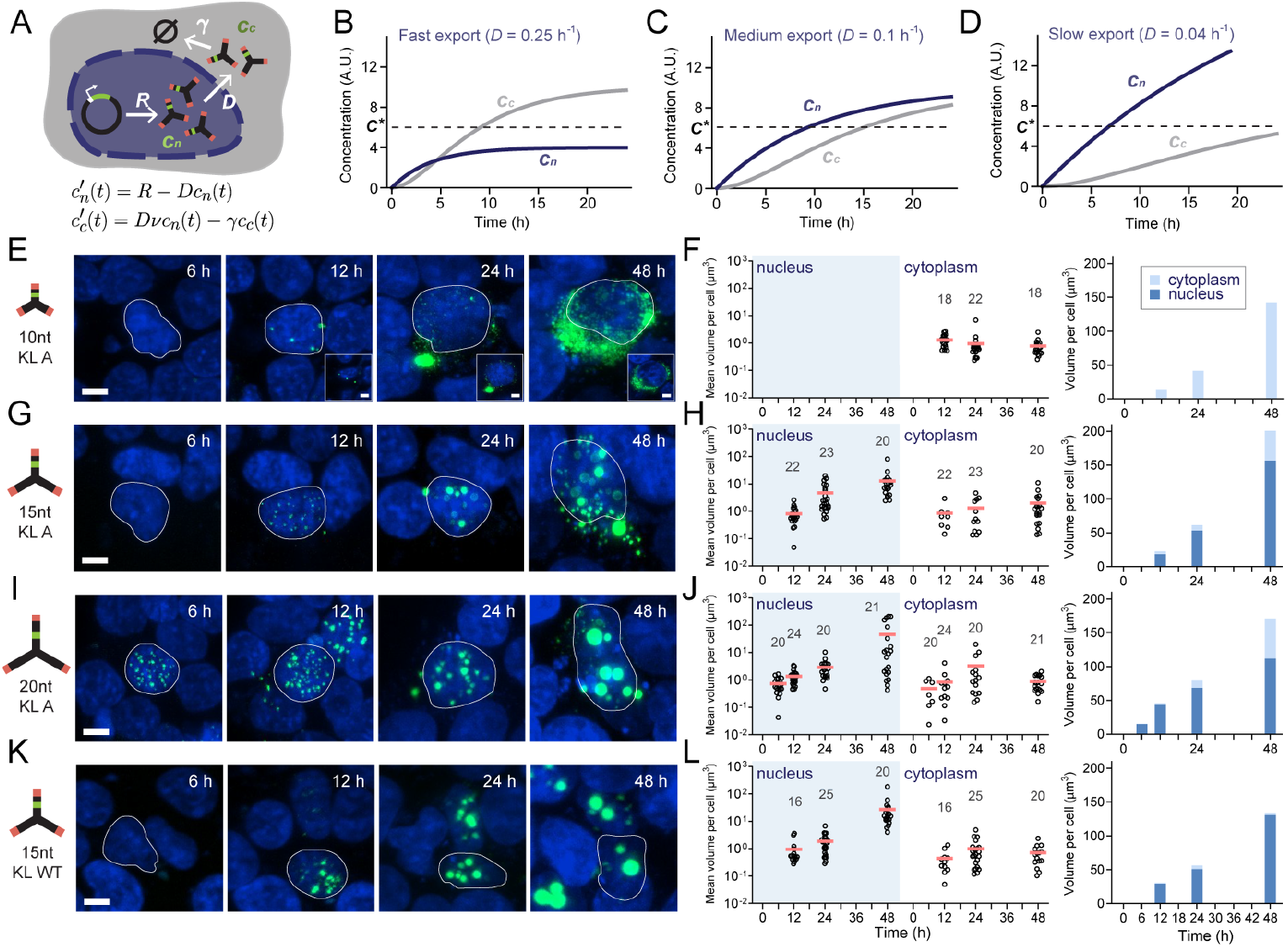
Nuclear and cytoplasmic condensate formation depends on nanostar design. **A**, Schematic depicting parameters determining nanostar concentration inside or outside the nucleus, where R is the transcription rate,D is the nuclear export rate γ is the degradation rate, ν is the volume ratio correction term, defined as V_n_/V_c_,C_n_, is the nanostar concentration inside of the nucleus,C_c_ is the nanostar concentration inside of the cytoplasm, and ∅ indicates RNA degradation products. **B, C, D**, Simulations generate different concentration profiles ( C_n_, nuclear. C_c_, cytoplasmic) with varied export rate ( D ), while other parameters remain constant with R = 1, γ = 0.2, and ν=2., and . Phase separation is initiated, and visible condensates begin to form when the concentration surpasses the critical concentration (C^*^). While the transcription rate is determined by the amount of plasmid entering the cell, the nuclear export rate depends on the size of the nanostar. **E, G, I, K** Representative z-projections of confocal microscopy images showing condensate growth over the course of 48 hours from different nanostar designs. Cells were stained with Hoechst (blue) and DFHBI (Green). For each design, the nucleus of the same cell is outlined at different time points. Inserts in E are single slice midplane images, showing all condensates localized outside of the nucleus. Additional midplane images are in Fig. S29. **F, H, J, L**, Dot plots showing temporal evolution of average condensate volume per cell inside the nucleus and the cytoplasm. Above each dot plot, we report the number of sampled cells at each time point, across multiple fields of view, from three replicates. Bar plots show the total condensate volume averaged across analyzed cells. Scale bar, 5 μm.

Here *γ* is the degradation rate constant in cytoplasm and *ν* is a correction term accounting for the change in volume ratio, defined as *V*_***n***_*/V*_***c***_. The nuclear production rate of nanostars in each cell depends primarily on the amount of plasmids delivered into the cell during transfection, a factor that should be independent of small variations in the nanostar design. Similarly, the degradation rate should only depend on the nanostar concentration. In contrast, the specific structural design features of the nanostar can affect the export constant, which is governed by the size-sensitive permeability of the nuclear pore. Consequently, the export rate is sensitive to changes in subunit design, meaning that tuning the nanostar size and intermolecular affinity offers a way to modulate nuclear export^29^. In particular, depending on the export rate of subunits, the critical concentration can be reached at different times in the nucleus and in the cytoplasm, or may never be reached, as shown in Fig. 2B-D.

To test this, we sought to affect nuclear export rate by modifying the nanostar arm length^11^. We designed nanostars with arm length ranging between 10 and 20 nt, and measured the RNA condensate volume over time, 6-, 12-, 24-, and 48-hours post-transfection (Fig. 2E-J, Supplementary Fig. 26-27)^30^. We found that 10-nt arm nanostars (arm length 3.4 nm, molecular weight (MW) 56.8 kD) form condensates exclusively in the cytoplasm as they likely diffuse quickly through the nuclear pore complex (Fig. 2E and F, Supplementary Fig. 27E), demonstrating a similar behavior as simulated in Fig. 2B. These nanostars do not colocalize with nucleoli, nor with Cajal bodies (Supplementary Fig. 20B and 21B), likely due to their rapid export to the cytoplasm. Nanostars with 15-nt arm (arm length 5.1 nm, MW 66.3 kDa) and 20-nt arm (arm length 6.8 nm, MW 75.8 kDa), having sizes close to the nucleus’ permeability barrier, form condensates inside the nucleus first, and the cytoplasmic fraction increases over time (Fig. 2G and J,Supplementary Fig. 27F, D). Nanostars with 20-nt condensed at an earlier time point as the growth rate of condensates scales with nanostar sizes, consistent with illustrative simulation results in Fig. 2C and with our related findings using *in vitro* DNA nanostars ^31^. In addition to being influenced by subunit size, export rate could also be impacted by subunit interactions, which can promote the assembly of clusters before nuclear export. To test this, we changed the 15 nt arm nanostar KL A (UCGCGA), to the HIV wild type kissing loop (WT, GCGCGC), the strongest possible variant ^32^. We observed a significant increase in the nuclear condensate ratio, indicating higher nuclear retention driven by faster complex assembly (Fig. 2K and 2L, Supplementary Fig. 27).

We then sought to further elucidate how design parameters influence condensate localization, abundance, and morphology, by systematically varying size (arm length), valency (number of arms) and avidity (kissing loop strength) of the nanostars^31^. We provide a comprehensive overview of our results in Fig. 3. For each variant, we measured corresponding condensate volume, number, and cellular localization 48 hours after transfection (Supplementary Fig. 29-33). Confirming the results of our kinetic experiments, nanostars differing by arm length produced condensates with distinct localization and size (Fig. 3A and B). Nanostars with 10-nt arms produced condensates localized exclusively in the cytoplasm, while nanostars with 15-nt, 20-nt, and 25-nt long arms generated condensates both in the cytoplasm and in the nucleus, where there were fewer but larger condensates. Next, we varied the nanostar arm number between 2 and 4, and found that the average nuclear condensate volume per cell correlated with the arm number (Fig. 3C and D). A reduction of valency by replacing kissing loop sequences with the same number of adenine bases yielded similar effects as deleting arms (Supplementary Fig. 7D,E). We also found that nanostars with more arms yielded fewer but larger condensates in the nucleus, and correspondingly, more small puncta in the cytoplasm. More arms likely accelerated nuclear aggregation of nanostars, thereby slowing down transport to the cytoplasm. At the same time, the increased valency enhanced nucleation in the cytoplasm, leading to the formation of more small puncta. Finally, we investigated the impact of kissing loop interaction strength on condensate formation through four variants, ranked from the weakest (F variant, UAUAUA) to the strongest (WT variant, GCGCGC) (Fig. 3E and F, Supplementary Table 2). The strongest variant (WT) yielded large condensates in the nucleus and a higher number of smaller puncta in the cytoplasm, resembling the behavior of the 4-arm nanostar. In contrast, weak kissing loop variants E and F primarily formed nuclear shells. Overall, larger and fewer condensates accumulated in the nucleus as we increased arm number and strength of the kissing loops (Fig. 3G, Supplementary Fig. 30, and Supplementary Table 3), likely due to the rapid formation of nuclear aggregates which hindered export.

**Figure 3.**
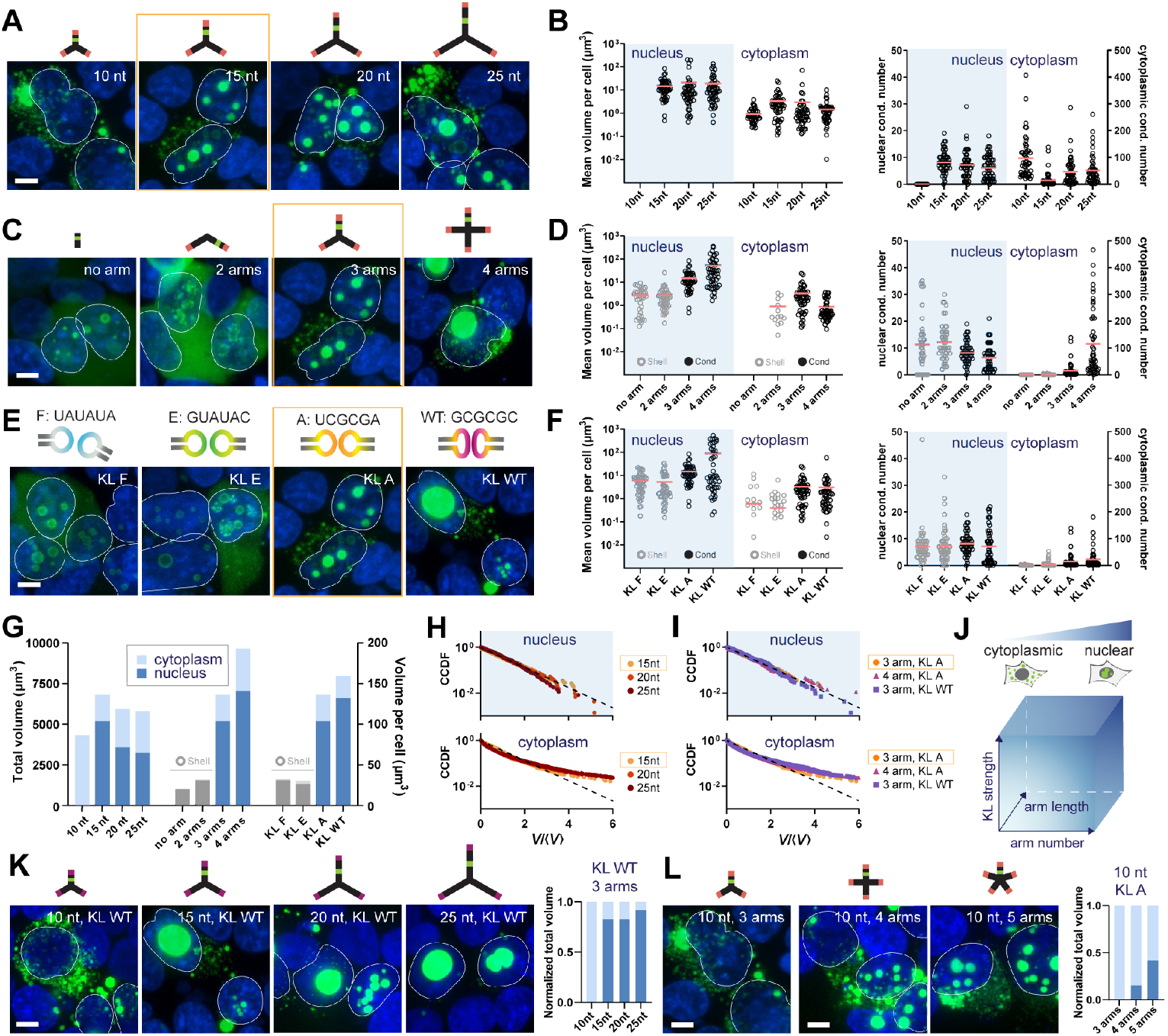
Engineering condensate volume distribution and localization through nanostar design. **A, C, E**, Representative z-projection of confocal microscopy images showing condensates produced by RNA nanostar variants with different arm length (A), arm number (C), and kissing loop sequence (E). Cells were stained with Hoechst (blue) and DFHBI (green). The yellow box highlights the same design. The nuclei of condensate-expressing cells are outlined for convenience. **B, D, F**, Left: Average condensates’ volume in the nucleus and cytoplasm of individual cells on a log scale. Right: Number of condensates in the nucleus or cytoplasm measured in individual cells. Each black circle represents one cell. Red lines indicate the mean. Statistical significance annotations are in Fig. S30. **G**, Bar chart comparing the total condensate volume for 50 cells, and average volume per cell. **H, I**, Nuclear condensate volume is exponentially distributed, while cytoplasmic condensates display power-law-like distribution for nanostars with different arm lengths (H), arms numbers and kissing loops (I). To generate cumulative distribution functions (CCDFs) of condensate volumes, condensates were individually segmented in three dimensions and rescaled by the mean condensate volume in each cell. After rescaling, data from all cells were pooled and plotted collectively. Each dot represents a condensate. The yellow box highlights the same nanostar variant. The CCDF was also compared with the expectation for an exponential distribution (dashed line). **J**, Schematic illustrating the correlation between condensate nuclear localization and stronger kissing loops, longer arms, and greater arm numbers. **K, L**, Confocal micrographs (left) show enhanced nuclear localization of condensates with increased arm length (K) or arm number (L). Normalized total condensate volume from fifty cells (right) confirms an increased nuclear ratio (dark blue) compared to cytoplasmic (light blue). All experiments were replicated three times. Scale bar, 5 μm.

Across designs, the size distributions of condensate volumes differ between the nucleus and the cytoplasm, as shown by the complementary cumulative distribution functions (CCDF) in Fig. 3H and I^33^. We normalized condensate volumes by the average condensate volume per cell and found that nuclear condensates follow an exponential distribution, for which the normalized CCDF appears as a straight line with slope −1 on a semilog plot. In contrast, the volume distribution of cytoplasmic condensates is broader than an exponential, and more consistent with a power law. This difference may be explained by faster nanostar production in the nucleus than export to the cytoplasm^33^, consistent with the simple model presented in Fig. 2A. No significant deviation in the CCDF was observed across designs, suggesting comparable nucleation and coalescence time scales. The CCDF of condensate volume measured at different time points (experiments reported in Fig. 2) shows similar trends, with nuclear condensate volume scaling exponentially, and cytoplasmic condensates scaling like a power law across designs (Supplementary Fig. 31).

Overall, these results show that RNA nanostar design provides means to tune the nuclear-to-cytoplasmic distribution of condensates. The proportion of nuclear condensates increases with stronger kissing loops, more arms, and longer arms (Fig. 3J, Supplementary Fig. 32). We further validated this programmable control by (1) maximizing nuclear localization and (2) relocating cytoplasmic-only condensates into the nucleus through sequence modifications. First, we introduced the WT kissing loop to enhance interaction strength and observed almost exclusively nuclear condensates for the 25-nt nanostar (Fig. 3K, Supplementary Fig. 33). Next, we increased the arm number from 3 to 5, reasoning that higher valency and molecular size might synergistically promote nuclear localization. As expected, 10-nt nanostars with more arms formed nuclear condensates whose size and number scaled with arm number (Fig. 3L, Supplementary Fig. 33).

Collectively, these experiments demonstrate that RNA nanostars are a robust motif to build cellular condensates, and that structural and sequence variations make it possible to influence the cellular localization and the size of condensates. Partition coefficient analysis across all designs confirmed higher nuclear enrichment compared to cytoplasmic condensates, while shell-forming constructs consistently displayed low partition coefficients (Supplementary Fig. S34). We verified that variations to the nanostar design do not compromise the ability of condensates to recruit target proteins. We modified nanostars with A and WT kissing loops to include an MS2 aptamer, enabling their corresponding condensates to recruit MCP-mCherry (Supplementary Fig. 35A). FRAP experiments revealed slower mCherry recovery for the WT kissing loop compared to A, indicating that stronger kissing loop interactions result in higher condensate viscosity (Supplementary Fig. 35B). Finally, we observed that, across designs, cells expressing RNA nanostars exhibit an enlarged nucleus, likely as a consequence of increased local osmotic pressure (Supplementary Fig. 36 and 37).

## Sequence design produces diverse condensates with tunable interactions and cellular localization

Diverse RNA nanostars can be created that carry different sequences in their kissing loop domains (Fig. 4, Supplementary Fig. 38). For example, a two-nanostar system can generate condensates *in vitro* where mutual specific interactions are introduced by non-palindromic, complementary kissing loops^11^ (Fig. 4A). To test this in living cells, we labeled these nanostars with distinct fluorogenic aptamers, Broccoli, and Pepper. When only one of these two nanostars was expressed, we observed diffuse fluorescence in the cytoplasm, and the hollow shell structures in the nucleus^13^ (Fig. 4B, top and middle). When both nanostars are co-expressed, condensates exhibiting both Broccoli and Pepper fluorescence appear in the cytoplasm and nucleus, as confirmed by a high Pearson correlation coefficient (PCC). Interestingly, the ratios of Broccoli to Pepper fluorescence signals differ between cytoplasmic and nuclear condensates, as shown by distinct linear relationships in the fluorescence scatter plot (Fig. 4B, bottom): while the Broccoli-carrying nanostar yields condensates both in the nucleus and the cytoplasm, the Pepper-carrying nanostar yields condensates primarily in the cytoplasm. Given the same expected hybridization energy of their kissing loops, this difference is likely attributed to the aptamer which may influence nanostar folding and structure, yielding a faster nuclear export for Pepper-carrying nanostars.

**Figure 4.**
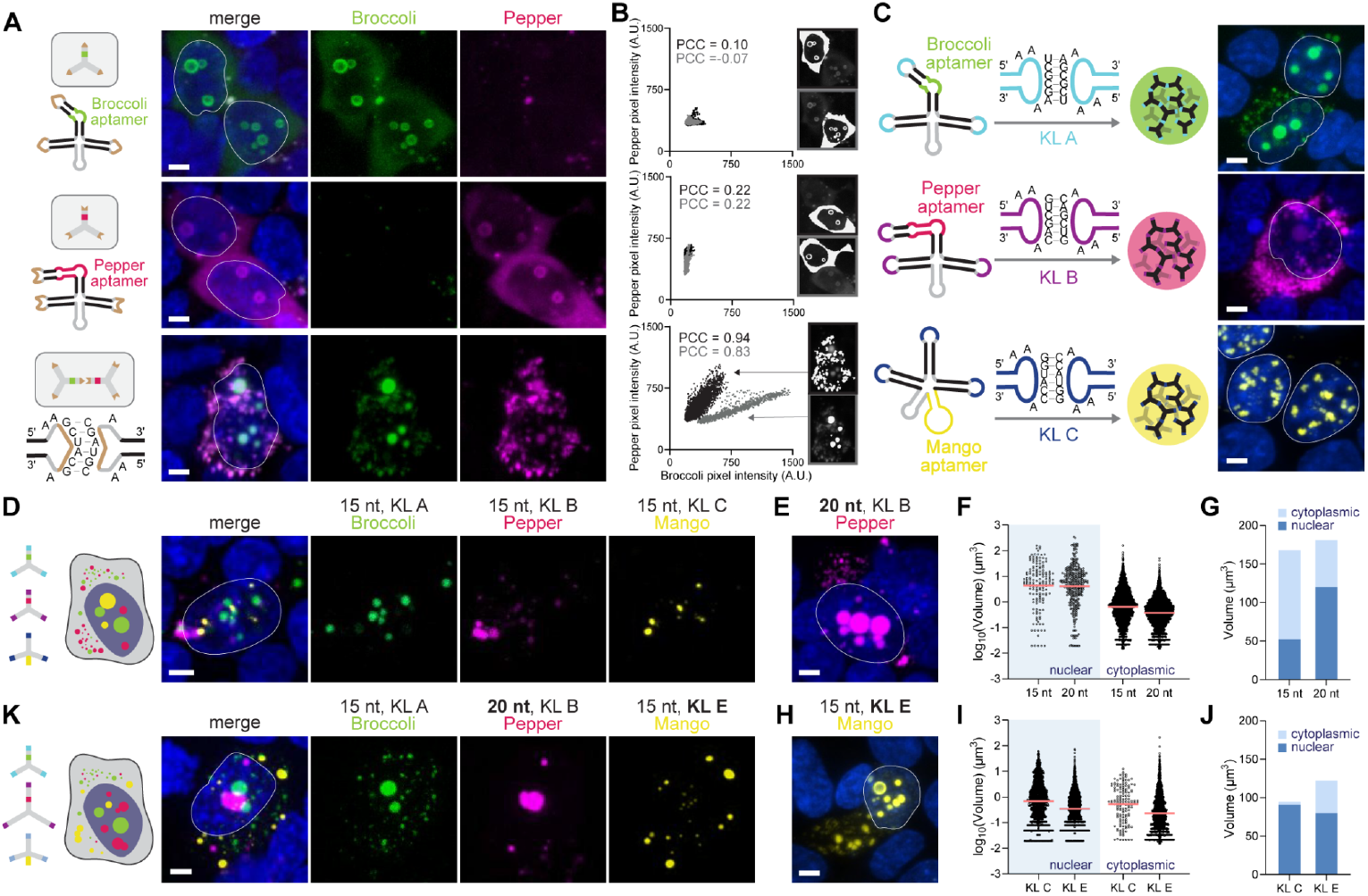
Nanostar kissing loop sequences determine condensate miscibility. **A**, Schematic of nanostars labeled with Broccoli or Pepper interact through complementary but non-palindromic kissing loops. Fluorescence micrographs showing that condensates form only when both nanostars are produced. **B**, Scatter plots showing pixel intensities in the Broccoli and Pepper channels for each pixel within the white-highlighted regions of interest, along with the corresponding Pearson correlation coefficients. In the bottom plot, nuclear and cytoplasmic condensates exhibit distinct correlations, reflecting differences in the stoichiometry of the two nanostars. These variations are likely driven by differences in fluorescence tagging, which may affect nanostar folding and, consequently, their distribution patterns. **C**, Schematic of nanostars with kissing loops A, B, C, respectively labeled with Broccoli, Pepper, or Mango aptamers and fluorescence micrographs of the corresponding condensates in cells. **D**, Distinct condensates that do not mix emerge upon co-transfection of plasmids each carrying one of the 15-nt arm nanostars with kissing loops A, B, C, respectively labeled with Broccoli, Pepper, or Mango aptamers. **E, H**, Confocal fluorescence micrographs of cells expressing modified nanostars tagged with Pepper (longer arms) or Mango (weaker kissing loops) to modify their subcellular location. **F, I**, Dot plots illustrate changes in the distribution of nuclear and cytoplasmic condensate volume in log scale. Each dot represents a condensate. Red lines indicate the mean. **G, J**, Bar charts comparing the average volume of nuclear and cytoplasmic condensates. **K**, We redesigned the cellular localization of distinct condensates using modified Pepper and Mango-tagged nanostars (Broccoli design unchanged). Nuclei are outlined for convenience. HEK293T Cells were stained with Hoechst, DFHBI (for Broccoli), HBC620 (for Pepper), and TO1-B (for Mango). Mango-tagged condensates were imaged after fixing cells, due to poor permeability of the cognate dye into the nucleus. Tri-color images were acquired by live cell imaging, followed by on-stage fixation and imaging of the same field of view. All experiments were replicated three times. Images are z-projections of stack images captured by a confocal microscope. Scale bar, 5 μm.

Leveraging the sequence-specificity of loop-loop interactions, RNA nanostars can be optimized to generate a variety of distinct, non-interacting condensates. To illustrate this we characterized a set of three nanostars, termed A, B, and C^12^, each carrying distinct kissing loops and distinct fluorogenic aptamers, Broccoli, Pepper, and Mango (produces yellow fluorescence upon the addition of TO1-Biotin (TO1-B)) respectively^34^. Each of the A, B, and C kissing loops was designed to engage in homomeric binding but not heteromeric interactions with other kissing loops^12^. Fig. 4C shows that each nanostar generated condensates, with differences in their cellular localization that depended on the choice of aptamer (Supplementary Fig. 39). While nanostar with the Broccoli and Pepper aptamer demonstrated distributions consistent with the variants described earlier, nanostar with kissing loop C and Mango primarily yielded condensate with a less spherical morphology in the nucleus. In this case, the Mango aptamer may introduce additional loop-loop bonds that increase the overall nanostar interaction strength and promote nuclear aggregation. The simultaneous transfection of multiple plasmids each carrying a distinct nanostar with non-interacting kissing loops resulted in the formation of condensates that did not mix (Fig. 4D, Supplementary Fig. 40) and maintained the subcellular localization observed when expressed individually (Fig. 4C).

Taking advantage of the design guidelines identified in Fig. 2 and 3, we then sought to shift the cellular localization of Pepper-tagged condensates (from cytoplasm to nucleus) and Mango-tagged condensates (from nucleus to cytoplasm). To localize Pepper-tagged nanostars to the nucleus, we extended their arm length from 15 nt to 20 nt, and to localize Mango-tagged nanostars to the cytoplasm, we adopted weaker kissing loops, switching from design C (5’-GGUACC, 2 AU pairs and 4 GC pairs) to design E (5’-GUAUAC, 4 AU pairs and 2 GC pairs). With this change, we observed Pepper condensates primarily in the nucleus (Fig. 4E, Supplementary Fig. 41A). While the size of nuclear condensate remained similar, their number significantly increased resulting in a higher nuclear-to-cytoplasmic volume ratio (Fig. 4F and 4G). For Pepper-carrying nanostars, enhancing kissing loop strength from A to WT showed low efficiency in relocating condensates to the nucleus (Supplementary Fig. 42A-C). For Mango-carrying nanostars, weakening the kissing loop from C to E successfully relocated condensates into the cytoplasm, although a substantial fraction still formed in the nucleus (Fig. 4H-J, Supplementary Fig. 41B). Further weakening the kissing loop to F (6 AU pairs) eliminated condensate formation and yielded only nuclear shells (Supplementary Fig 41D). Cotransfection of the new plasmid triplet resulted in distinct condensates with the expected changes in subcellular distribution (Fig. 4K). Collectively, these results indicate that the principles governing the interaction specificity, morphology, and cellular localization of condensates remain valid across variants; however, the efficiency in manipulating condensate properties is affected by details of nanostar and aptamer sequence.

We then demonstrated how to systematically co-localize two non-mixing condensates, or even completely mix the corresponding nanostars, through the production of RNA linker motifs (Fig. 5, Supplementary Fig 43) previously characterized *in vitro*^*12,35*^. A linker is a “chimeric” nanostar in the sense that it includes kissing loops complementary to two of the distinct nanostars allowing them to join. We tested two types of linkers, with two or four arms (Fig. 5A). The colocalization level of distinct nanostars depended on the ratio of the linker plasmid concentration relative to the concentration of plasmids of the distinct nanostars (Fig. 5B, Supplementary Fig. 44). Fig. 5C shows example images of nanostars A and B (20nt arm, labeled with Broccoli and Pepper respectively) in living cells which were transfected with increasing levels of a plasmid encoding a two-arm linker. At nanostar:linker:nanostar plasmid ratios of 2:1:2, 1:1:1, 1:2:1, 1:3:1, nuclear condensates with Broccoli and Pepper fluorescence were colocalized while they remained distinct compartments producing Janus-like morphologies. At higher ratios, nanostars became completely mixed. We observed similar results for the four-arm linker (Fig. 5D), however, a 1:2:1 ratio of nanostar:linker:nanostar was sufficient to produce mixed condensates in the nucleus. Because now each linker carried twice as many arms, this result was consistent when compared with the two-arm linker yielding mixed condensates at a 1:4:1 ratio. In contrast, cytoplasmic condensates exhibited the opposite trend. Compared to the three-arm nanostars, two-arm linkers are smaller, facilitating faster nuclear export and resulting in complete mixing at lower linker ratios in the cytoplasm (Fig. 5C). Meanwhile, four-arm linkers are larger than the three-arm nanostars, leading to slower nuclear export. Consequently, a higher linker ratio was required to achieve complete mixing in cytoplasmic condensates (Fig. 5D). The degree of mixing between the two phases of nuclear condensates was quantified through mixing indices *J*_*R*_ and *J*_*G*_ that measured the fraction of red in green, and green in red condensate^12^ (Fig. 5E and 5F). Compared with *J*_*R*_, index *J*_*G*_ showed more fluctuation due to a lower signal-to-noise ratio of Broccoli compared to Pepper (Supplementary Fig. 45). Notably, at the ratios of 1:4:1 and 1:5:1, we observed non-homogeneous mixing with the four-arm linker. To our surprise, by labeling the two-arm and four-arm linkers with the Broccoli aptamer, we found that four-arm linkers formed condensates when expressed individually (Fig. 5G), suggesting that the non-homogeneous mixing observed in Fig. 5D (white arrows) likely resulted from the local formation of non-fluorescent labeled linker clusters, as indicated by the lower fluorescence in the microscopy images.

**Figure 5.**
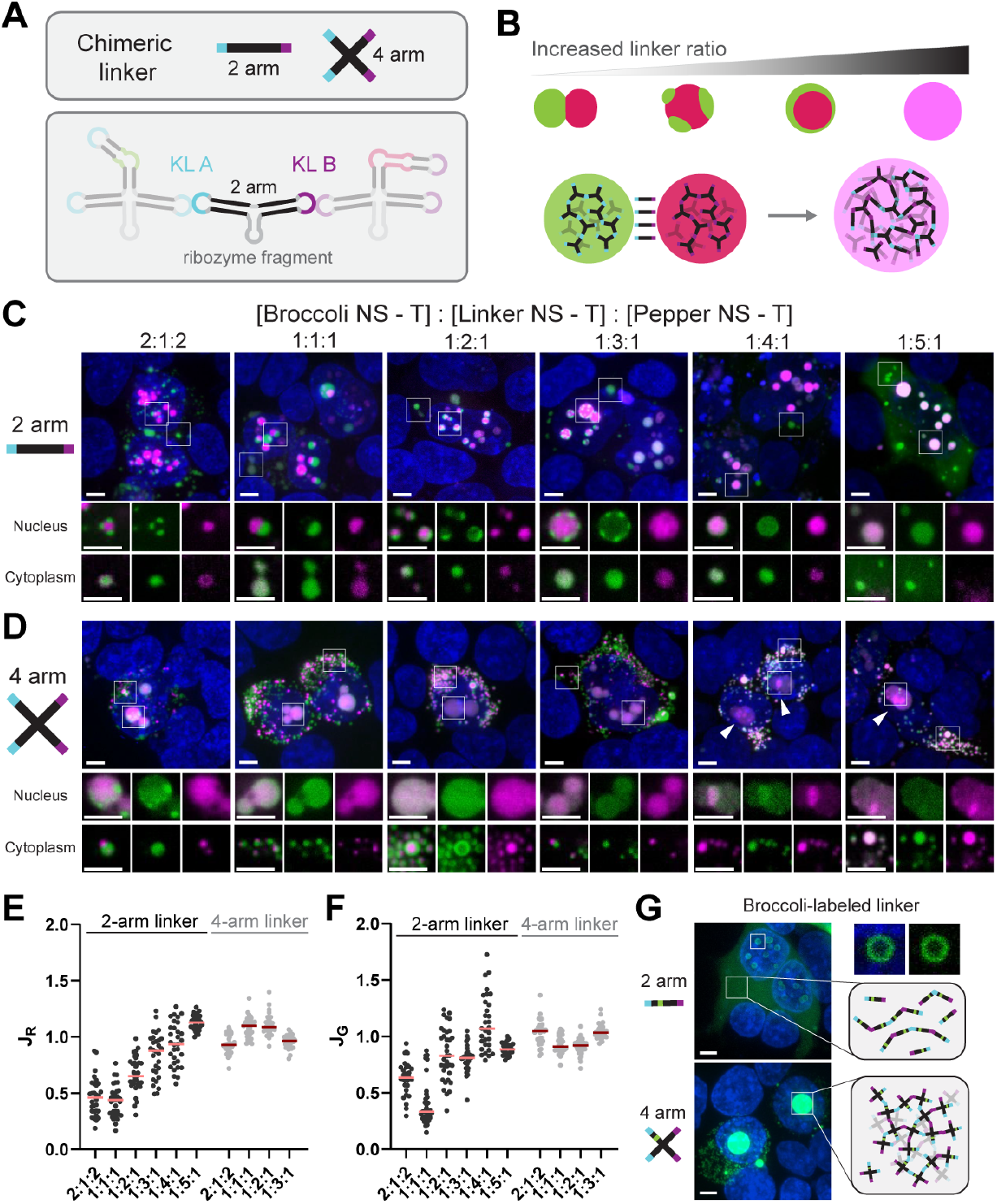
Chimeric linker nanostars at different ratios yield condensates with programmable sub-compartmentalization. **A**, Schematic of two-arm and four-arm chimeric linker nanostars. **B**, Increasing the ratio of chimeric linker over distinct nanostars yields a higher level of mixing. **C, D**, Larger field-of-view (top) and zoomed-in (bottom) confocal micrographs of cells producing nanostars with different ratios between DNA templates ([Broccoli NS - T]:[Linker NS - T]:[Pepper NS - T]) using two-arm (C) or four-arm (D) linkers. **E, F**, For two-arm linkers, mixing indices and increase with the linker ratio. Higher mixing indices for four-arm linkers resulted from doubled valency. **G**, Confocal micrographs show Broccoli-labeled two-arm linkers form no condensate; while four-arm linkers form condensates by themselves. HEK293T cells were stained with Hoechst, DFHBI (for Broccoli), and HBC620 (for Pepper). White arrows indicate non-homogeneous mixing. Images were acquired 48 hours after transfection. Large field-of-view images (top) in panel C are z-projections of stack images captured by a confocal microscope. Zoomed-in view (bottom) on the bottom are single slices from the stack. All experiments were replicated three times. Scale bar, 5 μm.

## Recruitment of target RNAs in trans

One unique advantage of RNA nanostars is their ability to recruit specific target RNA molecules through base-pairing. To demonstrate this, we first built a model RNA target that included a single stem-loop domain with a KL complementary to the nanostar KL. This target was successfully colocalized with the condensates (Fig. 6A), as indicated by high PCC (0.84 and 0.87 for analyzed cells, Fig. 6C). In contrast, we found no colocalization when the KL sequence was missing on either the nanostar or the target RNA (Fig. 6B, Supplementary Fig. 46D and E). While successful, this approach requires modification of the target RNA and thus offers limited flexibility for biological applications. To bypass this constraint, we modified nanostars to carry a dedicated hybridization domain that can include a sequence complementary to the target RNA (Fig. 6D). This is a sequence-specific recruitment strategy applicable in principle to any RNA transcript, provided that the intermolecular hybridization outcompetes local secondary structures. As a proof of concept, we used an arbitrary 21-nt sequence as the recruitment domain. Colocalization was observed only when this domain matched the target RNA sequence (Fig. 6E and F and Supplementary Fig. 46F-H). To quantify colocalization, we provided Manders’ overlap coefficients to quantify the fraction of overlapping independent from signal intensities, as well as PCC values, whose low value is due to the low expression level of target RNA molecules. Although the longer hybridization domain slightly reduced condensate formation likely due to steric hindrance, recruitment efficiency was unaffected, and the effect could be mitigated by increasing the nanostar-to-target RNA ratio (Supplementary Fig. 46C).

**Figure 6.**
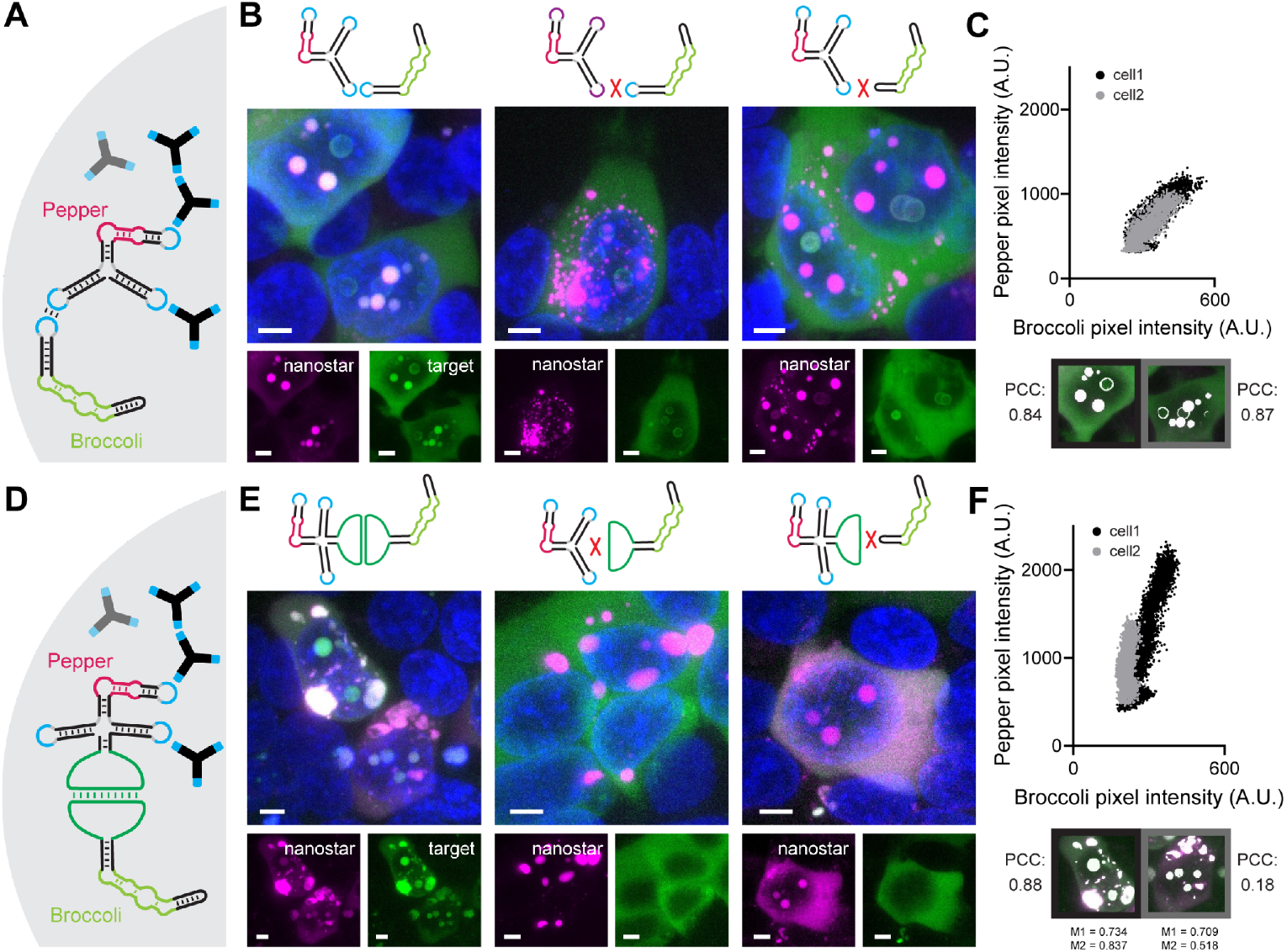
RNA nanostar condensates recruit cellular RNA molecules through sequence-specific interactions. **A**, schematics showing condensates formed by Pepper-labeled RNA nanostars recruit Broccoli-labeled RNA molecules through the addition of the same kissing loop. **B**, confocal micrographs show colocalization occurs only when both the nanostar and target RNA contain matching kissing loop sequences. **C**, scatter plots display pixel intensities in the Broccoli and Pepper channels for each pixel within the white-highlighted regions of interest, along with the corresponding Pearson correlation coefficients. **D**, schematic showing recruitment of target RNA molecules to condensates formed by Pepper-labeled nanostars engineered with a complementary recruitment domain. **E**, confocal micrographs show colocalization occurs only when both the nanostar and target RNA contain matching recruitment domains. **F**, scatter plots showing pixel intensities in the Broccoli and Pepper channels for each pixel within the white-highlighted regions of interest, together with the corresponding Pearson correlation coefficients. Although the signals appear colocalized, the low expression level of the target RNA (green) in cell 2 reduces the signal-to-noise ratio and results in a low PCC. Manders’ overlap coefficients are included here because they quantify only the fraction of overlapping signal. HEK293T Cells were stained with Hoechst, DFHBI (for Broccoli) and HBC620 (for Pepper). PCC scatter plots for control experiments are in Fig. S46. All experiments were replicated three times. All images are z-projections of stack images captured by a confocal microscope. Scale bar, 5 μm.

## Discussion and conclusion

We have demonstrated a simple, yet highly programmable strategy to build RNA condensates in living mammalian cells from short single strands of RNA folding into star-shaped motifs. Leveraging advances in DNA and RNA nanotechnology^35–40^, our work establishes that sequence-specific RNA interactions can act as a structural and functional driver of condensate formation, enabling precise control of condensate material properties and cellular localization through rational sequence design^2,41–46^

A major contribution of this study is the demonstration that synthetic RNA condensates can spatially localize small molecules, proteins, and other RNA with unprecedented granularity. Because most cellular reactions occur in specific subcellular compartments, the ability to program condensate localization offers a powerful means of modulating intracellular pathways. For the first time, we showed that RNA morphology determines condensate distribution independently of any cellular localization signals and allows gradual tuning of the nuclear-to-cytoplasmic ratio through defined design parameters. Another unique feature of RNA-based condensates is their capacity to form discrete, multicomponent compartments, driven solely by rationally designed kissing loop interactions. This capability is difficult to achieve with synthetic protein-based condensates, which typically rely on intrinsically disordered regions that lack the base-pairing programmability of RNA^47^. Together, these properties allow multiple RNA condensates to coexist as distinct cellular compartments that provide simultaneous control over the location, dynamics, and biochemical reactions of recruited biomolecules.

Previously, recruitment of target RNAs into condensates relied on protein–RNA systems such as MCP-MS2, which require engineered recognition tags on the target RNA^48,49^. Here we introduce a more general and programmable approach: direct sequence-specific hybridization between nanostars and endogenous RNA sequences. This strategy eliminates the need for target RNA modification and, in principle, can be applied to any transcript as long as intermolecular hybridization is favorable relative to local secondary structures. Such design flexibility provides a foundation for controlling the subcellular localization of native RNAs and for manipulating biological processes such as rRNA processing and mRNA translation^23,50,51^.

Highly structured circular RNAs are known to participate in diverse cellular processes and interact with native organelles, and we cannot exclude the possibility that their overexpression may contribute to metabolic stress, as suggested by our immunostaining results. Future iterations of this technology could address this through improved expression control, for example by incorporating inducible promoters to fine-tune RNA levels. The emergence of shell-like structures also warrants further investigation. These shells may originate from weak intermolecular interactions, such as G-quadruplex formation within fluorescent aptamers^52^, or from metabolic pathways associated with circular RNAs, consistent with their interaction with G3BP1^26^. Alternatively, they could represent another form of condensate, akin to previously reported anisosomes^27^. Regardless of the underlying mechanism, we view this as an intriguing feature of the system that expands its functional landscape. With the growing recognition of RNA as a central player in cellular phase separation, we anticipate that future versions of this technology will achieve greater selectivity and adaptability, ultimately enabling the construction of synthetic organelles with tailored and novel biological functions.

## Supporting information

Oligonucleotide sequences

## Author contributions

Conceptualization: SL, KP, DLB, EF

Methodology: SL, YK, KW, EPJ, AAT, MVN, DO, MY, DD, AB, WX, MMHL, NYCL, KP, DLB, EF

Visualization: SL, YK, KW, EJP, MVN, DO, AB

Funding acquisition: EF

Supervision: MMHL, NYCL, KP, DLB, EF

Writing – original draft: SL, EF

Writing – review & editing: SL, AB, MMHL, NYCL, KP, DLB, EF

## Acknowledgements

We thank Paul Rothemund and Jongmin Kim for advice and helpful discussions. E.F. acknowledges support from the National Science Foundation (NSF) grant FMRG: Bio award 2134772, by the Alfred Sloan Foundation through awards G-2021-16831 and G-2024-22575, and by the National Institutes of Health through award 1R35GM155833-01.

