## Supplementary materials for "Programmable artificial RNA condensates in mammalian cells"

**Materials and Methods** **2**
1.1 Sequence design 2
1.2 RNA synthesis for in vitro characterization 2
1.3 Plasmid development 3
1.4 Cell culture and maintenance 3
1.5 Transfection 3
1.7 Fluorescence microscopy and live cell imaging 4
1.8 Image processing 5
1.9 Fluorescence recovery after photobleaching (FRAP) 7
1.10 Time-dependent coalescence analysis 8
1.11 Flow cytometry 8
1.12 qPCR 8
1.13 PAGE gel electrophoresis 9
**Supplementary Note 1** **9**
**Supplementary Note 2** **9**
**Supplementary Figures** **10**
**Supplementary Tables** **49**
**References** **51**

#### Materials and Methods

##### 1.1 Sequence design

Nanostars were designed  using NUPACK[2] based on published *in vitro* results[1]. For each design, 10 NUPACK trials were run, the one that generated the lowest defect score was selected. Broccoli, Pepper, Mango and MS2 aptamer sequences were taken from the literature[3–6]. All sequences are listed in a separate Supplementary Dataset 1.

###### 1.1.1 Design of nanostar stem sequences and inclusion of aptamer domains

The stem sequence used in nanostar variant 15nt-3A-Br (Fig. 1 of the manuscript) was adapted from Stewart et al. (Stem 1 design)[1]. We used NUPACK scripts to generate the sequence of 10 and 20 nt-long stems *de novo,* as summarized in Supplementary Note 1. The 25 nt-long stem sequence was adapted from Fabrini et al[7]. Stem sequences were modified to insert Broccoli and Pepper aptamer as part of one of the arms, assuming that nanostar folding assists with correct aptamer folding. We adapted the length of aptamer-including stems to remain comparable with the other nanostar arms. For 10nt-/15nt-/20nt- long arms, we respectively used 4bp+4bp/6bp+6bp/8bp+8bp long domains to flank aptamers. For the 25nt-arm, 8bp+8bp domains were used to flank aptamers due to limitations in DNA synthesis. The sequence of the arm with aptamers was optimized using the NUPACK script in Supplementary Note 2. Because Mango and MS2 aptamers include a functional loop, they cannot be inserted into an arm. For this reason, we included them as an additional arm at the 3' end of their nanostar. We chose the 3' end to prioritize nanostar transcription and folding relative to aptamer folding. Broccoli, Pepper, and Mango aptamers were selected to demonstrate orthogonality because 1) they have non overlapping emission spectra; 2) their fluorescence does not require aptamer dimerization, which would introduce undesired interactions between nanostars. To develop orthogonal nanostars (Fig. 4 and 5 of the manuscript), the stem of 15nt-3A-Br was modified to minimize interactions between nanostars. Using a script similar to the one in Supplementary Note 1, we generated multiple 15nt-long stems. We selected two presenting the lowest defect scores and minimal interaction with the 15nt-3A-Br nanostar, and we modified them to include Pepper and Mango aptamers.

###### 1.1.2 Design of nanostar kissing loop sequences

All kissing loops are 9 nt long and include a 6 nt interaction sequence flanked by 3 unpaired adenine residues, 2 upstream and 1 downstream of the interaction sequence (5'-AA…A-3'). The wild-type kissing loop (5'-GCGCGC) was adapted from the HIV-1 palindromic kissing loop sequence[8]. Orthogonal kissing loops 5'-UCGCGA, 5'-GUCGAC, and 5'-GGUACC were taken from Fabrini et al. [7] Kissing loops 5'-GUAUAC and 5'-UAUAUA were designed by simply replacing GC pairs with AU pairs. Non-palindromic kissing loops were adapted from the 3sβ set designed by Stewart et al[1].

##### 1.2 RNA synthesis for *in vitro* characterization

All RNA strands for *in vitro* experiments were transcribed from custom DNA templates synthesized by Integrated DNA Technologies as LabReady resuspension, standard desalt purification. We annealed non-coding DNA templates with a 21-nt complement including the T7 promoter region and a 4 nt sealing domain (5'-GCGC). These templates were annealed in 1X TE/50 mM NaCl from 90°C to RT at -1°C/min  at 5 µM for storage and used at 0.01 µM during *in vitro* transcription. RNA strands were transcribed in vitro at 37 °C using 7.5% (v/v) T7 polymerase from the AmpliScribe T7-Flash transcription kit (ASF3507, Biosearch Technologies), and transcription buffer prepared in-house: 40 mM of Tris-HCl, 10 mM of NaCl, 30 mM MgCl2, 2 mM spermidine, 7.5 mM each NTP, 10 mM DTT.

#### 1.3 Plasmid development

The highly stable nanostar stem-loop domains facilitate polymerase dissociation. As a result, polymerase chain reactions (PCR) of nanostar DNA templates generate products with incorrect lengths and sequences. For this reason, inserts were directly purchased from Integrated DNA Technologies as two single-stranded, 5’ phosphorylated oligonucleotides containing the sequence of interest, flanked by NotI and SacII restriction sites. The two strands were annealed in 50 mM NaCl and 1x TE buffer using a heat treatment protocol including a 5-minute melt at 90°C, followed by a slow temperature ramp at -1°C/min, and held at 20°C. The resulting products were double-stranded DNA fragments with sticky ends ready for ligation. After annealing, strands were purified with a DNA cleanup kit (NEB T1030). The DNA encoding the nanostar sequences was inserted in the pAV-U6+27-Tornado-Broccoli (Addgene 261587) plasmid. Plasmids were prepared by (1) digestion with NotI-HF (NEB R3189S) (2 µL for 20 µL reactions) at 37°C for 1 hour; (2) purification with the DNA cleanup kit; (3) digestion with SacII (NEB R0157S) (2 µL for 20 µL reactions) at 37°C for 1 hour; and (4) purification with a 0.8% 1x TAE agarose gel to select the product with the correct size. Digested backbones were finally purified using a gel extraction kit (Qiagen 28704). Digested backbone and inserts were ligated at a 1:10 molecular ratio by overnight incubation with T4 DNA ligase (NEB M0202S) at 4°C. Ligated plasmids were transformed into 50 µL DF5Hα competent cells (Thermofisher EC0112 and 18258012) following the manufacturer’s protocol. We then extracted plasmid DNA using a Miniprep kit (Qiagen 27106) following the manufacturer’s protocol. Extracted plasmids were finally sequenced by Eurofins Genomics (whole plasmid sequencing service). The plasmid expressing MCP-mCherry was purchased from Addgene (207668).

#### 1.4 Cell culture and maintenance

HEK293T (ATCC® CRL-3216™), HeLa (ATCC® CCL-2™), and U-2 OS (ATCC® HTB-96™) cells were grown in Dulbecco Modified Eagle’s Medium (DMEM), high glucose, pyruvate (ThermoFisher 11995065) containing 10% Fetal Bovine Serum (FBS) and 100 U/ml Penicillin/Streptomycin (Thermo Fisher) and maintained at 37 °C with 5% CO2 in a humidified incubator. Cells used for imaging were cultured in µ-Slide 8 Well high slides (Ibidi GmbH).

#### 1.5 Transfection

Seeding density was adapted across cell types to achieve ~70% confluence at transfection. Three wells were seeded as replicates for each condition tested. Lipofectamine 2000 (ThermoFisher 11668019) was used for transfecting HEK293T cells. FuGene HD (Promega E2311) was used for transfecting HeLa and U-2 OS cells as it demonstrated less cytotoxicity (additional details below).
For experiments involving the expression of multiple nanostars, the total amount of plasmid DNA used in each experiment was kept constant, with an equal proportion of each nanostar variant. For experiments involving the co-delivery of plasmids expressing nanostars and MCP-mCherry, the total amount of plasmid DNA also remained constant. The plasmids encoding nanostars and MCP-mCherry were mixed and delivered in a 9:1 ratio.

##### 1.5.1 Transfection using Lipofectamine 2000

HEK293T cells were seeded at $5*10^5$ cells/mL, 500 µL/well into 24 well plates one day before transfection. Transfection was performed following the manufacturer's protocol, with a 500 ng final plasmid concentration and a 2 µL final Lipofectamine 2000 volume per well. The DNA-lipid complex was incubated with cells for 4-6 hours and then aspirated and changed to complete DMEM as mentioned above. Cells were reseeded into µ-Slide 8 Well high slides (Ibidi GmbH) the following day at $5*10^5$ cells/mL for imaging. Slides were pre-coated by incubating with 0.001% poly-L-lysine for at least 30 minutes, followed by washing once with PBS before the addition of cells. We incubated reseeded cells overnight and imaged them the next day.

##### 1.5.2 Transfection using FuGene HD

HeLa cells were seeded at $1.5*10^5$/mL; U-2 OS cells were seeded at $4*10^5$/mL, 500 µL/well into 24 well plates one day before transfection. Transfection was performed by mixing 1.65 µg of plasmid with OptiMEM to a total volume of 78 µL. The mixture was vortexed for 1-2 s and centrifuged down. Then we added 4.95 µL Fugene HD to the mixture, vortexed for 1 s, centrifuged down, and incubated at room temperature for 15 minutes. For HeLa and U-2 OS cells, a 15 µL mixture was added to each well. Media change was performed the next day before reseeding. Cells were reseeded into 8-well Ibidi slides following the same protocol as HEK293 cells using the seeding density mentioned at the beginning of this paragraph.

#### 1.6 Total RNA extraction

Cells were transfected in 24 well plates, as described in section 1.6. We changed the media 24 hours after transfection, and collected cells 48 hours after transfection. For collection, we aspirated media, washed with PBS, and trypsinized the cells. After trypsinization, cells were resuspended in PBS, lysed, and RNA was purified using the Monarch® Total RNA Miniprep Kit (NEB T2010S). RNA concentration was estimated using a Nanodrop 2000c by measuring absorption at 260 nm.

#### 1.7 Fluorescence microscopy and live cell imaging

##### 1.7.1 Live cell staining

The culture medium from overnight incubation was aspirated and replaced with fresh medium supplemented with 2 drops of NucBlue Live reagent (Hoechst 33342 nuclear dye, Thermo Fisher R37605) per mL of media, along with the appropriate staining dyes according to experimental conditions. For conditions involving the Broccoli aptamer, we used 40 µM of 3,5-Difluoro-4-hydroxybenzylidene imidazolinone (DFHBI) (Lucerna, 400-5mg); for experiments involving the Pepper aptamer, we supplied 10 nM of (4-((2-hydroxyethyl)(methyl)amino)-benzylidene)-cyanophenylacetonitrile 620 (HBC620) (MedChemExpress, HY-133520). Live cells were then incubated for at least 15 minutes at 37°C before imaging. Cells were imaged in the presence of dyes.

##### 1.7.2 Fixed cell staining and immunostaining

Mouse anti-Coilin (Cajal body colocalization) was purchased from Abcam (ab11822, 1:1900, 1 µg/mL). Mouse anti-SC35 (nuclear speckle colocalization) was purchased from Abcam (ab11826, 1:200, 5 µg/mL). Mouse anti-fibrillarin (nucleolus colocalization) was purchased from Antibodies.com (A85370, 1:200). Mouse anti-G3BP1 (stress granules colocalization) was purchased from Thermo Fisher (66486-1-IG, 1:200, 5 µg/mL). Mouse anti-DCP1A (P body colocalization) was purchased from Novus biological (H00055802-M06, 1:200).
Before fixation, cell culture media were removed and cells were rinsed with PBS (Thermo Fisher 10010023). Cells were then fixed in PBS buffer (Thermo Fisher 14190144) containing 4% paraformaldehyde (Thermo Fisher 043368.9M) for 10 minutes at room temperature, and washed with the PBS buffer three times, each for 5 minutes. Next, we permeabilized cells with 0.5% Triton X-100 (Sigma Aldrich 9002-93-1) in the PBS buffer for 10 minutes and washed them three times. For imaging condensates involving the Mango aptamer, we added PBS supplemented with NucBlue reagent (Thermo Fisher R37605) and 200 nM of TO1-Biotin

(TO1-B) (ABM, G955). Cells were incubated in the buffer for 15 minutes before imaging. For experiments involving immunostaining, cells were further blocked using 3% BSA (w/v, Sigma Aldrich 9048-46-8) in the PBS buffer for 1 hour, and washed three times. Then, cells were stained with corresponding primary antibodies diluted to the above-mentioned concentrations with 3% BSA in the PBS buffer and incubated at 4°C overnight. The next day, primary antibodies were removed and cells were washed three times before the addition of secondary antibodies (Thermo Fisher A-21236, 1:1000 in 3% BSA in PBS buffer). We incubated cells in secondary antibodies for 1 hour before removing the buffer and washing them three times with PBS. For the final wash, PBS was supplemented with 40 μM DFHBI and NucBlue reagent. Cells were incubated in the buffer for 15 minutes before imaging.

##### 1.7.3 Microscopy

FRAP and fusion experiments were performed with epifluorescence imaging using a Nikon Eclipse TI-E inverted microscope and a 60x oil immersion objective. Z-stack confocal images were acquired using a Nikon Ti microscope equipped with an NL5+ camera. Images in Figure 4D and 4K were captured using a Yokogawa CSU X1 spinning disk confocal on an inverted Zeiss stand. Hoechst (NucBlue staining) signals were detected in the UV channel (Ex 405 nm). Broccoli aptamer fluorescence was measured using the GFP channel (Ex 488 nm). Mango aptamer fluorescence was measured using the YFP channel (Ex 514 nm). Pepper aptamer, CY3, and mCherry fluorescence was detected using the RFP channel (Ex 561 nm). Finally, Alexa Fluor™ 647-labeled secondary antibody fluorescence was detected using the 647 nm channel (Ex 647 nm).

#### 1.8 Image processing

Confocal micrographs in the manuscript figures are max-pixel-intensity Z projections, unless otherwise specified in the figure caption. We chose to use Z projections rather than single-plane images because they provide a more comprehensive view of condensate signals across all planes. Midplane images are provided in the Supplementary Figures.

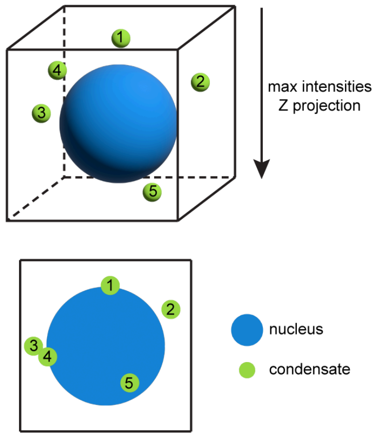


It is well known that Z projections distort the actual volume and localization of objects. For example, condensates in different planes may appear to overlap (as shown for condensates 3 and 4 in the schematic on the right). Condensates located above or below the nucleus, or within the nuclear indentation, may appear to be inside the nucleus itself (as shown for condensates 1 and 5 on the left). These artifacts might cause confusion about the actual cellular localization of condensates. All confocal scans were processed to obtain careful estimates of condensate location and volume, as explained in the following subsections.

##### 1.8.1 Measuring the volume and number of condensates in the cytoplasm and in the nucleus

Cells expressing condensates were cropped into small regions of interest (ROI). At least 50 expressing cells were selected for each nanostar structure. For each ROI, condensates were segmented using Labkit, a machine learning-based plugin for ImageJ, by manually providing labels followed by a training step[9]. The plugin generates masks where condensates in the nucleus and in the cytoplasm are classified into different groups, manifested as distinct pixel intensities. The masks were transformed into Imaris files (Imaris, Oxford Instruments), and reconstructed into 3D surfaces using Imaris (Supplementary Figure S28 and S36). Condensates in the nucleus and in the cytoplasm generated separate surfaces. Any surface object with a volume smaller than 0.015 μm$^3$ (~ two voxels) was considered noise and excluded from our dataset. Adjacent surfaces connected to one object by the software algorithm were manually eliminated. Individual condensates were assigned to different cells manually by comparing reconstruction results and cropped ROIs. Surface volumes were exported for each ROI and pooled into one dataset for each nanostar variant.

##### 1.8.2 Calculating nuclear volume with or without nanostar expression

Nuclear volumes were calculated using the surface tool in Imaris (Oxford Instruments). Before processing, we prepared a cropped ROI when large areas in the field of view lack condensate-expressing cells. Surfaces were generated using the machine learning segmentation tool. A general training algorithm was saved, and the same training classifier was applied to each image. Once machine learning segmentation is implemented, a watershed algorithm of 8 μm is applied to split touching nuclei. We removed surfaces with a high average intensity (likely apoptotic), near the border of the field of view, and volumes smaller than 450 μm (likely segmentation noise). The thresholds for intensity and volume cutoff were adjusted slightly for each image. Manual cleaning of the dataset was done by removing nuclei cut in the z-axis, apoptotic nuclei, and by splitting any touching nuclei. We removed nuclei stacked on top of each other, a frequent event in HEK293T cells. Touching nuclei were removed when Imaris failed to split touching nuclei due to errors in the cut surface. Multiple touching nuclei, or nuclei with a border difficult to distinguish, were removed as well. Cleaned datasets were further processed to identify two classes of nuclei, either expressing or non-expressing condensates. We examined multiple fields of views in multiple samples, gathering data for at least 50 condensate-expressing cells and hundreds of non-expressing cells for each nanostar structure. The volume of each nucleus was normalized by the average volume of all non-expressing nuclei for every sample group.

##### 1.8.3 Partition coefficients for linked condensates

Linked condensates were detected and segmented using FIJI and Python3 scripts adapted from previous work by Fabrini et al[7]. The pipeline included preparing cropped ROI, image enhancement and mask generation, and partition coefficients calculation. ROI were defined from single plane confocal microscopic images with two channels (Em 488 nm and Em 561 nm, corresponding to Broccoli and Pepper channels) acquired with 60x lens and multiple FOVs per sample from three replicates. Cropped ROIs were segmented in FIJI using a macro: for each channel, denoising via a Gaussian Blur (sigma = 1.5), sliding paraboloid background subtraction (smoothing disabled, radius = 10), contrast enhancement (saturated = 0.35), and finally convert to binary masks using the Li and Otsu masks. Because these two segmentation methods can over- or under-segment, the final partition coefficients were calculated as the average of results generated from both methods. Fluorescence images were analyzed using a Python-based pipeline leveraging libraries such as numpy, pandas, and skimage. For each image, green and red fluorescence channels were normalized to a [0,1] scale, and masks were applied to extract mean fluorescence intensities from regions of interest. In Figure 5 of the manuscript we report the quantities $I_G^G$, $I_G^R$, $I_R^R$ and $I_R^G$, where $I_X^Y$ is the average intensity of channel X within the binary mask of channel Y; G stands for the Green channel and R stands for the Red channel. To calculate these quantities we used two approaches, based on the nature of mixing patterns, partial or complete. For 2 arm linkers, ratios ≥ 1:4:1 were considered as complete mixing; for 4 arm linkers, ratios ≥ 1:2:1 were considered as complete mixing due to doubled linker valency. Discussion about the effect of mixing type selection can be found at Supplementary Figure 45. For partial mixing, non-overlapping regions of the masks were identified by subtracting their intersection (nonoverlapping_mask). For complete mixing, the union of the masks was used (union_mask). Mean fluorescence intensities from these regions for both channels were used to compute overlap strengths, represented as J-values, which quantify the ratio of cross-masked to self-masked intensities. $J_R$ was defined as $I_R^G / I_R^R$ and $J_G$ was defined as $I_G^R / I_G^G$. J-values were calculated separately for masks generated by Li and Otsu methods, with the final values averaged between the two.

##### 1.8.4 Partition coefficients for peptides and small molecules

All partition coefficients were calculated using a Python script (see code availability) as:

$$\text{Partition coefficient} = \frac{\text{Fluor. density}_{\text{condensed}}}{\text{Fluor. density}_{\text{dilute}}}$$

Where $\text{Fluor. density}_{\text{condensed}}$ were calculated separately for nuclear and cytoplasmic condensates from z-stacked confocal microscopy images by 1) extracting background fluorescence as the minimal voxel intensity across the field of view, 2) masking condensates region, 3) subtracting background fluorescence for all voxels, 4) summing all non-zero voxel intensities. Then:

$$\text{Fluor. density}_{\text{dilute}} = \frac{\sum \text{background subtracted voxel intensities}}{\text{number of voxels}}.$$

$\text{Fluor. density}_{\text{dilute}}$ were calculated by 1) extracting background fluorescence as the minimal voxel intensity across the field of view, 2) cropping 10*10*10 voxels separately from the dilute phases within the nucleus and cytoplasm, 3) subtracting background fluorescence for all voxels, and finally computing: $\text{Fluor. density}_{\text{dilute}} = \text{mean Fluor. intensity}$

##### 1.8.5 Pearson correlation coefficient (PCC) and Manders' overlap coefficient

All PCC were calculated using ImageJ. Masks of nuclear or cytoplasmic condensates were generated using Labkit and converted to ROIs. Then correlations within the ROI between the two channels were calculated using the plugin Colocalization Finder[11].

As pointed out by Dunn et al.[10], Pearson correlation coefficients "depend upon a simple linear relationship, they will be depressed if measured over a field of cells with heterogeneous expression or uptake of the target molecules, thus under-representing the degree of correlation". For this reason, it is recommended to use a region of interest (ROI) for PCC calculation. In our experiment, we saw distinct linear relationships between cytoplasmic and nuclear condensates due to different preferences of cellular localization due to the Broccoli or Pepper tagging. Nuclear condensates turn to have stronger Broccoli signals, while cytoplasmic condensates turn to have stronger Pepper signals (Figure 4A, bottom). These differences can also be seen on scatter plots shown in Figure 4B, bottom.

In Fig. 5F, cell 2 expressed a low level of target RNA molecule, and the low signal-to-noise ratio resulted in low PCC value although the colocalization is visually apparent. To resolve this, we include Manders' overlap coefficients as they are independent of the intensities of each channel, focusing instead on the proportional overlap of signals where both are present. M1 measures the fraction of the first channel's intensity that overlaps with the second, and M2 measures the fraction of the second channel's intensity that overlaps with the first. The thresholding and Manders coefficient calculation was done by the JaCoP plugin (v2.1.4) on ImageJ.

#### 1.9 Fluorescence recovery after photobleaching (FRAP)

FRAP experiments were performed using a Nikon Eclipse TI-E inverted microscope with a temperature control unit. Temperatures were maintained at 37 °C for all experiments. For *in vitro* experiments, RNA strands were transcribed at 37°C using 7.5% (v/v) T7 polymerase from the

AmpliScribe T7-Flash transcription kit (ASF3507, Biosearch Technologies), and a customized transcription buffer: 40mM Tris-HCl, 10 mM NaCl, 30 mM MgCl2, 2 mM spermidine, 7.5 mM each NTP, 10 mM DTT. We supplied 1% CY3-labeled UTP (ENZ-42505). After three hours, transcription samples were diluted 10 times using our customized transcription buffer, to reduce background fluorescence caused by excess CY3-UTP. Samples for FRAPing Broccoli aptamer were diluted in transcription buffer with DFHBI to a final concentration of 40 μM. *In vitro* samples were loaded into a house-made chamber and sealed with epoxy (Gorilla, 5-minute set) for imaging. For *in vivo* experiments, cells were stained using the protocol described in section 1.7.1.

Condensates were bleached with a 488 nm laser for 200 ms. For *in vitro* samples, imaging was captured once before bleaching and every 5 seconds for 10 min after bleaching. For *in vivo* samples, imaging was captured once before bleaching and every 200 ms for 2 min after bleaching or every 1.5 s for 5 min after bleaching. Images were analyzed by extracting time-dependent average intensities within the bleached area and unbleached area. Normalization was performed to correct photobleaching caused by repetitive imaging using the equation below:

$$\frac{I_{\text{bleach},t}/I_{\text{bleach,max}}}{I_{\text{unbleach},t}/I_{\text{unbleach,max}}}$$

where I denotes the mean pixel intensity in the bleached or unbleached area, t denotes the time point, max denotes the highest pixel intensity within the area among all time points. FRAP plots report the mean ± error bar from N=3 (one region of interest from one replicate was quantified, total three replicates).

#### 1.10 Time-dependent coalescence analysis

For *in vitro* experiments, RNA strands were transcribed, labeled with 1% CY3-UTP, diluted 10 times with transcription buffer, and sealed in a chamber with epoxy, following the same protocol as FRAP experiments. Samples were imaged every 5 minutes for the first 4 hours, then every 20 minutes until 10 hours. We monitored condensate fusion events using a Nikon Eclipse TI-E inverted microscope with a temperature control unit. The temperature was maintained at 37°C for all experiments.

For *in vivo* fusion experiments, we imaged cells (in media supplemented with 40 μM DFHBI and 2 drops/mL NucBlue) every 5 minutes for 60 minutes under the confocal microscope. The temperature was maintained at 37°C. Fusion events were identified manually. Data processing was performed using a script in Python3 described in our previous work[1]. To summarize, each fusion event was identified manually and segmented using Otsu thresholding. The binary mask was then labeled to extract the centroid position, major and minor axis lengths, and orientation. Best-fit-ellipses were generated based on the extracted data. The aspect ratio was calculated as the major-to-minor axes ratio and used for curve fitting. The time constant $\tau$ was calculated by fitting the Aspect Ratio vs time profiles with an exponential decay with the formula $1 + Ae^{(-t/\tau)}$.

#### 1.11 Flow cytometry

Flow cytometry experiments were performed using a BD FACSAria flow cytometer. Cells were transfected in 24 well plates, as described in section 1.6. We changed the media 24 hours after transfection, and collected cells 48 hours after transfection. For collection, we aspirated media, washed with PBS, and trypsinized the cells. After trypsinization, cells were resuspended in PBS supplemented with 10% FBS and dyes and filtered through the 40μm cell strainer (Fisher

Scientific Cat.# 22363547) for flow cytometry. Hoechst was detected using a laser with Ex 405 nm and a 450/50 nm filter; DFHBI was detected using a laser with Ex 488 nm and a 530/30 nm filter.

##### 1.12 qPCR

Reverse transcription was carried out with equal amounts of RNA using the Protoscript II First Strand cDNA Synthesis Kit and random hexamers (New England Biolabs). RT-qPCR was then performed using ten-fold diluted cDNA and the Luna Universal qPCR Master Mix (New England Biolabs) in the CFX Real-Time PCR system (Bio-Rad), courtesy of the UCLA Virology Core. qPCR conditions used as previously described[12,13]. Target transcript levels were determined by normalizing the CT (cycle threshold) value of the target transcript to that of the housekeeping gene RPS11 transcript. Fold change was calculated using this normalized value relative to lipofectamine control expression levels. For RT-qPCR primers, see Supplemental Table 4.

##### 1.13 PAGE gel electrophoresis

Gel premix was prepared by adding 42 g of urea to nanopure water, the mixture was then heated until the urea completely dissolved. This mixture was allowed to cool to room temperature, and then a 40% (v/v) 19:1 acrylamide/bis-acrylamide solution was added in the appropriate volume for the desired percentage (final volume 100 mL). To start polymerization, 8 mL of pre-mix was added in appropriate ratios with TBE and nanopure water, ammonium persulfate (APS), and tetramethylethylenediamine (TEMED). Gels were cast in 8 × 8 cm, 1 mm thick disposable mini gel cassettes (Thermo Scientific, #NC2010) and allowed to polymerize for 30 minutes before electrophoresis. After curing, the gel was pre-run in a 1X TBE buffer for 30 minutes. Wells were washed carefully to remove excessive urea. Samples and low-range ssRNA ladder (NEB, N0364S) were prepared by mixing individual strands with denaturing RNA loading dye (NEB, B0363S), then heated at 70 °C for 10 minutes and immediately placed on ice. Due to the low expression of exogenous RNA in mammalian cells, 5 µg of total RNA extraction was loaded onto each well. Gels were run at room temperature at 100 V in 1X TBE unless otherwise noted. After electrophoresis, the gels were washed three times, each for 5 minutes, with nanopure water, then stained with DFHBI-1T staining buffer (10µM DFHBI-1T, 40mM HEPES, 100mM KCl, and 1mM $MgCl_2$) for 15 minutes. After staining, gels were imaged using the Bio-Rad Gel Imaging Systems. Then, gels were washed three times, each for 5 minutes again to remove the DFHBI-1T and stained in 1xSYBR Gold Nucleic Acid Gel Stain for 15 minutes and imaged again.

#### Supplementary Note 1

NUPACK script for 20nt-3WT stem sequence design. A similar script was used for the 10 nt long stem designs.

```
--------------------------------------------------------------------
material = rna1999
temperature = 37
trials = 10

structure NS= U1 D20 U9 U2 D20 U9 U2 D20 U9
```

domain arm1= N20AAGCGCGCAN20AA
domain arm2= N20AAGCGCGCAN20AA
domain arm3= N20AAGCGCGCAN20

NS.seq = arm1 arm2 arm3

prevent = AAAA, CCCC, GGGG, UUUU, KKKKKK, MMMMMM, RRRRRR, SSSSSS, WWWWWW, YYYYYY

#### Supplementary Note 2

NUPACK Script for 15nt-3A-Br stem sequence optimization

—--------------------------------------------------------

material = rna1999
temperature = 37
trials = 10

### DU+ notation
structure NS= U41 D6 (U12 D6 (U9) U17) U2 U39

### Broccoli aptamer sequence underlined
domain arm1= GCGAGAGCGCUGCCCAAUCGCGAAGGGCAGCGCUCUCGCAA
domain arm2= N6ACGGUCGGGUCCN6AAUCGCGAAN6GUCGAGUAGAGUGUGGGN6AA
domain arm3= GCGUUCACACUGACCAAUCGCGAAGGUCAGUGUGAACGC

NS.seq = arm1 arm2 arm3

prevent = AAAA, CCCC, GGGG, UUUU, KKKKKK, MMMMMM, RRRRRR, SSSSSS, WWWWWW, YYYYYY

#### Supplementary Figures

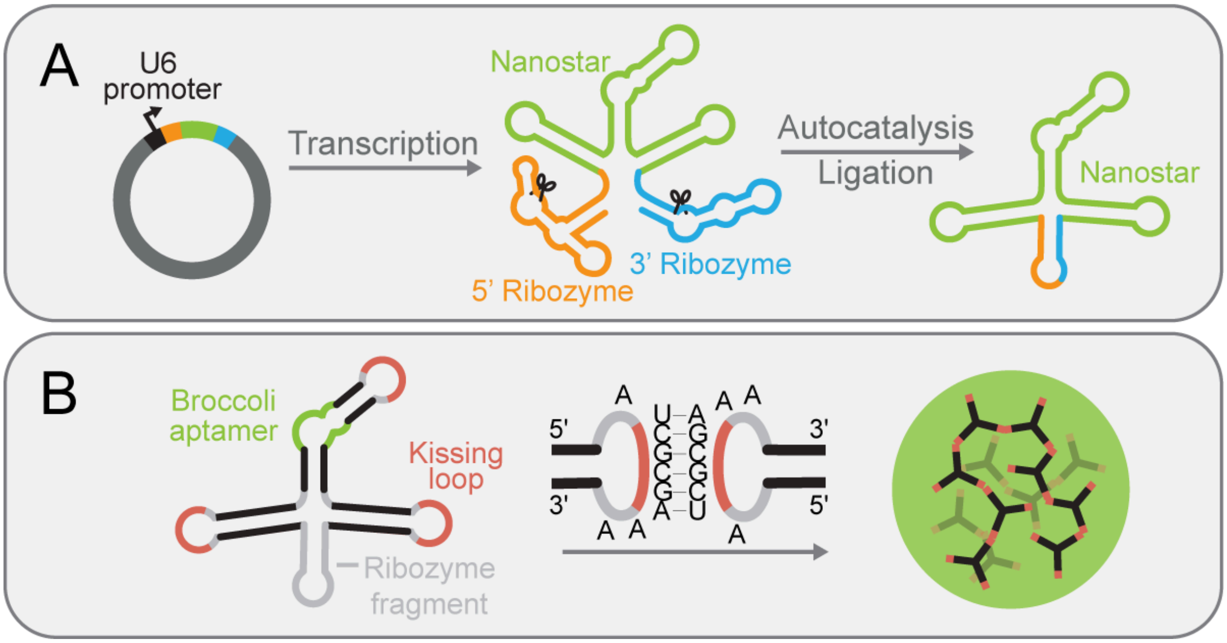


**Figure S1. Autocatalytic circularization of RNA nanostar and condensate formation.**
**A,** We adopted an expression system in which each nanostar sequence (in green) is flanked by 5'- (in orange) and 3'- (in blue) self-cleaving ribozymes designed by Litke et al[14]. Once transcribed, ribozymes autocatalyse and generate functional groups on the new RNA ends. The RNA molecule then becomes a substrate for an endogenous RNA ligase, and becomes circularized before ligation. **B,** After endogenous circularization an RNA nanostar includes three stems (in black), three identical and self-complementary kissing loops (in orange), a ribozyme fragment (in gray, at the bottom), and other adenine spacers (in gray). Phase separation is induced by inter-molecular hybridization between kissing loops.

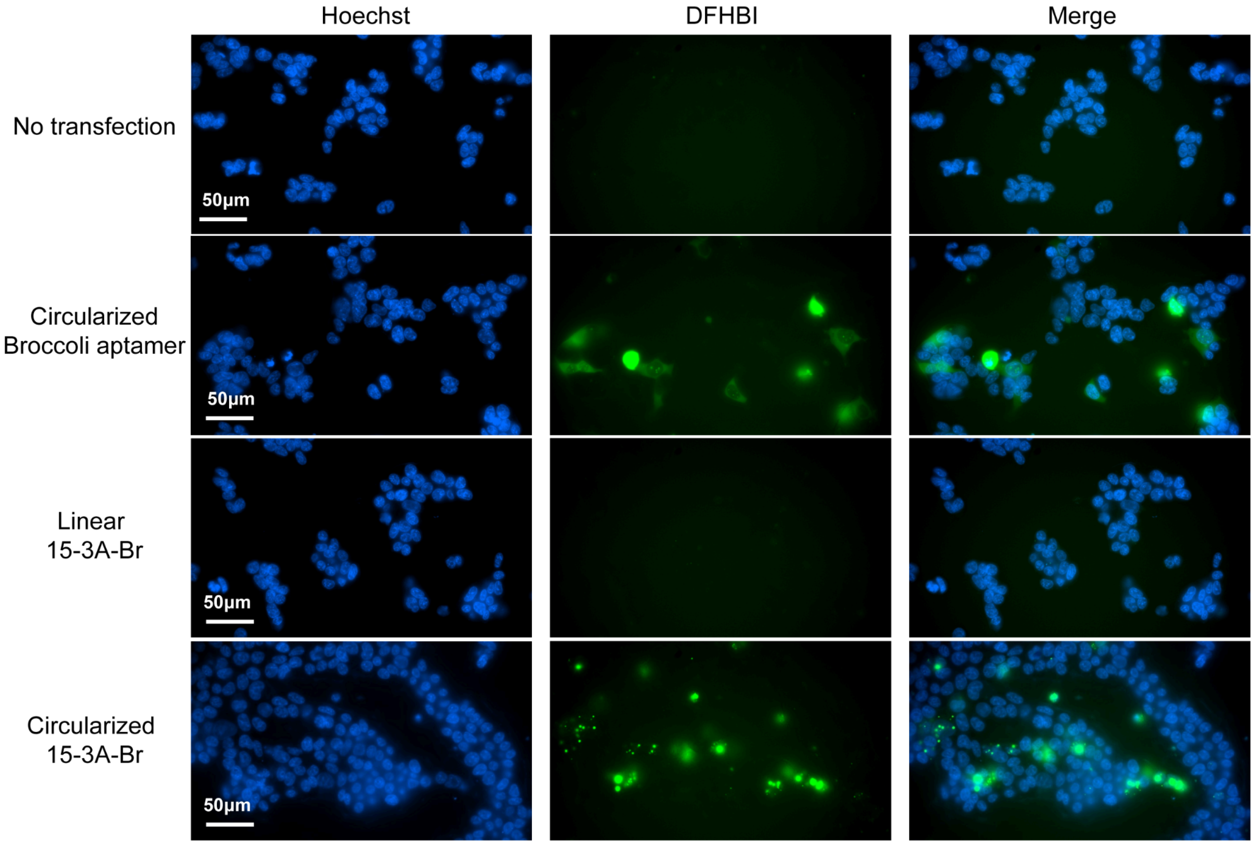


**Figure S2. Circularization significantly enhances RNA accumulation.** Representative images showing fluorescence from HEK293T cells with no transfection, transfected with plasmids expressing circular Broccoli, linear, or circularized nanostars presenting three 15 nucleotide arms carrying the Broccoli aptamer (15nt-3A-Br). Linear 15nt-3A-Br was expressed using the same vector, but without the 3'- and 5'- ribozyme sequence. Without circularization, fluorescence signals were barely detectable. Images are representative of three replicates taken 48 hours after transfection. All images were taken using the same exposure time. Scale bar, 50 µm.

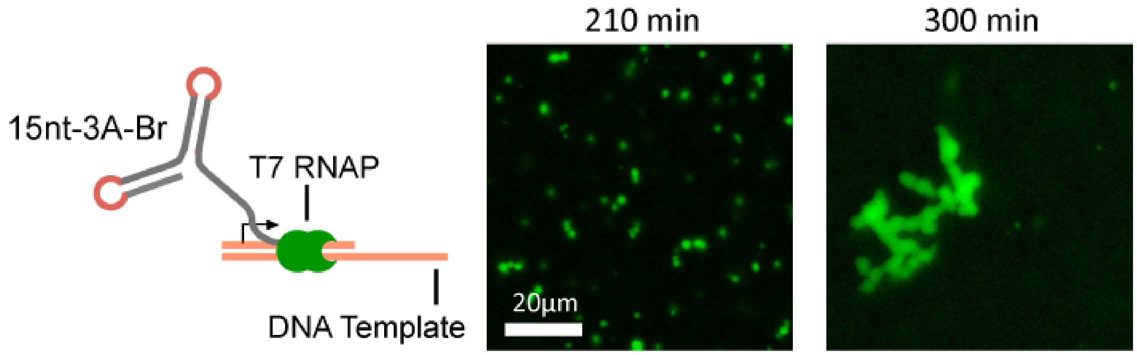


**Figure S3. Cotranscriptional formation and growth of 15nt-3A-Br condensates in vitro.** Condensate formation during *in vitro* transcription. DNA templates are partially annealed to have a double-stranded promoter. RNA strands were transcribed in vitro at 37°C using 7.5% (v/v) T7 polymerase and transcription buffer and stained by 40 µM DFHBI. Images are representative of three replicates.

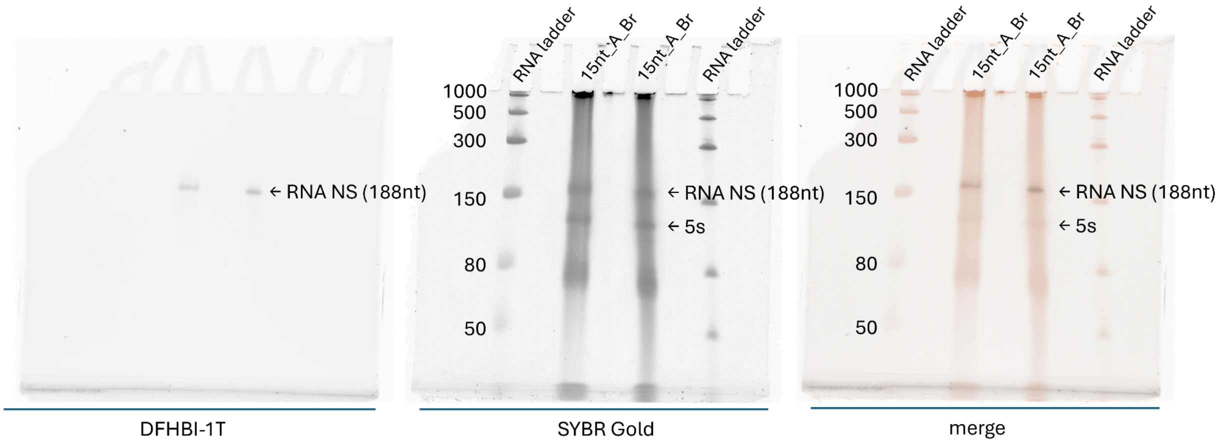


**Figure S4. In-gel staining of Broccoli-tagged RNA nanostar in HEK293T cells.** HEK293T cells were transfected with a plasmid encoding 15nt-3A-Br. After 48 hours of expression, total RNA was isolated, separated by denaturing PAGE, and stained with DFHBI-1T and 1x SYBR Gold. RNA nanostars tagged with Broccoli aptamer are clearly detectable. Circularized RNA nanostars migrated faster than ssRNA ladder strands of the same length, consistent with previous observations[14].

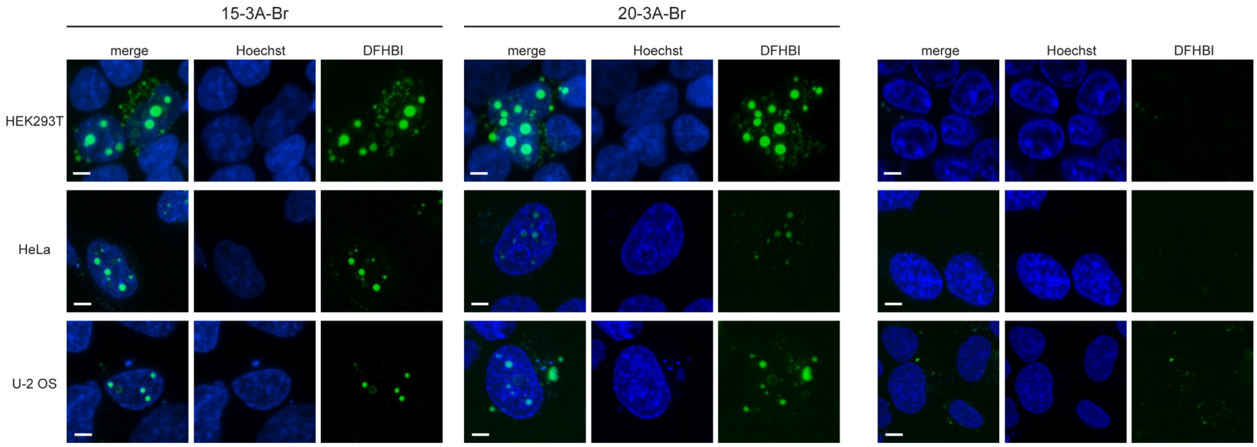


**Figure S5. RNA nanostars form condensates in HEK293T, HeLa, and U-2 OS cells.** Cells are stained with NucBlue reagent and 40μM DFHBI. Expression level is higher in HEK293T cells as the presence of T antigen facilitates plasmid replication. We tested two variants of our nanostar design: one with 15-nucleotide-long arms (15nt-3A-Br, left) and another with 20-nucleotide-long arms (20-3A-Br, middle), compared to non-transfected cells (right). Images are representative of three replicates. Scale bar, 5 μm.

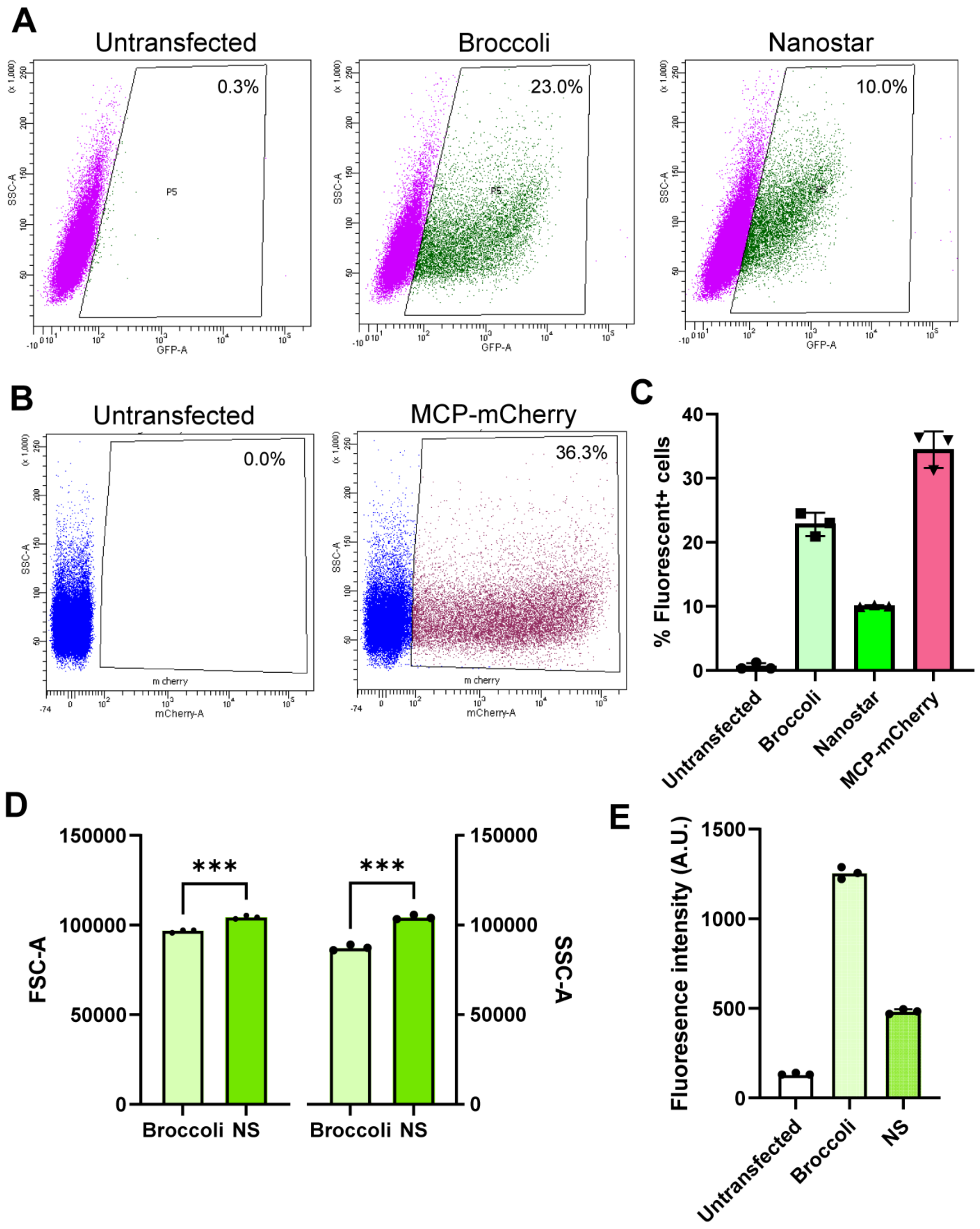


**Figure S6. Flow cytometry plots and quantification comparing untransfected cells with cells expressing circularized Broccoli aptamer, circularized Broccoli-tagged RNA nanostars, or MCP-mCherry.** In general, cells expressing circularized RNA showed lower transfection efficiency than those expressing mCherry, as shown in B and C. This may also reflect the detection limit of the flow cytometer, since Broccoli fluorescence is much weaker than mCherry. Among circularized RNA constructs, cells expressing nanostars showed lower transfection efficiency (C), larger size, higher granularity (D), and lower fluorescence intensity (E) compared to those expressing Broccoli alone. Cells were analyzed 24 hours post-transfection. SSC-A, side scatter amplitude; FSC-A, forward scatter amplitude. Significance was determined by unpaired t-test; ***$p < 0.001$.

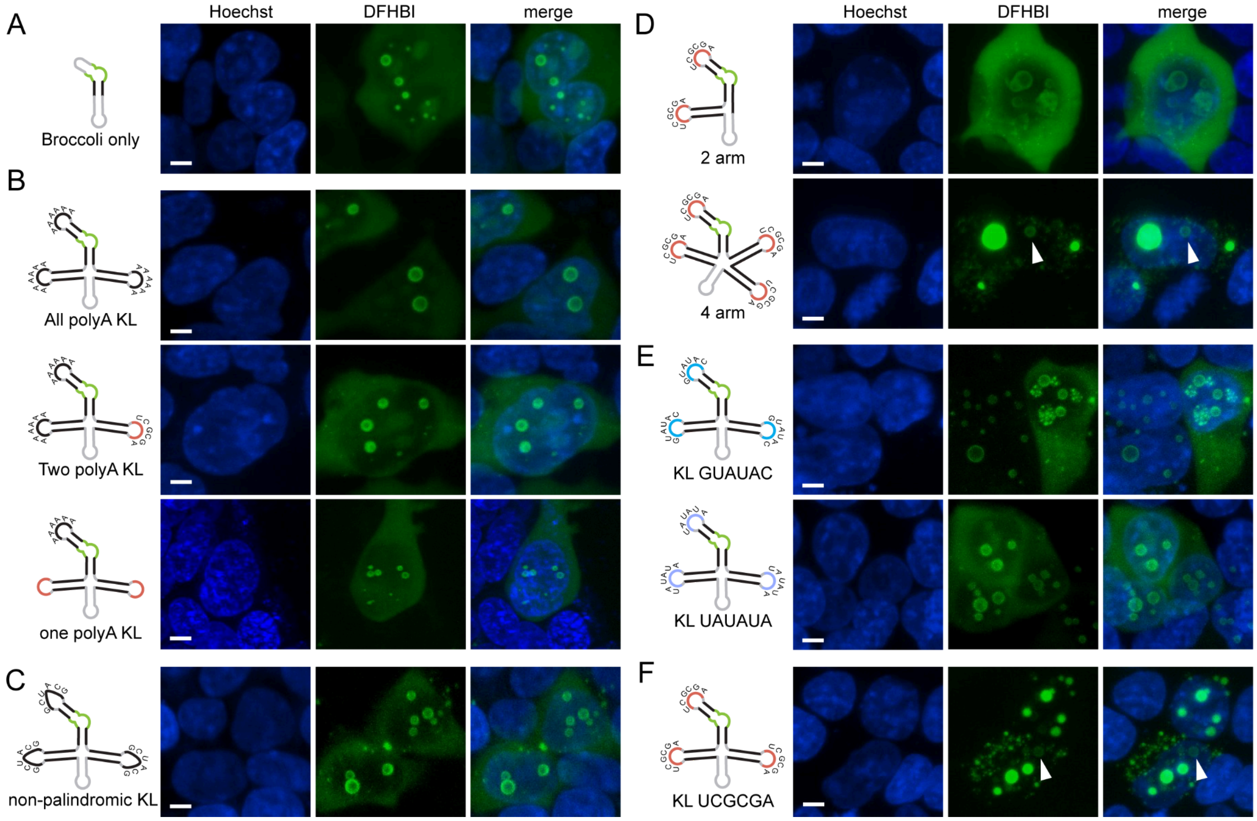


**Figure S7. Shells emerge in cells expressing circularized RNA and a variety of control nanostars, regardless of their capability to form condensates. A,** Expression of circularized Broccoli yield diffused fluorescence in the cytoplasm and formed shells in the nucleus. **B,** Replacing kissing loops with polyadenine domains disrupted condensate formation and generated shells. Changing one kissing loop was sufficient to disrupt condensate formation. **C,** Expression of nanostar with a non-palindromic kissing loop formed shells in the nucleus. **D,** Eliminating one arm from the nanostar formed exclusively nuclear shells (top). Addition of one arm formed big, bright condensates and shells in the nucleus and small puncta in the cytoplasm. **E,** Motifs designed to have weak nanostar-nanostar interactions formed abundant nuclear shells. **F,** shells formed in the nucleus of cells expressing nanostar 15nt-3A-Br. Cells were stained with NucBlue reagents and 40 μM DFHBI. Images are all z projections, and are representative of three replicates. Scale bar, 5 μm.

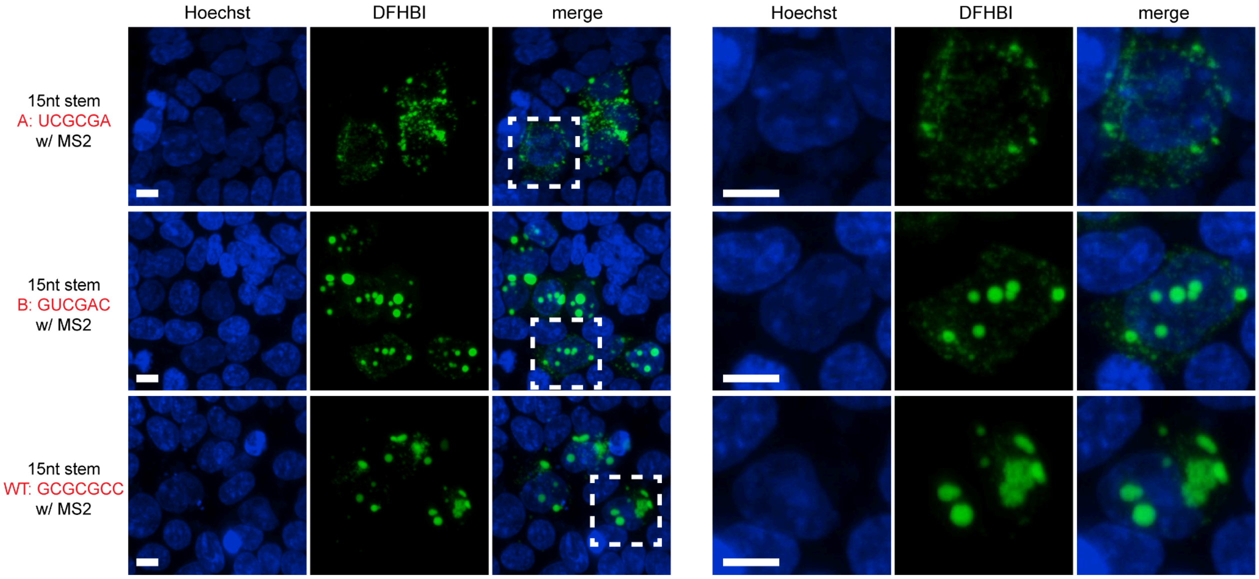


**Figure S8. Nanostars including the MS2 aptamer form condensates when adopting different kissing loops with varying strength.** Representative images (left) and zoomed-in views (right) of the white-squared area showing cells transfected with nanostars that include an MS2 domain and differ by kissing loop sequence (A/B/WT). The strength of kissing loop interactions increases as GC content increases from top to bottom, resulting in more nuclear retention and more aggregate-like condensates in the cytoplasm. Cells were stained with NucBlue reagents and 40 µM DFHBI. Images are representative of three replicates. Scale bar, 10 µm.

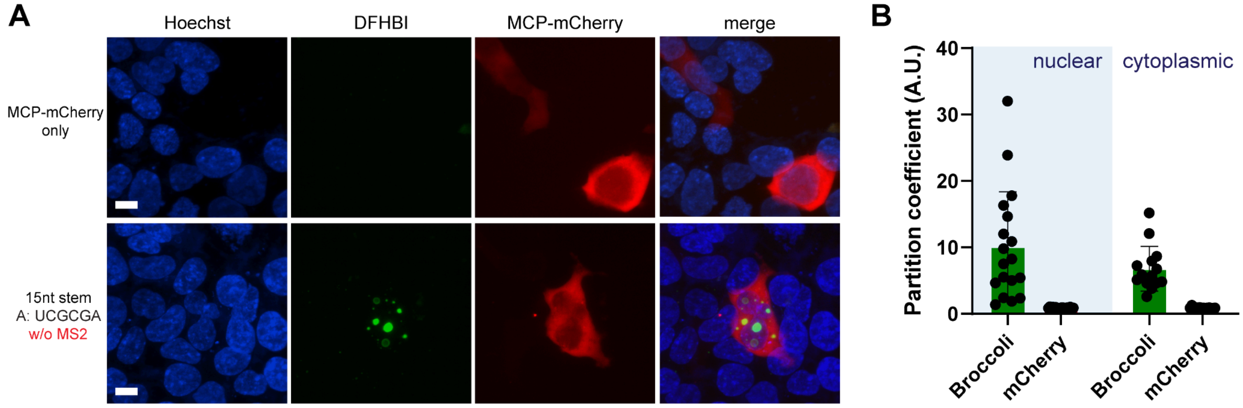


**Figure S9. RNA nanostars lacking the MS2 domain do not recruit mCherry. A,** Representative images showing transfection of plasmids expressing MCP-mCherry (top) and co-transfection plasmids expressing MCP-mCherry and nanostars (bottom). MCP-mCherry expression in cells leads to a homogenous distribution of fluorescent signals across cells. Protein concentration in the cytoplasm is higher than in the nucleus. Without MS2 aptamer, nanostars form condensates that do not colocalize with MCP-mCherry. **B,** Partition coefficients of cells co-expressing nanostars that lack MS2 aptamer, in the presence of MCP-mCherry. Broccoli-tagged nanostars demonstrating slightly higher nuclear partition coefficients than the cytoplasmic ones. Partition coefficients of mCherry for both nuclear or cytoplasmic condensates averaged around 1, indicating no recruitment. Images are representative of three replicates. Scale bar, 10 µm.

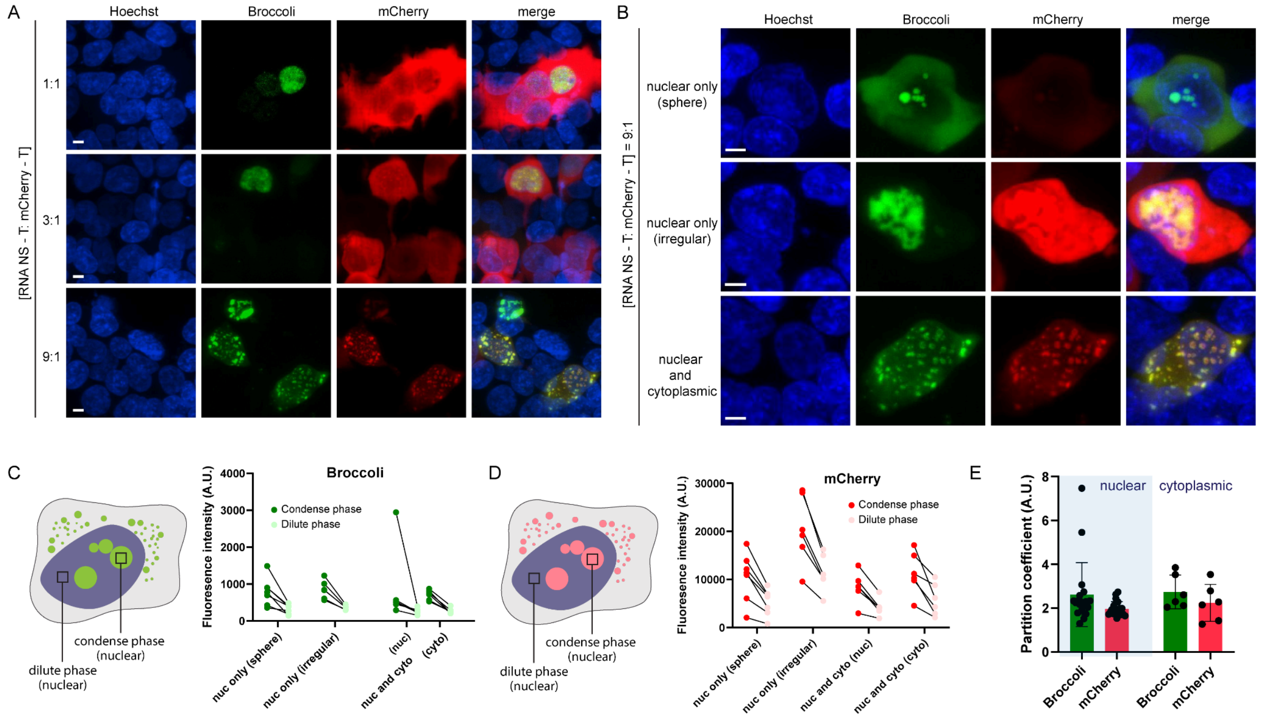


**Figure S10. Condensate morphology is influenced by both the overall expression levels of mCherry and RNA and their relative expression. A,** Example confocal images of cells co-transfected with mCherry and RNA nanostar plasmids at different transfection ratios (mCherry:RNA = 1:1, 1:3, 1:9). Excess mCherry binding to nanostars causes irregularly shaped aggregates, while higher RNA ratios promote spherical condensates. **B,** Example images at 1:9 transfection ratio, normalized to the same brightness range (brighter indicates higher expression), confirm that excessive mCherry expression correlates with irregular aggregate formation. **C,** Quantification of mean fluorescence intensity reveals that RNA (Broccoli) expression remains constant across spherical and irregularly shaped condensates, whereas **D,** mCherry levels increase in irregularly shaped aggregates, both in the dilute and condensed phases. In cells exhibiting both cytoplasmic and nuclear condensates, mCherry localizes primarily to the cytoplasm (lacking an NLS signal), and results in slightly higher fluorescence intensity in both dilute and condensed phases when compared to the nucleus. **E,** Partition coefficient analysis shows consistent values (~2) for RNA and mCherry across condensates in the nucleus and in the cytoplasm. The Broccoli partition coefficient is 3-4 fold lower when compared to RNA condensates that do not recruit mCherry (Fig. S9), indicating that the presence of cargo molecules has a significant impact on the composition of the dense phase.

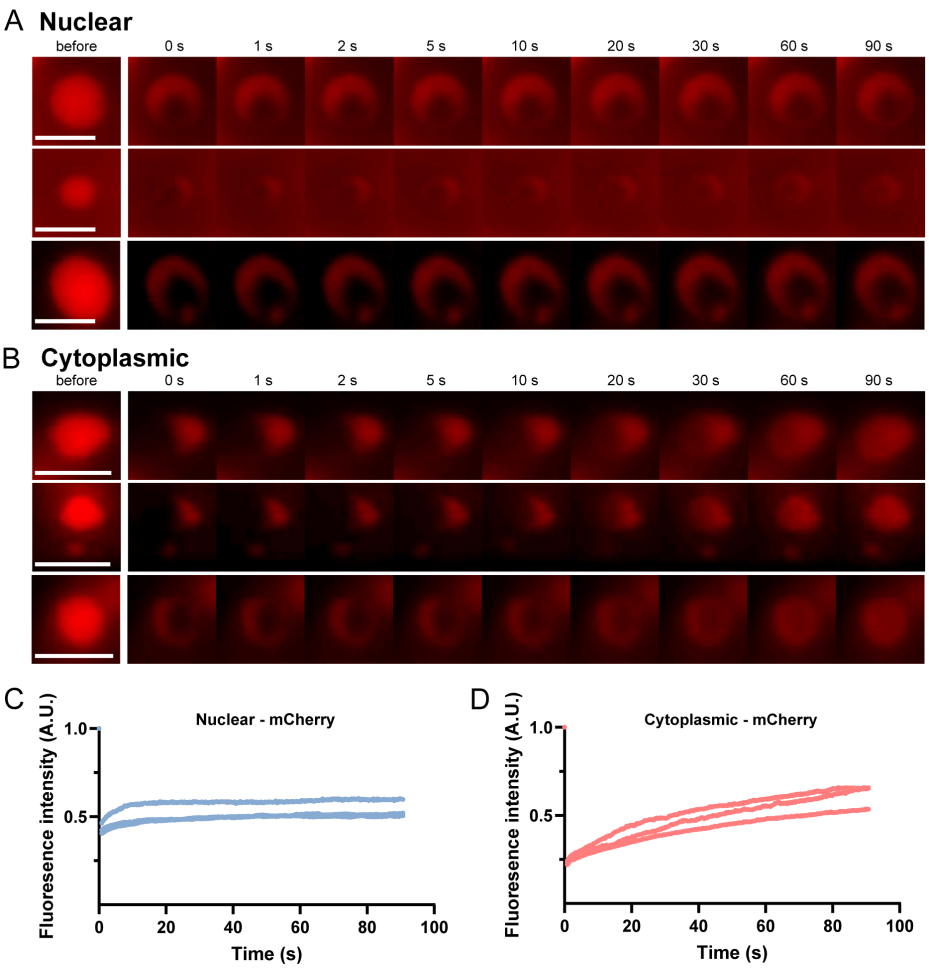


**Figure S11. FRAP of MCP-mCherry recruited to nuclear and cytoplasmic condensates formed by MS2-modified nanostars shown in Figure 1E.** Representative images (A and B) and plots (C and D) of normalized fluorescence intensity showing FRAP behavior of MCP-mCherry cargo recruited through the MS2 domain on nanostars. Scale bar, 5 μm.

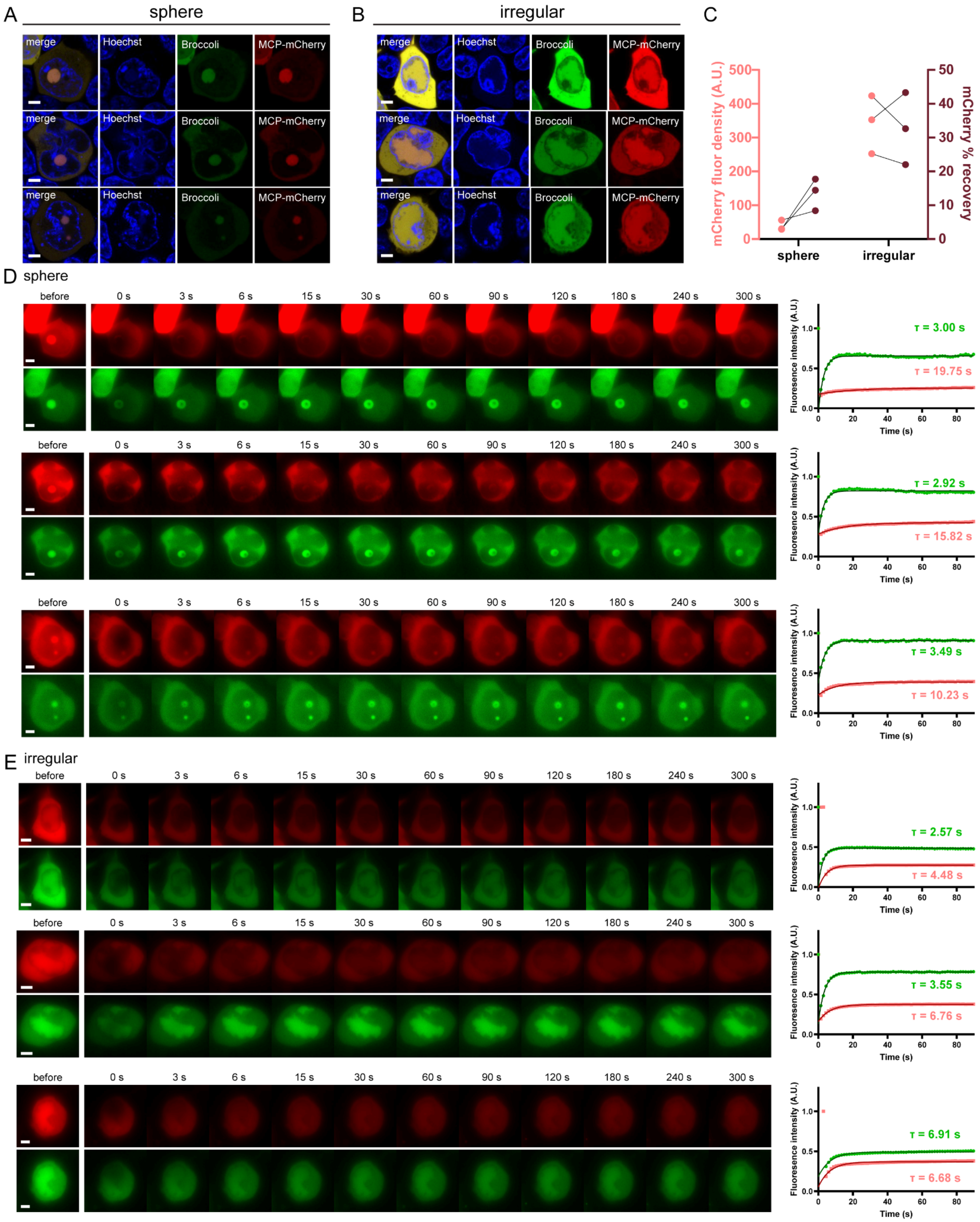


**Figure S12. Irregularly shaped aggregates show faster and greater mCherry recovery after photobleaching compared to spherical condensates. A, B,** Split-channel confocal images of cells forming spherical condensates (A) or irregularly shaped aggregates (B) (more discussion of the two morphologies can be found in Fig. S10B-D). All confocal images were normalized to the same visualization range (brighter indicates higher expression). As shown in Figure S10, irregular aggregates exhibit excessive mCherry expression relative to spherical condensates. **C,** Correlation between mCherry fluorescence density within condensates and percentage of mCherry recovery after photobleaching shown in D and E. Higher mCherry fluorescence density corresponds to a higher percentage of mCherry recovery. **D, E,** Dual-channel FRAP results for spherical condensates (D) and irregularly shaped aggregates (E). Quantification shows that while Broccoli recovery remains similar, mCherry recovery is consistently faster in irregular aggregates. This is likely because mCherry, as a bulky cargo that does not directly participate in condensation, loosens the RNA condensate network and increases its diffusivity. Dots represent mean fluorescence intensity at each time point, and lines represent fitted recovery curves $Y = Y_0 + (Y_\infty - Y_0) * (1 - e^{-\frac{t}{\tau}})$. Due to cellular movement, condensates drift along the z-axis during long observation periods, leading to apparent fluorescence increases after the plateau and affecting fitting parameters. To ensure consistency, curve fitting was performed using the first 90 s of data. Scale bar, 5 μm.

Scale bar: 5um

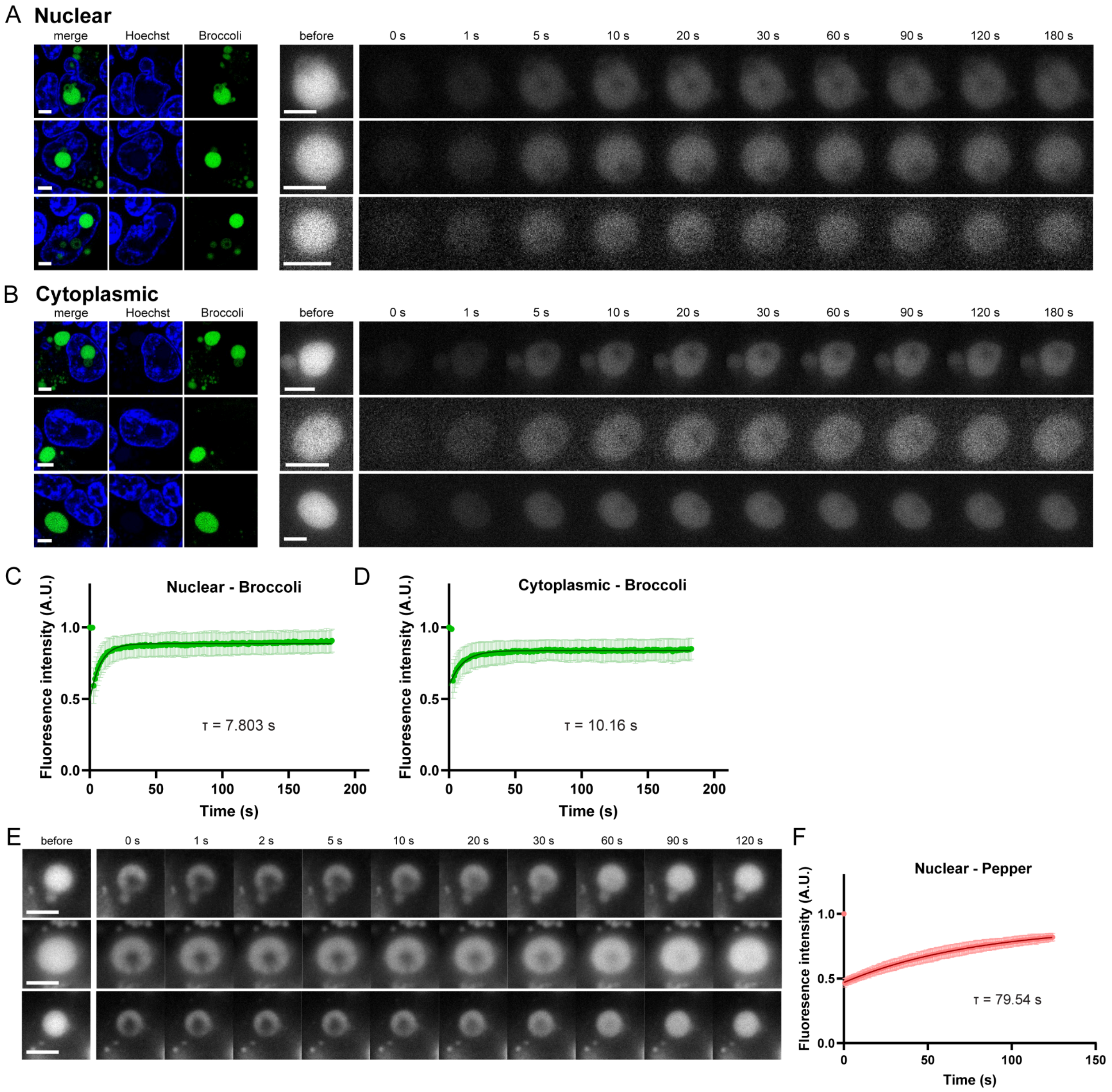


**Figure S13. FRAP of nuclear condensates formed by nanostars labeled with Broccoli (15nt-3A-Br) and Pepper (15nt-3B-Pp) aptamers. A, B,** Condensates' cellular location were confirmed by mid plane confocal images (left). Representative images (right) showing FRAP behavior of 15nt-3A-Broccoli. **C, D,** Dots indicate mean intensity at the corresponding time point. Error bars indicate standard error. Line indicates the fitted curve from equation $Y = Y_0 + (Y_\infty - Y_0) * (1 - e^{-\frac{t}{\tau}})$. **E, F,** representative images (E) and quantification (F) of 15-3A-Pepper. Scale bar, 5 μm.

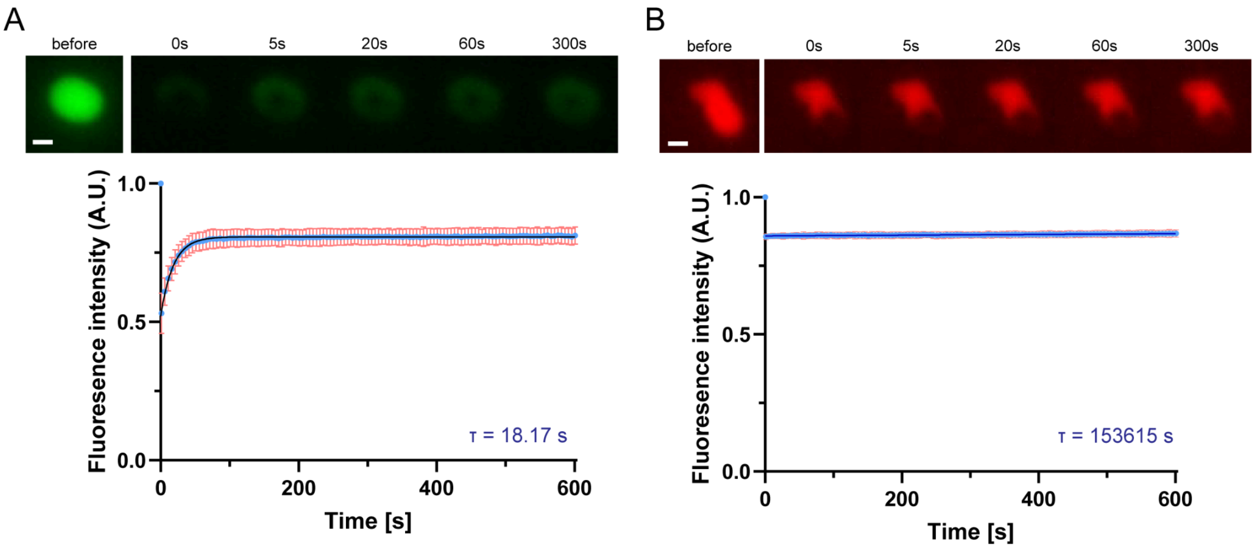


**Figure S14. FRAP of condensates (15nt-3A-Br nanostar) produced during *in vitro* transcription targeting Broccoli aptamer or Cy3.** Representative images (top) and plots of normalized fluorescence intensity showing FRAP behavior of Broccoli aptamer (**A**) and CY3-labeled RNA (**B**). The *in vitro* transcription reaction was supplied with 1% CY3-labeled UTP, and diluted 10x using 1x transcription buffer and DFHBI (final concentration 40 µM) before imaging to reduce strong background fluorescence caused by free Cy3-UTP. No recovery in the Cy3 channel might be attributed to the dilution, leaving little to no Cy3-labeled RNA in the dilute phase available for recovery. Images are captured in GFP channels for A and RFP channels for B. Blue dots indicate mean intensity at the corresponding time point. Orange error bars indicate standard error. The dark blue line indicates the fitted equation $Y = Y_0 + (Y_\infty - Y0) * (1 - e^{-\frac{t}{\tau}})$. The mean and standard error were calculated from 3 fields of view, each belonging to one replicate. Scale bar, 1 µm.

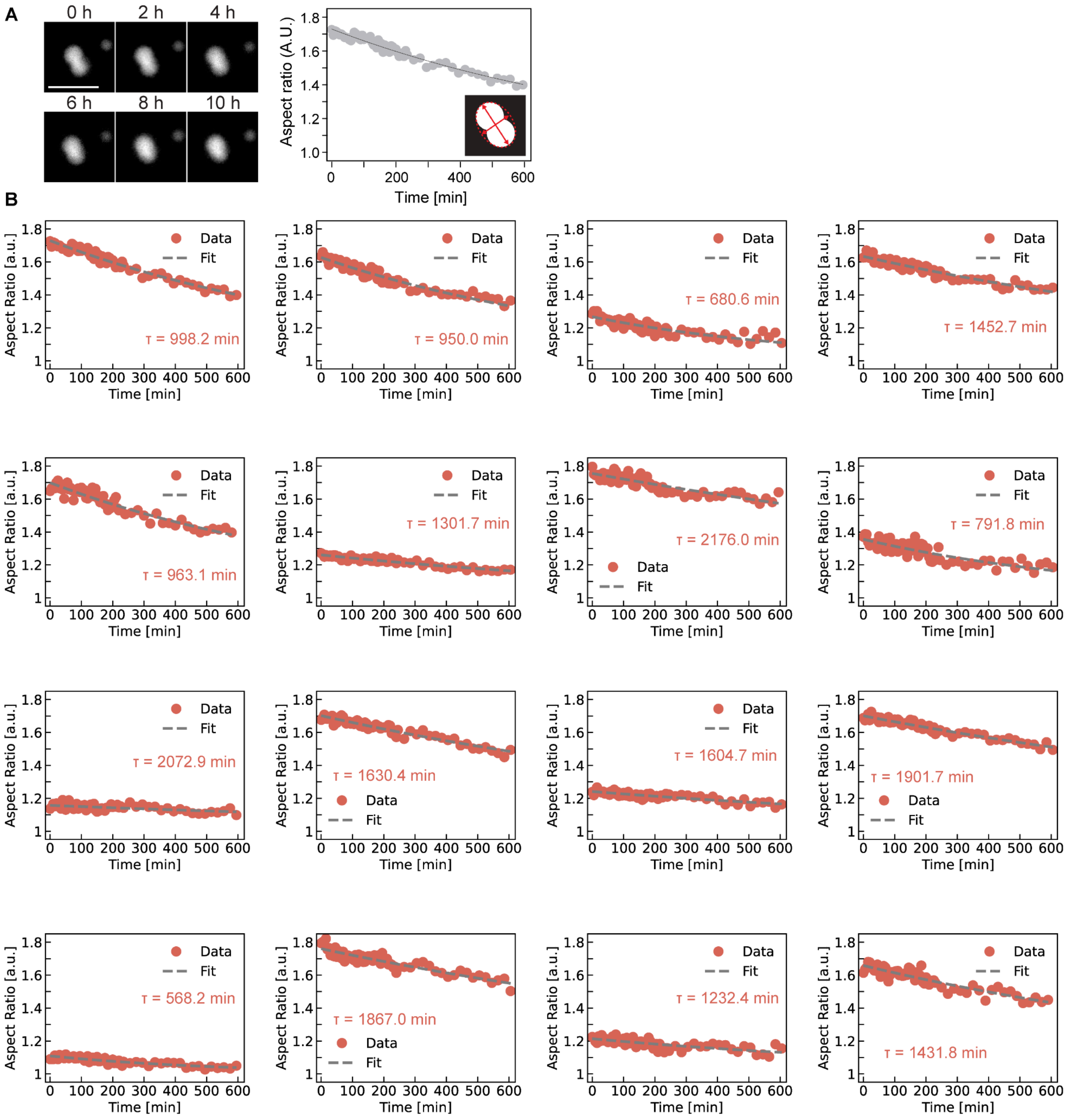


**Figure S15. Time-dependent coalescence of condensates produced during *in vitro* transcription of nanostar 15nt-3A-Br. A,** Example micrographs of one fusion event. Schematic shows the definition of aspect ratio as the major and minor axes ratio from the best-fit-ellipses. **B,** Analysis results tracking 16 condensate fusion events measured within one field of view from one sample. Nanostars were transcribed using 1% CY3-UTP for labeling; condensates were monitored in a sealed chamber, and imaged on a heat stage at 37°C. The dashed line is a fit of the exponential function $1 + Ae^{(-t/\tau)}$. Scale bar, 5 μm.

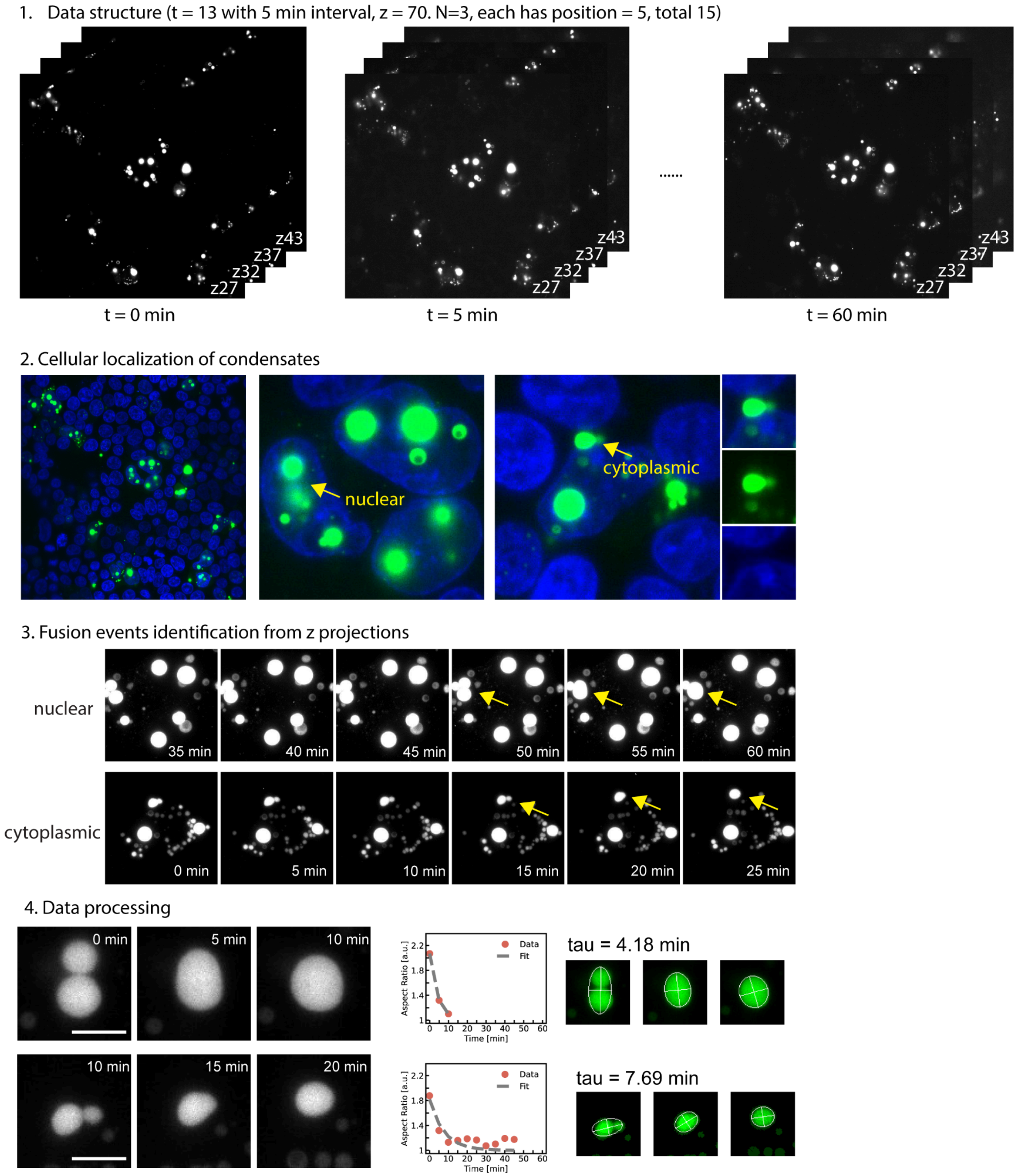


**Figure S16. In vivo fusion data processing workflow.** Cells were transfected to express 15nt-3A-Broccoli for 48 hours, and stained with DFHBI for Broccoli (green) and Hoechst for the nuclei (blue) before imaging. 1) Time-lapse images were collected using confocal microscopy as z-stacks every five minutes, across five positions per replicate and three biological replicates. 2) A single mid-plane image was selected to determine the cellular localization of condensates while minimizing phototoxicity. 3) Fusion events were manually identified. 4) A Python script was used to extract aspect ratios. Scale bar, 5 μm.

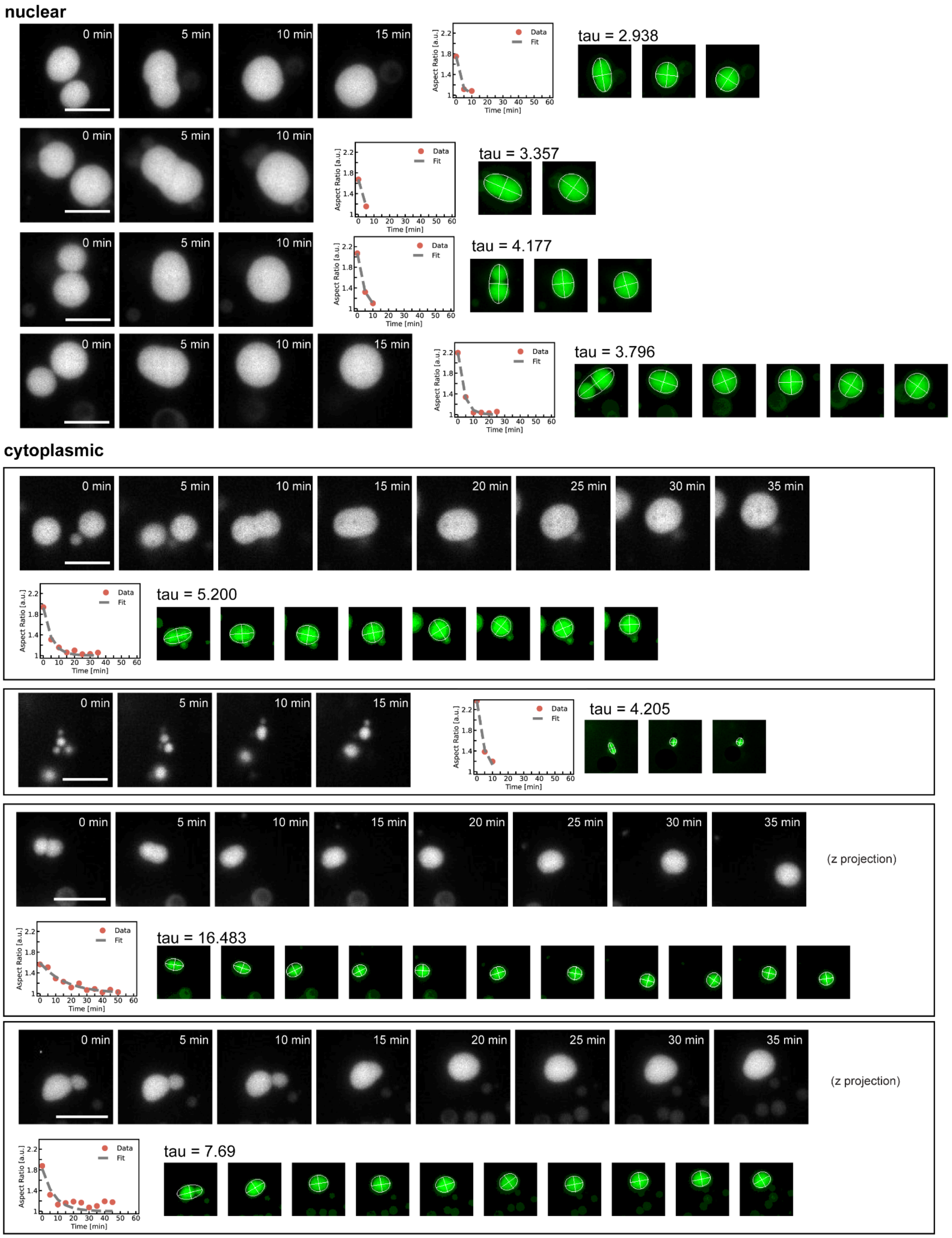


**Figure S17. Time-dependent coalescence of condensates in cells transfected to produce nanostar 15nt-3A-Br.** Mid plane confocal images show four nuclear and two cytoplasmic fusion events. Z projections show two cytoplasmic events with condensates on different z levels. Aspect ratios were calculated as the ratio of the major to minor axes from best-fit ellipses, as shown in the green micrographs. The dashed lines are fits of the exponential function $1 + Ae^{(-t/\tau)}$. Scale bar, 5 μm.

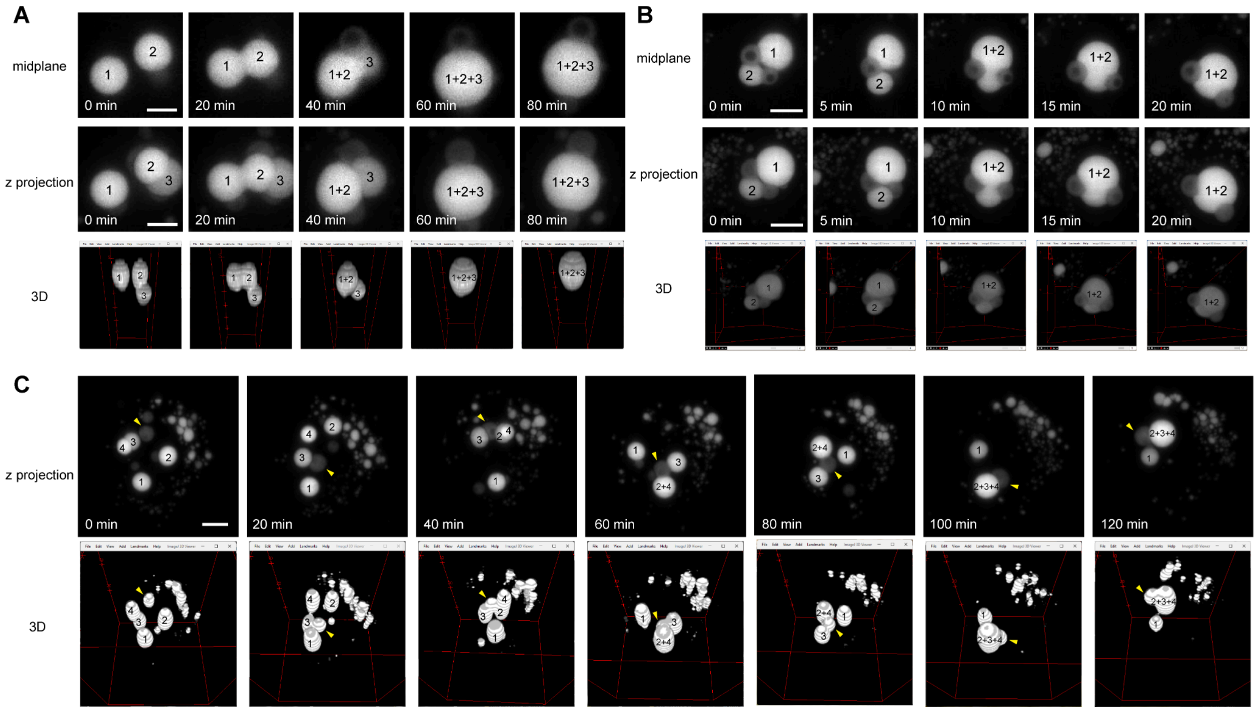


**Figure S18. Three complex fusion events in three dimensions.** Fusion events can involve multiple condensates and appear to be affected by the shells (as shown in the second and third events), where condensates fused but the shells remained distinct. Scale bar, 5 μm.

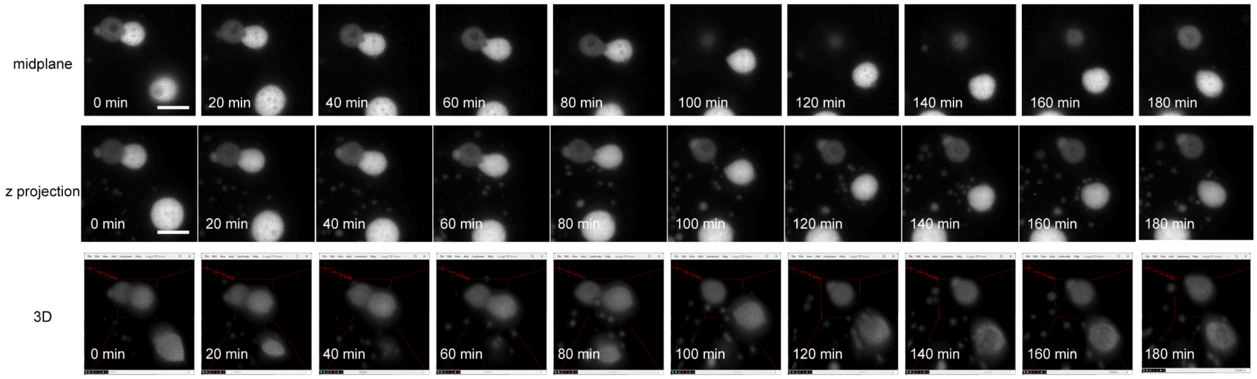


**Figure S19. Example splitting event.** Confocal images show a splitting event between a shell and a condensate. Scale bar, 5 μm.

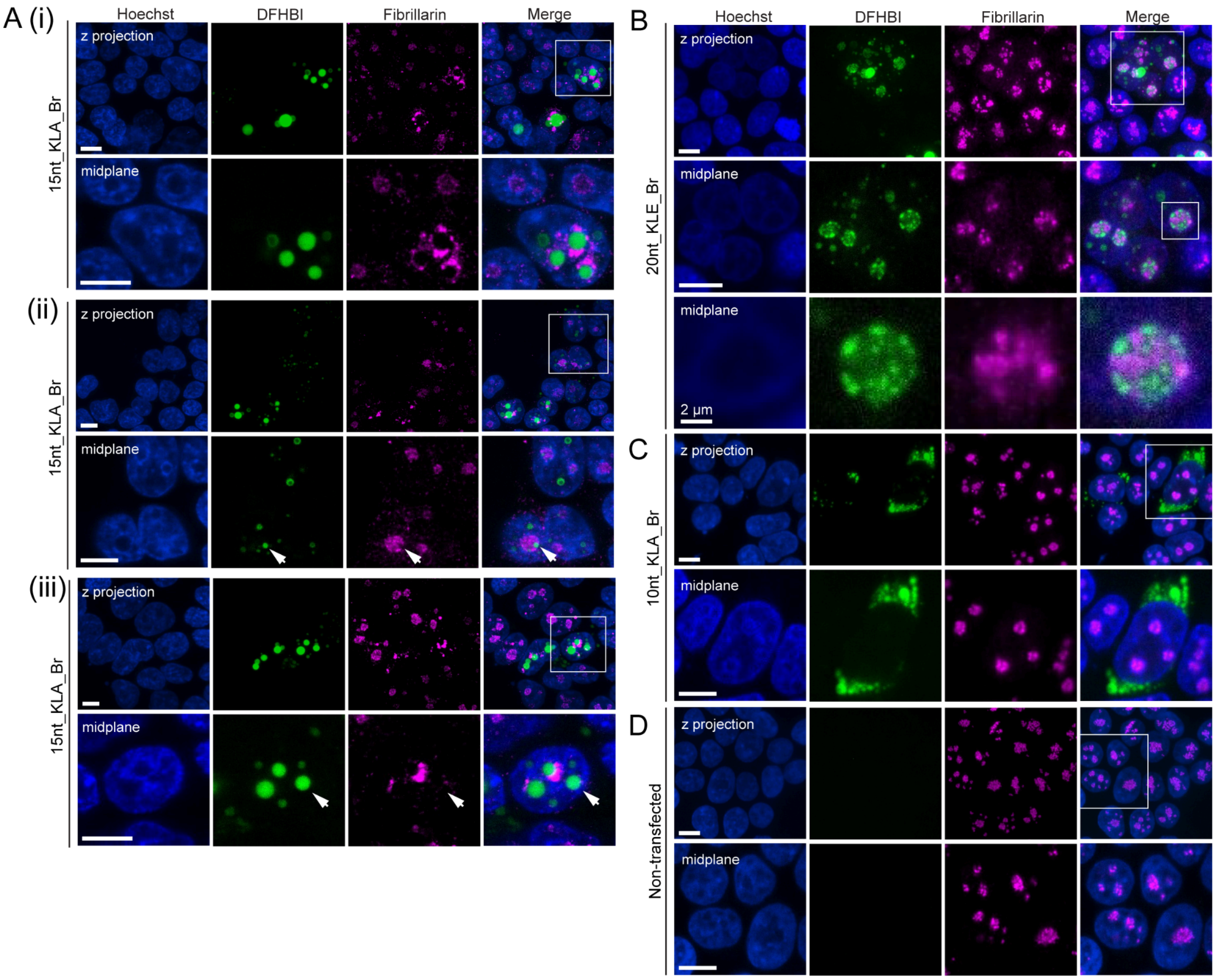


**Figure S20. RNA nanostars forming nuclear condensates colocalize with the nucleolus and exclude fibrillarin.** We tested nanostar designs that differ by arm length (10, 15 or 20 nucleotides) and kissing loop sequence (designs A or E). **A**, Z-projection and midplane confocal images of 15nt-A-Br showing condensates colocalizing with the nucleolus. In both high (i) and low (ii) expressing cells, condensates exclude fibrillarin, forming dark regions with reduced signal. Most nuclear condensates show fibrillarin accumulation on the surface but are not exclusively nucleolar, as in (iii) (white arrows). **B**, 20nt-E-Br nanostars form small puncta inside the nucleolus that exclude fibrillarin, similar to A(ii). **C**, 10nt-A-Br nanostars, which do not form nuclear condensates, show no change in fibrillarin localization. **D**, Fibrillarin distribution in non-transfected HEK293T cells. Representative images from three replicates. Scale bar, 10 µm.

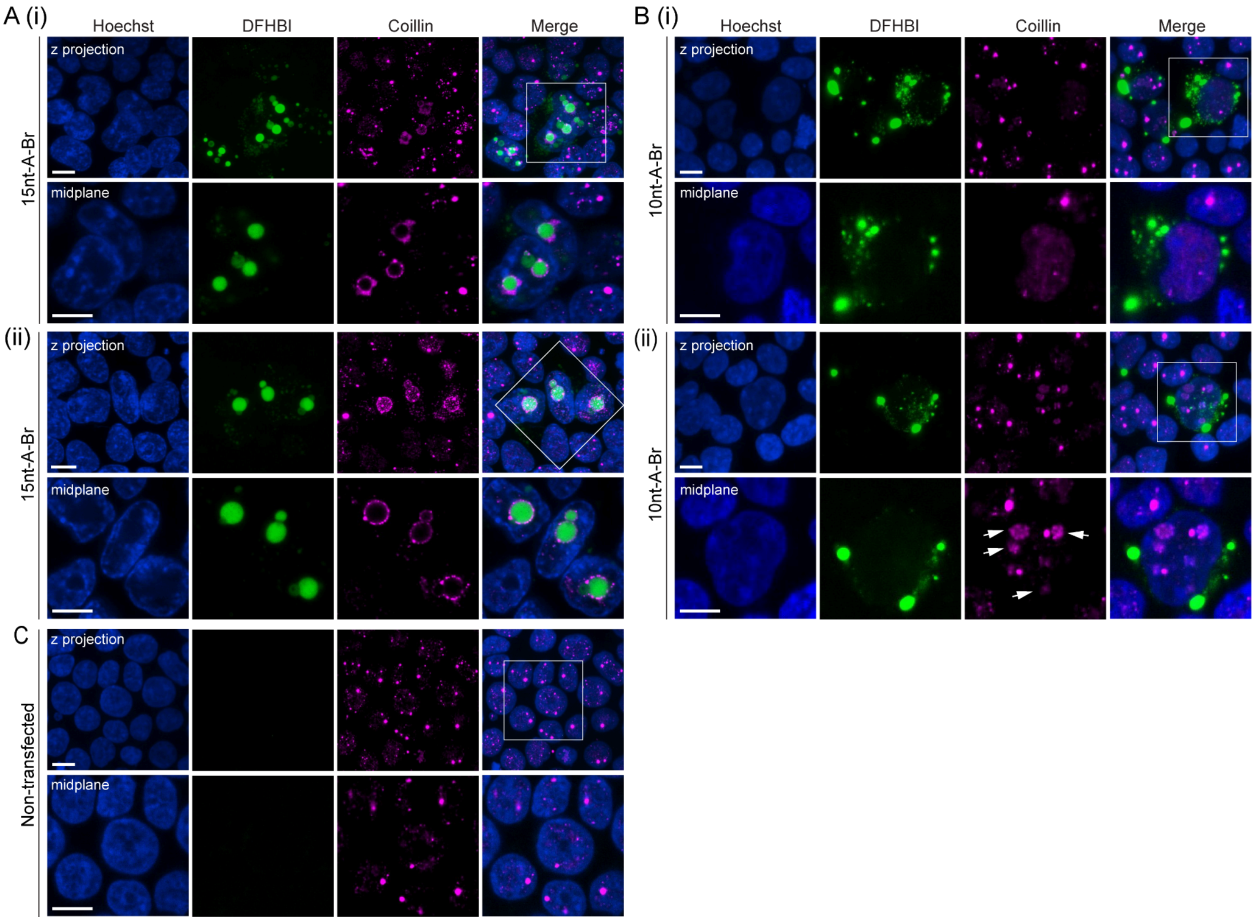


**Figure S21. Cajal bodies cluster on the surface of large nuclear condensates.** Like in Figure S20, we tested nanostar designs that differ by arm length (10, 15 or 20 nucleotides) and kissing loop sequence (designs A or E). **A,** Z-projection (top row) and single-slice confocal image (bottom row, white square in Z-projection) showing Cajal bodies clustered on the surface of condensates inside of nuclei. In both fields of view (i) and (ii), Coilin localizes at the surface of nuclear condensates. **B (i)**, RNA nanostars that form exclusively cytoplasmic condensates (10 nt arm length, see Fig. 2 of the manuscript) did not change Coilin localization in most nanostar-expressing cells. However, in some cases, localization of Coilin appeared altered (**B (ii)**). **C**, Cajal body in non-transfected HEK293T cells. Representative images from three replicates. Scale bar, 10 μm.

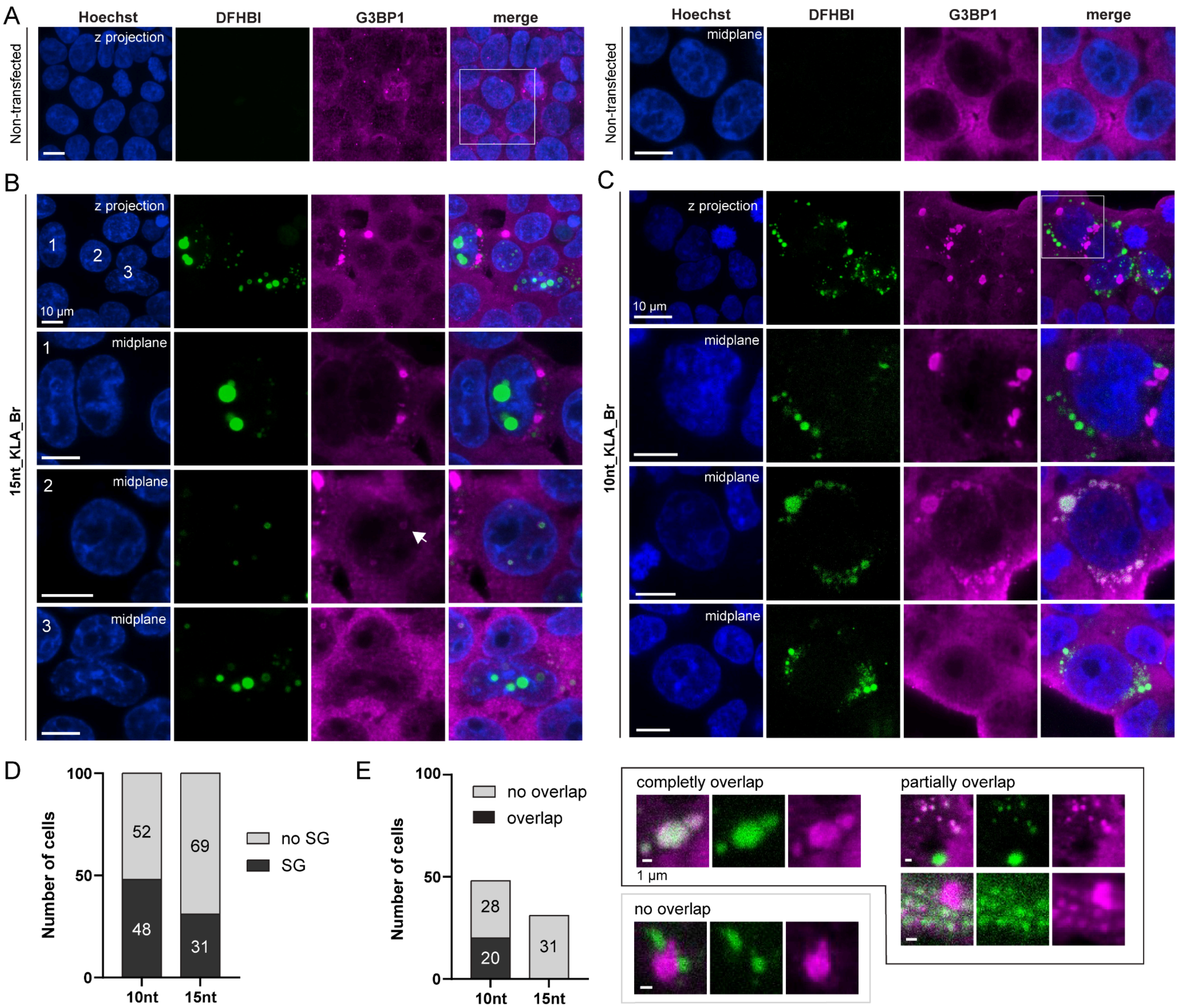


**Figure S22. The formation of abundant cytoplasmic condensates increases the likelihood of stress granule (SG) formation. A,** Z-projection (left) and zoom-in mid-plane confocal images (right) of untreated HEK293T cells stained with G3BP1 showing no SG formation. **B,** Z-projection (top row) and mid-plane images (bottom rows) of cells transfected with plasmids expressing 15-A-Br and stained with G3BP1. RNA condensation may correlate with SG formation without colocalizing with G3BP1, as in cell 1; this is not observed in all cells, as seen in cell 3. In cell 2, nanostars form shells that colocalize with G3BP1 (see Figure S23 for discussion). **C,** Z-projection (top row) and mid-plane images (bottom rows) of cells transfected with 10-A-Br and stained with G3BP1, showing examples of SG formation without colocalization (first row), with colocalization (second row, different FOV), and without SG formation (third row, different FOV). **D,** 10-A-Br nanostars show a higher tendency to trigger SG formation compared with 15-A-Br. **E,** Bar plot (left) showing that among cells with SG formation, only 10-A-Br displayed overlap between condensate and G3BP1 signals. Representative micrographs demonstrate the observed overlap patterns. Scale bar, 10 µm.

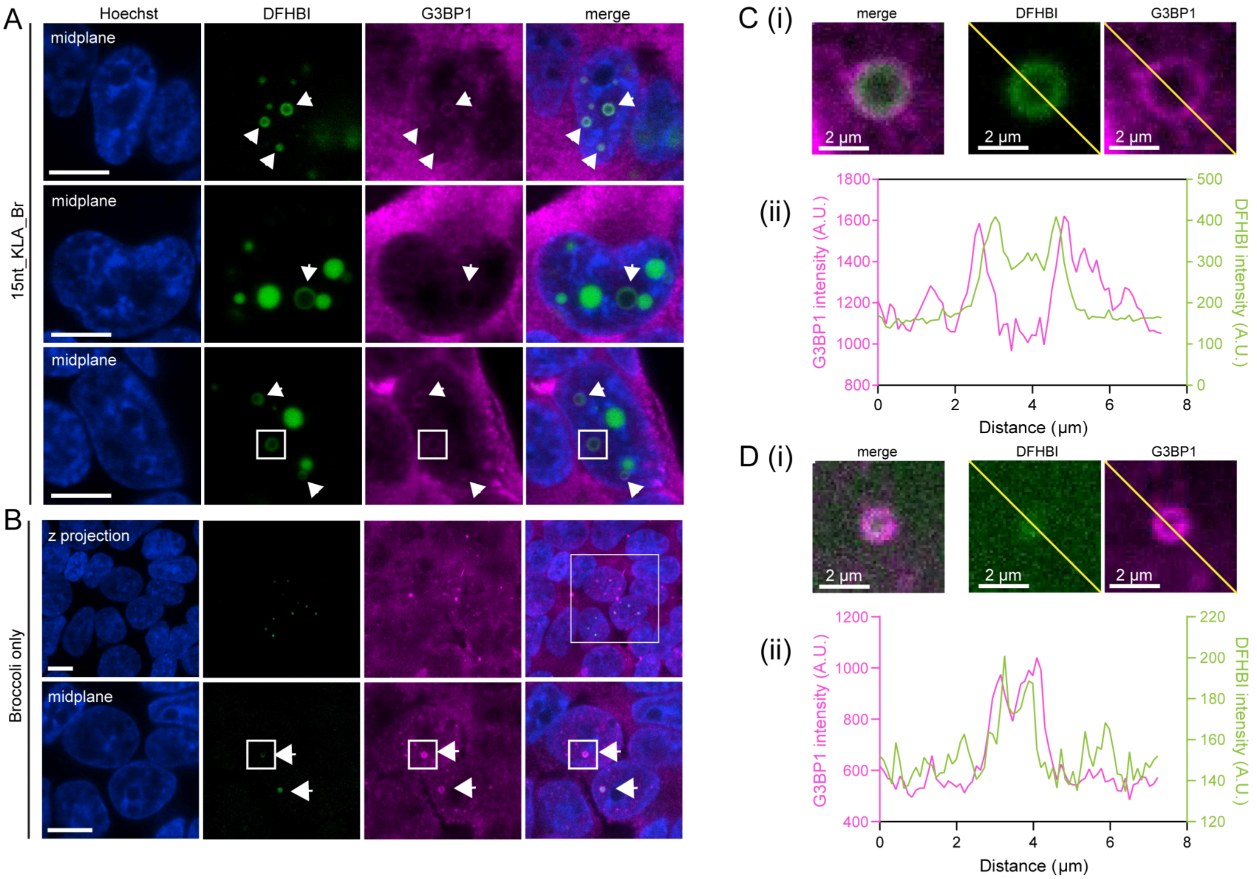


**Figure S23.** Shells are consistently observed across different fields of view in cells expressing RNA nanostars (**A**) or circularized Broccoli (**B**). G3BP1 colocalizes with shells but not with nuclear condensates, likely due to structure-mediated RNA decay. **C, D,** Pixel intensity profile of shells from nanostar-expressing cells (C, white square in A), or Broccoli-expressing cells (D, white square in B) illustrating its layered organization. Scale bar, 10 µm.

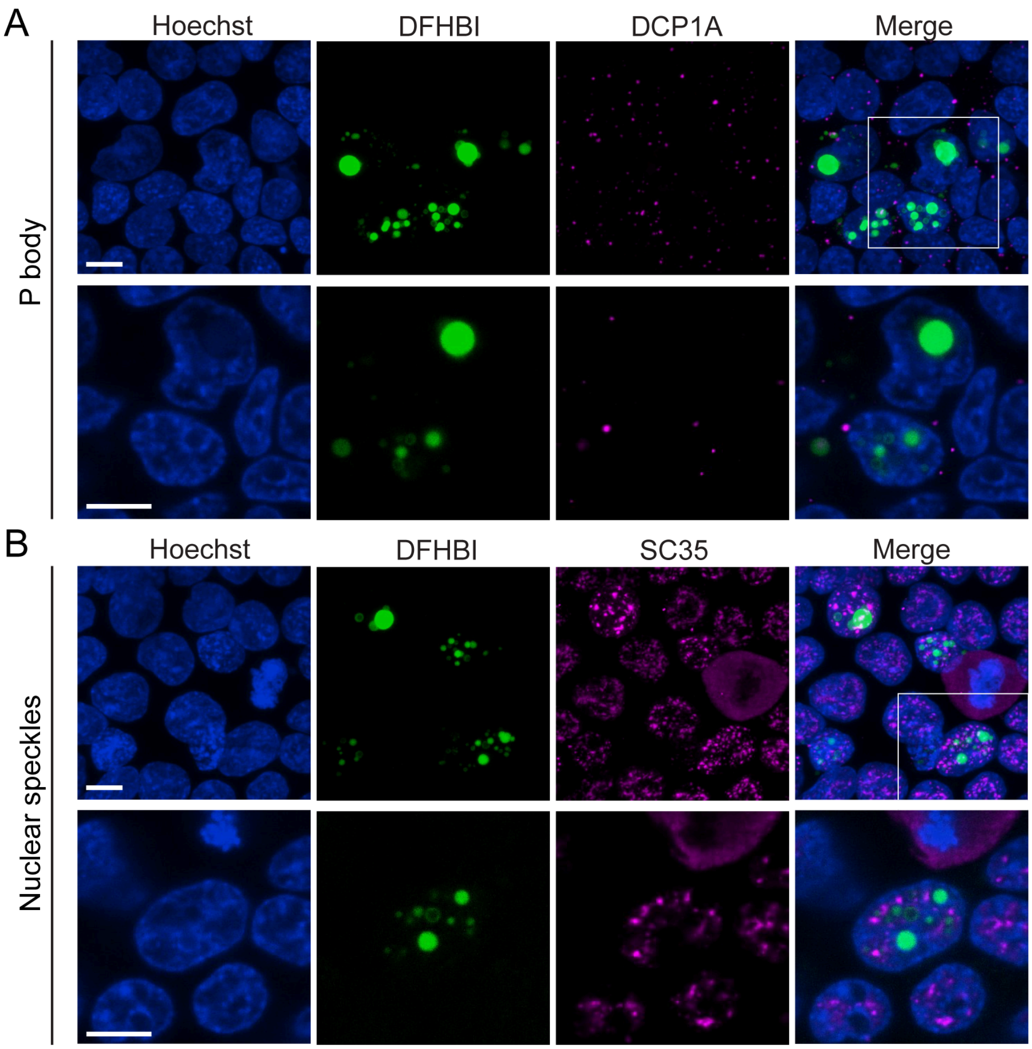


**Figure S24. Nuclear speckles (A) and P bodies (B) do not colocalize with RNA condensates.** Z-projection (top row) and single-slice (bottom row, white square in Z-projection) confocal images of cells were fixed 48 hours post-transfection. Images are representative of three replicates. Scale bar: 10 μm**.**

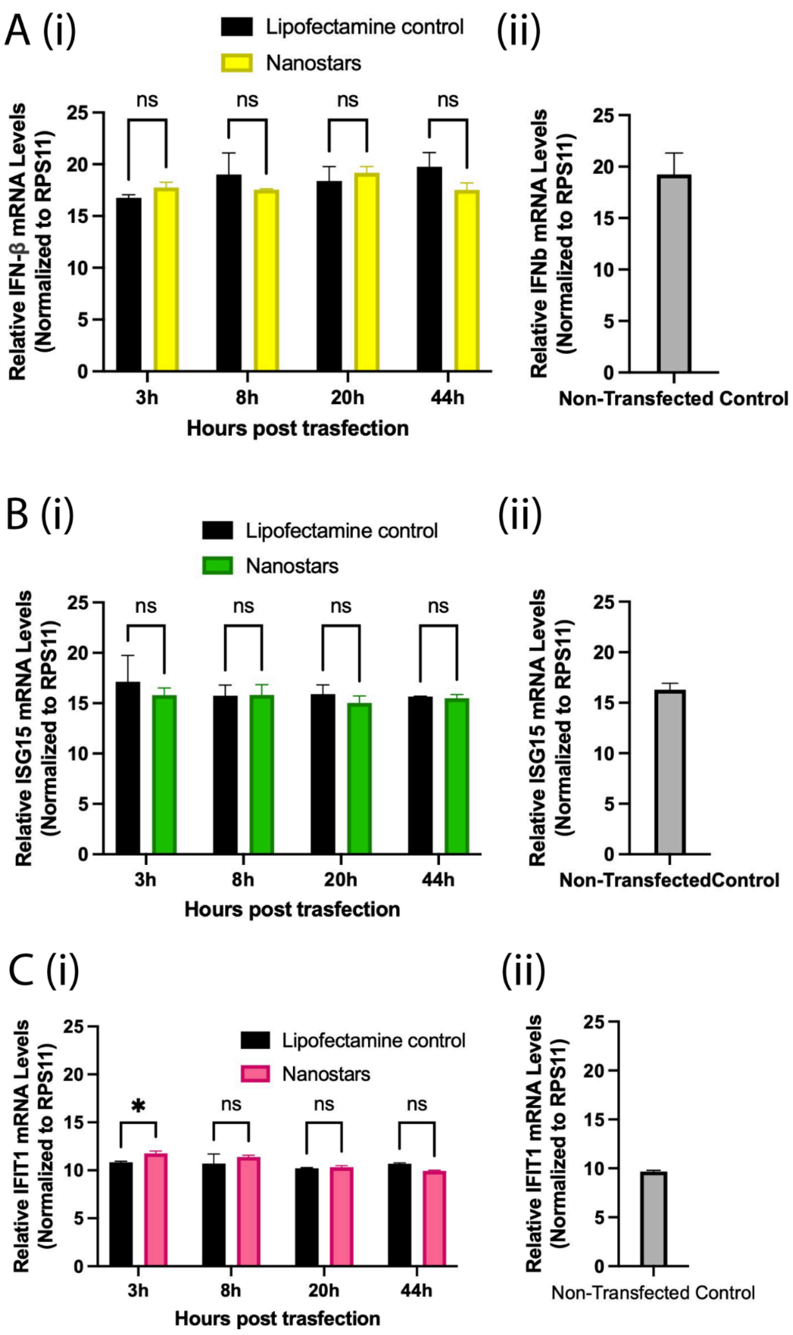


**Figure S25. Quantitative PCR (qPCR) analysis of interferon-stimulated gene (ISG) expression.** Expression levels of IFN-β (A), ISG15 (B), and IFIT1 (C) were measured and normalized to RPS11 in cells transfected with 15nt-3A-Br or treated with Lipofectamine alone at 3, 8, 20, and 44 hours post-transfection. (ii) Expression levels of the same genes in untreated cells are shown for comparison. All experiments were performed in triplicate. Statistical significance was determined by two-way ANOVA followed by multiple comparison tests.

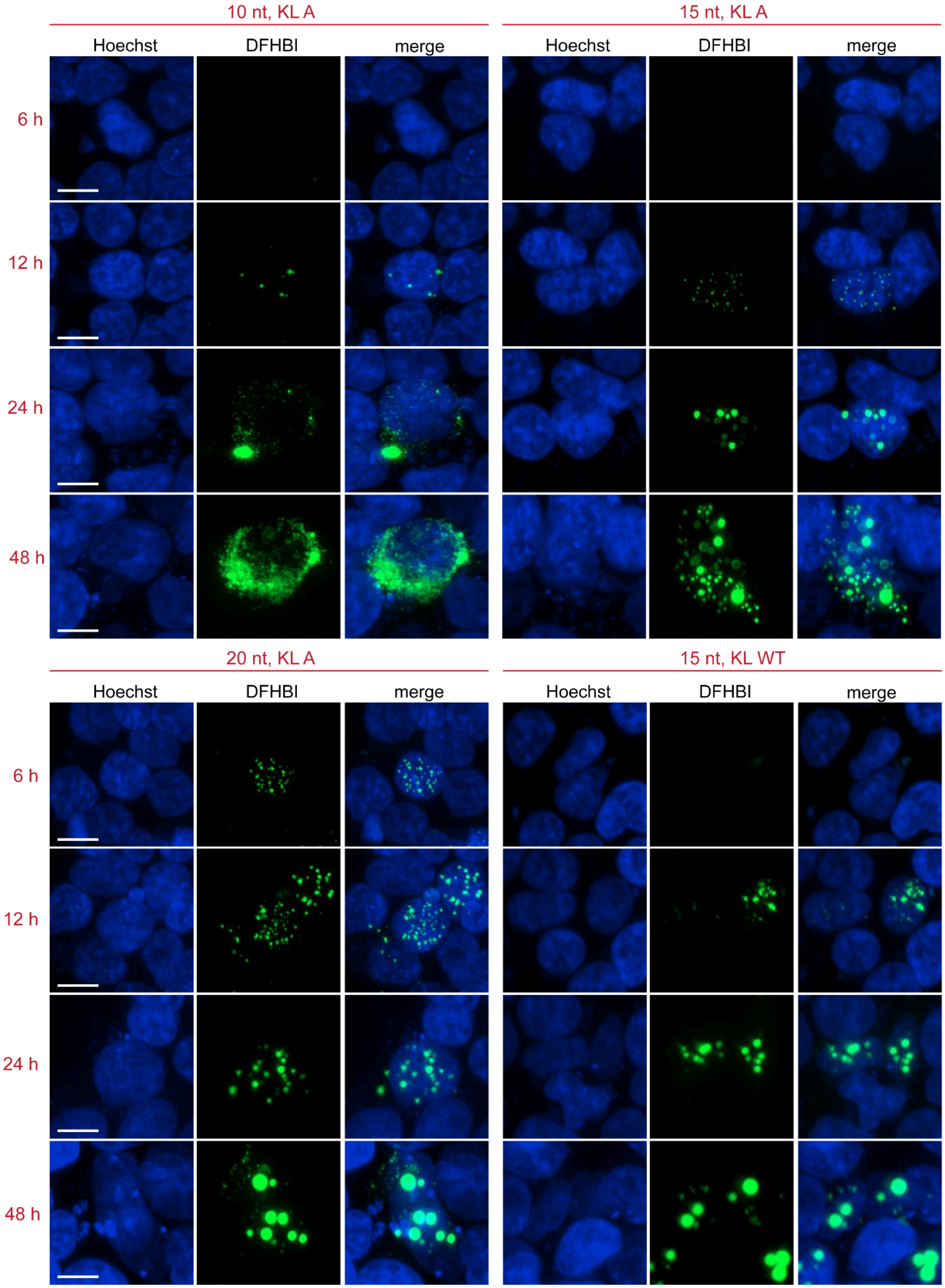


**Figure S26. Split-channel images of the micrographs shown in Figure 2.** Scale bar, 10 µm.

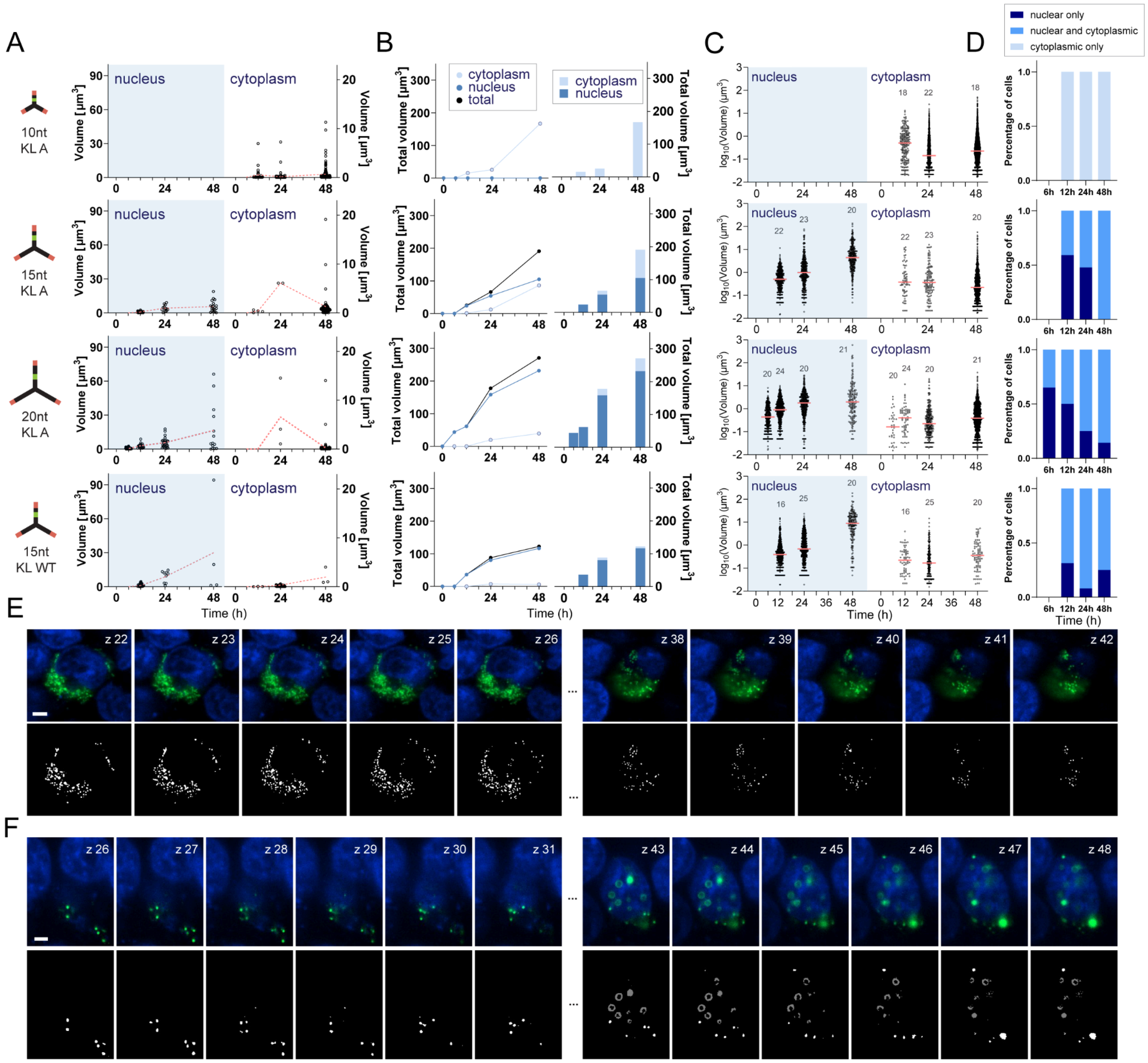


**Figure S27. Individual and total condensate volume of a representative cell. A,** Dot plots showing individual condensate volume at each time point. Dashed lines represent change of the mean volume. **B,** Line plots and grouped bar plots showing the change of total volume over time. Plots in B and C are for condensate volumes collected for an individual cell tracked over time in the corresponding row of A, indicated by white arrows. **C,** Dot plots showing temporal evolution of individual condensate volume (nuclear and cytoplasmic). Above each dot plot, we report the number of sampled cells at each time point, across multiple fields of view, from three replicates. **D,** Bar plots reporting the fraction of cells showing exclusively nuclear, nuclear and cytoplasmic, and exclusively cytoplasmic condensates at each time point. **E, F,** Single plane images (top) and the corresponding masks (bottom) for 10 nt KLA (E) and 15 nt KLA (F) at 48 hours. In the masks, cytoplasmic condensates are in white, while nuclear condensates are in gray. Increasing in slice number indicating intersecting from bottom to top of the cell. The masks were then reconstructed into 3D based on slice thickness for condensate volume quantification. Additional details are in Figure S28. Scale bar, 5 µm.

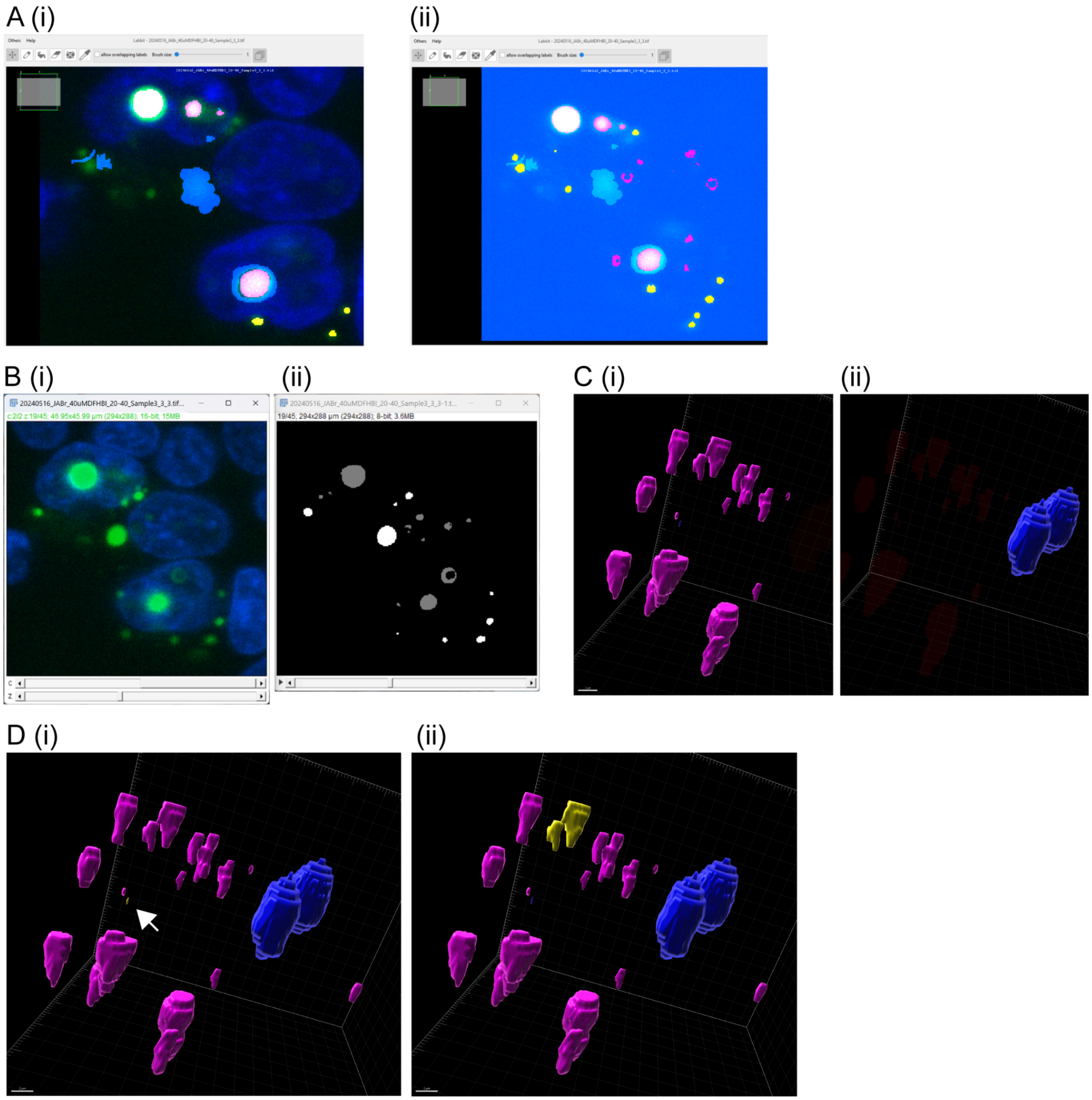


**Figure S28. Condensate volume quantification workflow. A,** For each confocal image, regions containing condensate-expressing cells were cropped and we manually labeled nuclear and cytoplasmic condensates (**i**) to generate a machine-learning segmentation pipeline using Labkit, an ImageJ plugin[9]. (ii). **B,** Comparison between a region of interest containing three condensate-expressing cells (i) and its mask (ii) that had condensates inside of the nucleus (gray) and the cytoplasm (white). **C,** Masks were analyzed in IMARIS to create surfaces for condensate populations inside (ii) and outside (i) of the nucleus. **D,** Data cleaning was performed to exclude single-voxel signals (i). Neighboring condensates that were fused were cropped manually. Statistics like condensate volume and number were automatically calculated by IMARIS and output for further analysis.

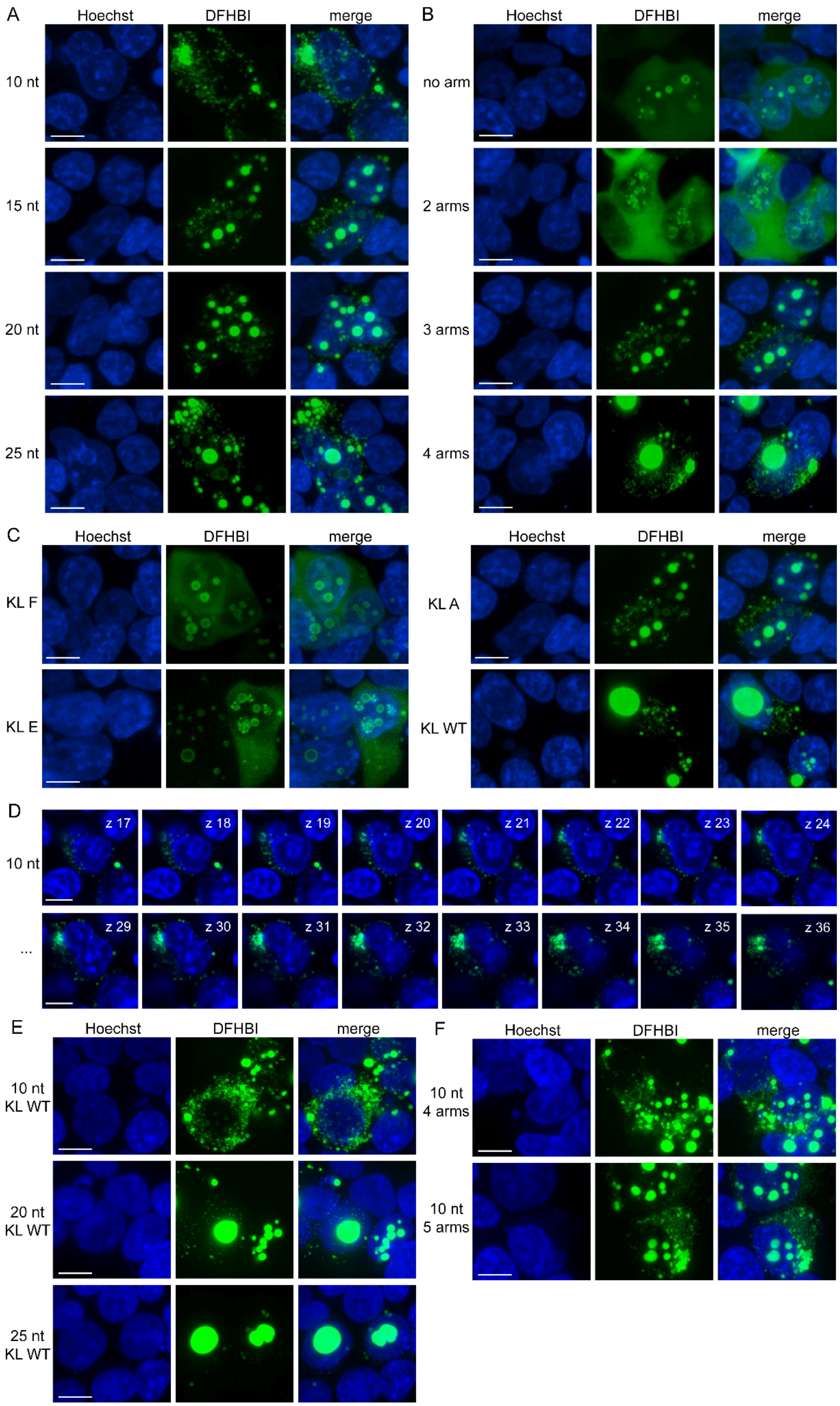


**Figure S29. Split-channel images (A, B, C, E, and F)** of the micrographs shown in Figure 3 and **single midplane images (D)** showing 10nt nanostar forming exclusively cytoplasmic condensates. Scale bar, 10 µm.

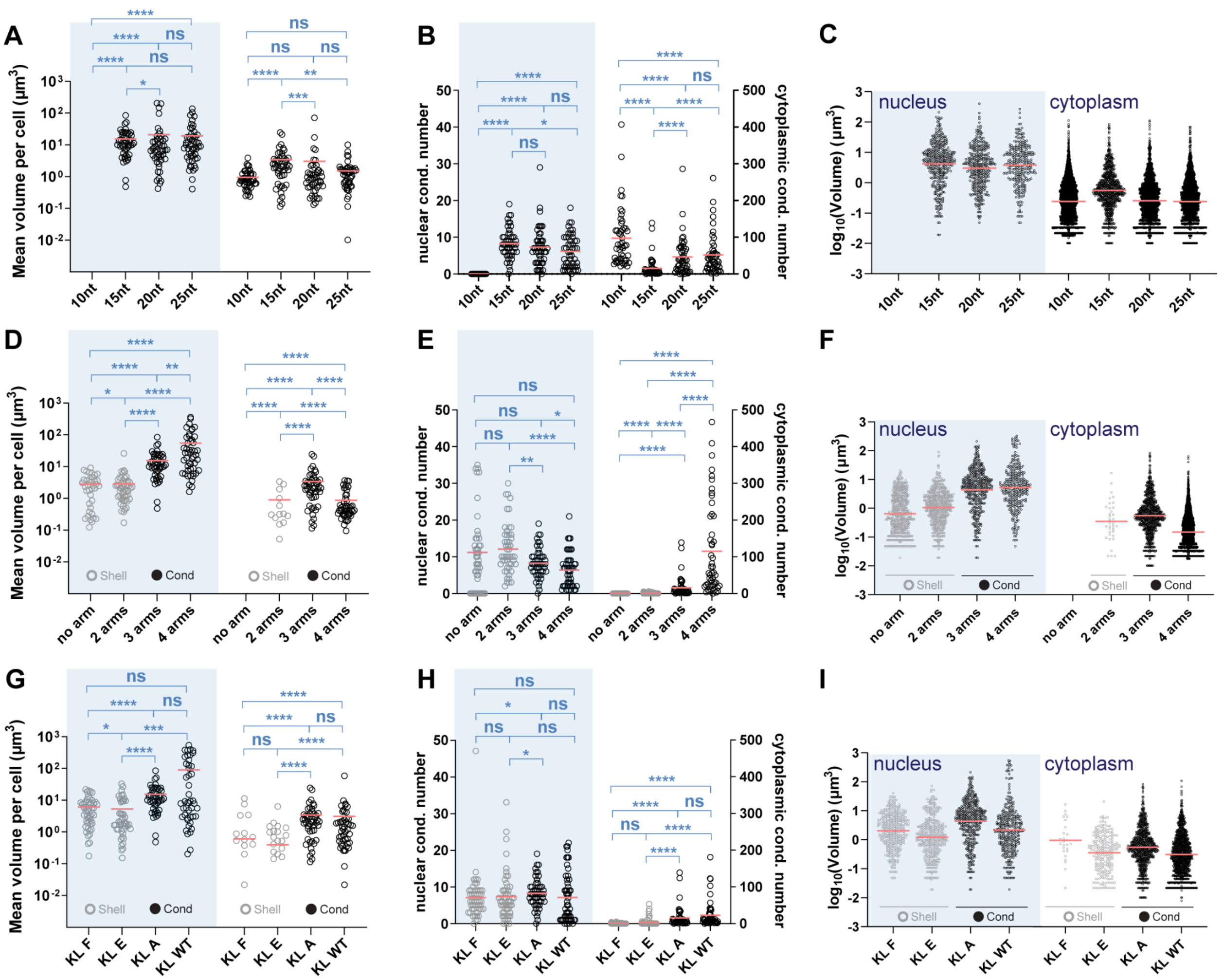

**Figure S30. Data shown in Figure 3 annotated with statistical significance. Distributions of condensate volume in the nucleus and cytoplasm comparing different arm lengths (A–C), arm numbers (D–F), and kissing loop sequences (G–I). A, D, G,** Average condensate volume per cell, where each dot represents a cell. **B, E, H,** Condensate number per cell, where each dot represents a cell. **C, F, I,** Individual condensate volumes, where each dot represents a condensate. Red lines indicate the mean. Cells with no nuclear or cytoplasmic condensates were assigned to have average condensate volume and condensate number per cell as zero. Significance was determined by the Mann-Whitney test, due to the zero-inflation. ; *p < 0.05; **p < 0.01; ***p < 0.001; ****p < 0.0001; ns, not significant. Zero values are not shown in panel A, D, G due to the use of the log axis. Detailed p-values are provided in Supplementary Table 3.

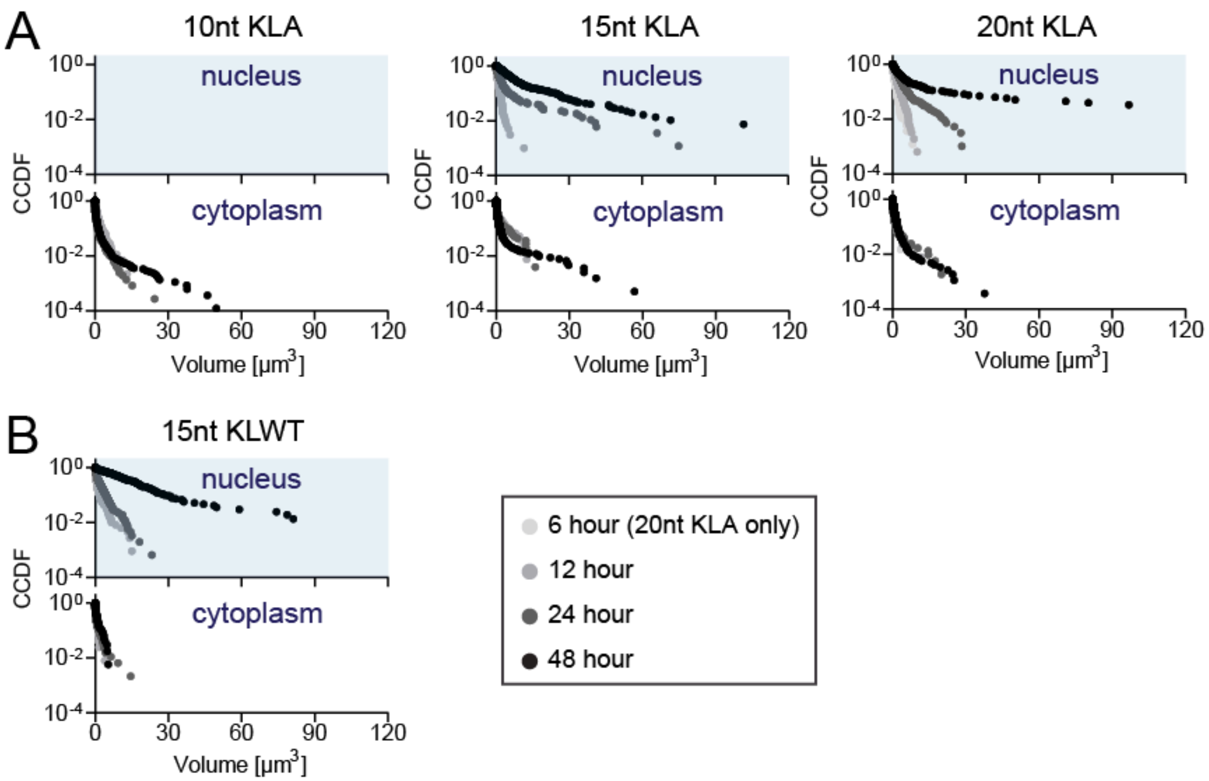


**Figure S31. Complementary cumulative distributions of condensate volume at different time points post-transfection.** Data in Fig. 2 and S27. To generate cumulative distribution functions (CCDFs) of condensate volumes, condensates were individually segmented in three dimensions and rescaled by the mean condensate volume in each cell. After rescaling, data from all cells were pooled and plotted collectively. Each dot represents a condensate.

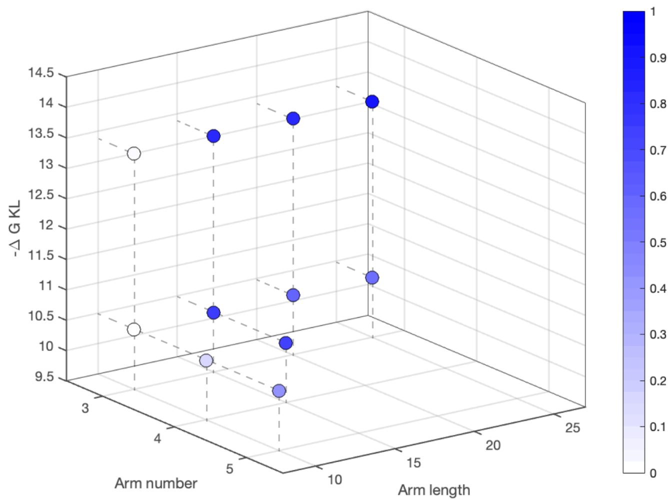


**Figure S32. Fraction of nuclear condensates as a function of design parameters.** This is a version of Figure 3J based on retrieved data. ΔG values were predicted by NUPACK using kissing loop RNA sequences assuming 37 °C, 1M strand concentration, 1M Na+. The color map shows normalized percentages of nuclear condensates over total condensate volume measured from 50 expressing cells. A cutoff effect was observed, as constructs with 10 nt arms formed exclusively cytoplasmic condensates, likely due to their smaller size relative to the nuclear pore complex. Overall, the ratio of nuclear condensates increases with longer arm length, higher arm number, or stronger kissing loop interactions.

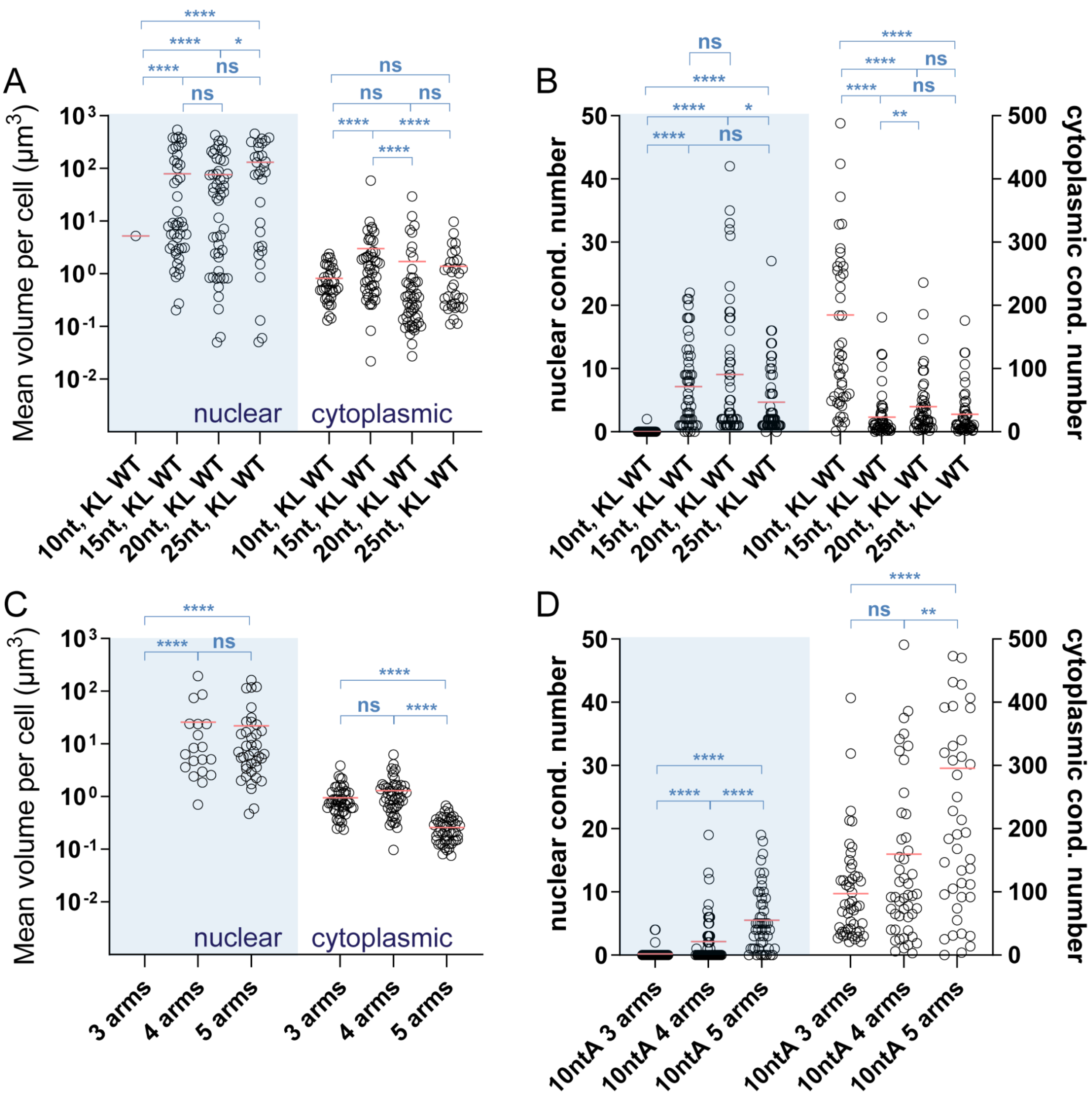


**Figure S33. Condensate volume quantification nanostar designs shown in Fig. 3K and 3L. A, C,** Average condensates' volume in the nucleus and cytoplasm of individual cells on a log scale. **B, D,** Number of condensates in the nucleus or cytoplasm measured in individual cells. Each black circle represents one cell. Red lines indicate the mean. Cells with no nuclear or cytoplasmic condensates were assigned to have average condensate volume and condensate number per cell as zero. Significance was determined by Mann-Whitney test due to the zero-inflation; *p < 0.05; **p < 0.01; ***p < 0.001; ****p < 0.0001; ns, not significant. Zero values are not shown in panel A, C due to the use of the log axis. Detailed p-values are provided in Supplementary Table 3.

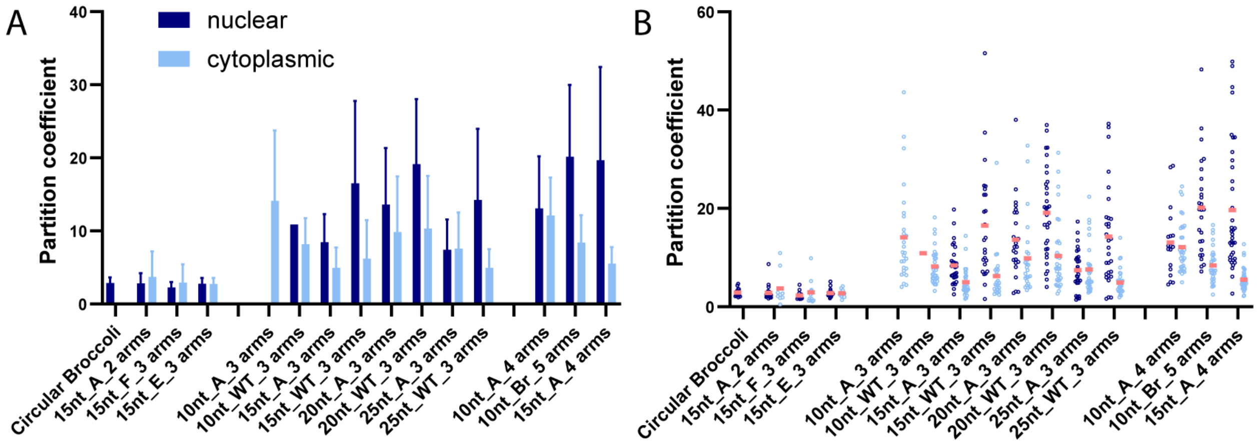


**Figure S34. Partition coefficients of all nanostar designs.** Partition coefficients, defined as the ratio of fluorescence intensity in the condensed phase to that in the dilute phase, are shown for each construct. A shows a summary plot of the individual data points presented in B. Nanostars forming only shells exhibit low partition coefficients (left four groups). In general, partition coefficients vary widely across cells. Nuclear condensates display higher mean partition coefficients than cytoplasmic condensates. Increasing nanostar arm length, kissing loop strength (from A to WT), or the number of arms (right three groups) all lead to higher mean partition coefficients in nuclear condensates.

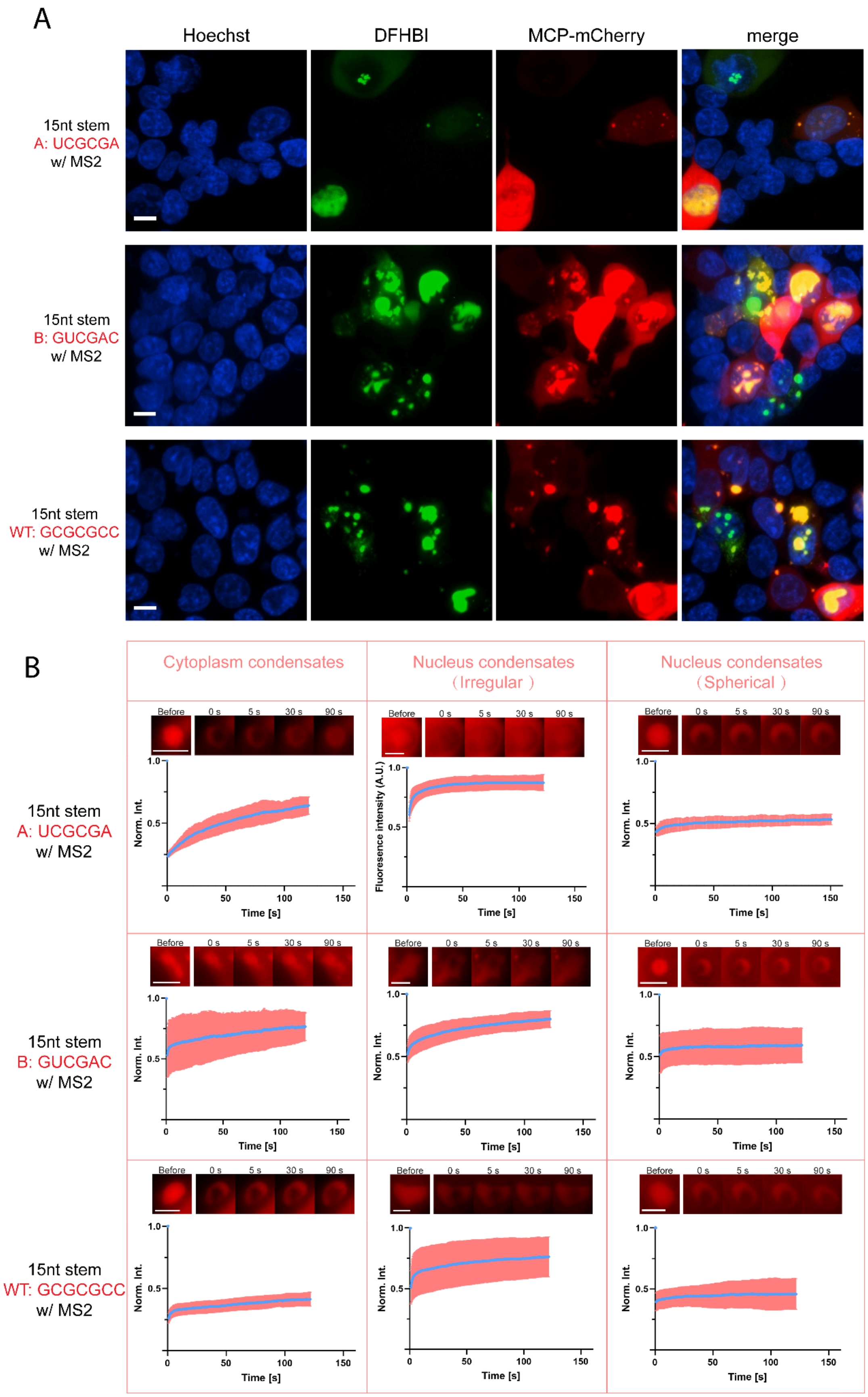
A
Hoechst
DFHBI
MCP-mCherry
merge
15nt stem
A: UCGCGA
w/ MS2
15nt stem
B: GUCGAC
w/ MS2
15nt stem
WT: GCGCGCC
w/ MS2
B
Cytoplasm condensates
Nucleus condensates (Irregular)
Nucleus condensates (Spherical)
Before
0 s
5 s
30 s
90 s
Norm. Int.
Fluorescence intensity (A.U.)
Time [s]
15nt stem
A: UCGCGA
w/ MS2
15nt stem
B: GUCGAC
w/ MS2
15nt stem
WT: GCGCGCC
w/ MS2

**Figure S35. mCherry recruitment to condensates is maintained across nanostar kissing loops of increasing strength, however protein diffusivity decreases. A**, Condensates formed from three different types of kissing loops can recruit MCP-mCherry. Plasmids expressing MCP-mCherry and RNA nanostar are transfected in a 1:9 ratio. The stoichiometry of plasmids delivered varies between individual cells, leading to diverse expression profiles. **B**, FRAP was performed on condensates colocalized with mCherry fluorescence, observed in panel A, including condensates located in the cytoplasm, and condensates located in the nucleus presenting a spherical or an irregular shape. Blue dots indicate mean values at the corresponding time point. Shaded areas indicate the standard error. The dark blue line indicates the fitted curve to the mean values from equation $Y = Y_0 + (Y_\infty - Y_0) * (1 - e^{-\frac{t}{\tau}})$. Mean and standard error were calculated from 3 fields of view each belonging to one replicate. Scale bars in A indicate 10 µm, in B indicate 5µm.

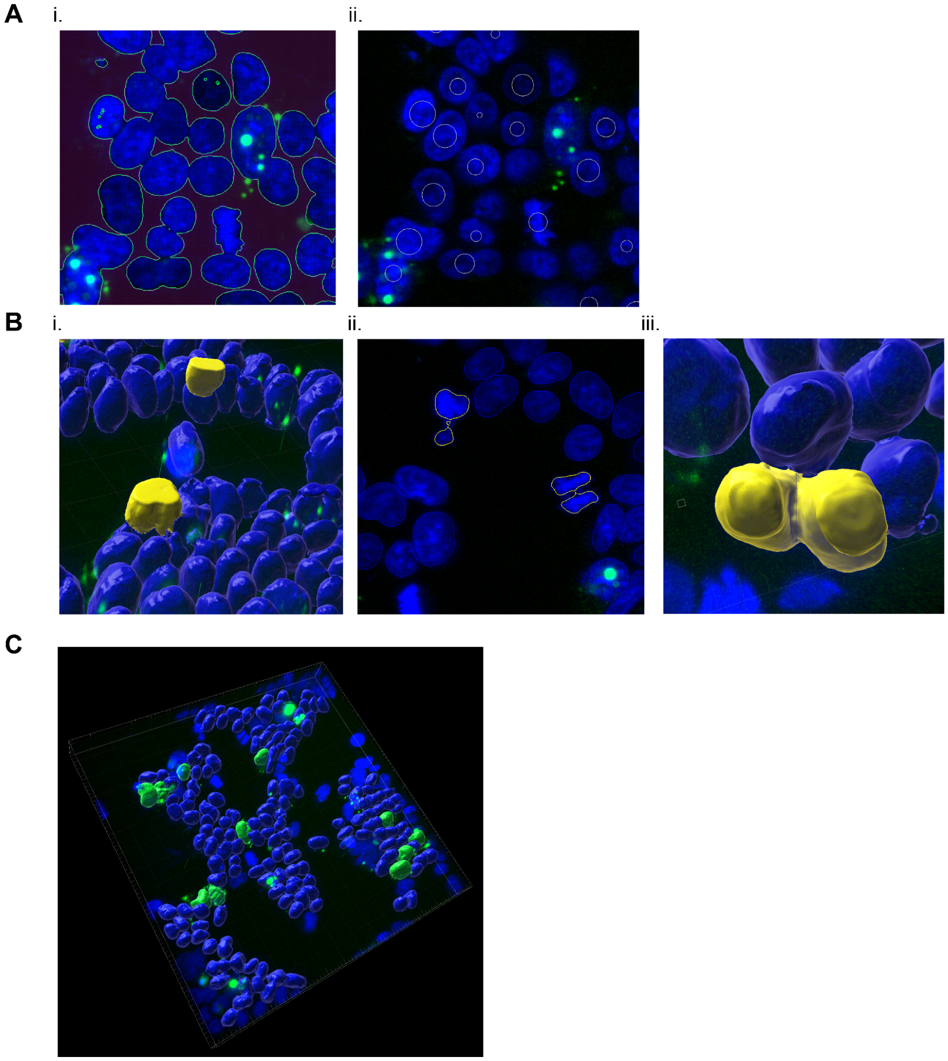


**Figure S36. Nucleus volume quantification workflow with IMARIS. A**, Example images showing the automated segmentation. We trained a machine learning model by manually labeling the nuclei and background on slices. The same model was applied to all confocal images and generated primary segmentation results as the example image showing in (i). To split touching nuclei, we applied a watershed method with estimated object size = 8 µm (ii), where circles indicate seed positions. **B**, Example images showing the data cleaning process. Nuclei split along the z-axis due to incomplete imaging (i) and nuclei with abnormally high pixel intensities indicating apoptosis or mitosis cells (ii) were manually excluded. Nuclei that failed to be split using the watershed algorithm were cut manually using the cutting tool in Imaris software to ensure proper segmentation of all nuclei (iii). Touching nucleus or nuclei without a distinguishable border were also removed. **C**, Example image showing classification results, where nuclei of condensate-expressing cells were highlighted in green and nuclei of non-expressing cells were in blue.

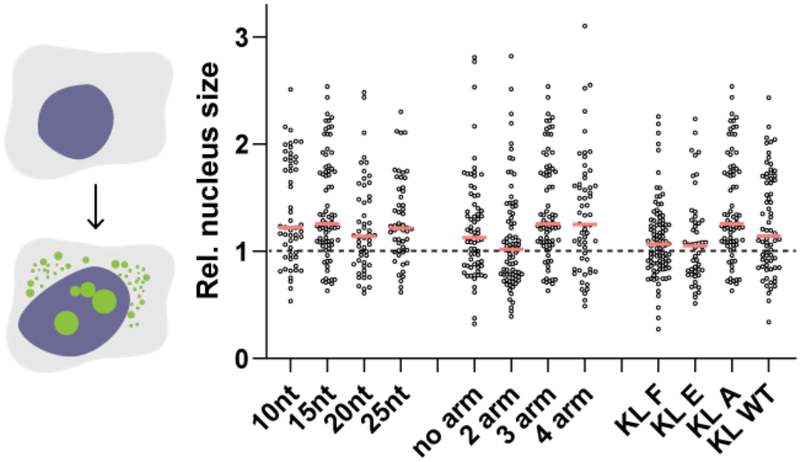


**Figure S37. Nucleus volume quantification for nanostar-expressing cells.** Nuclei of cells expressing nanostars exhibit an average volume that is consistently enlarged when compared to non-expressing cells (dashed line). The expression of condensate-forming nanostars seem to cause more nucleus enlargement, characterized by increased mean volume and by a shift in the overall volume distribution. This may be related to increased osmotic pressure within the nucleus caused by excessive production of circularized RNA. This analysis corresponds to data shown in Fig. 3 of the manuscript, imaged 48 hours after transfection. Each dot represents the nucleus volume from a nanostar-expressing cell that is normalized to the average nucleus volume of non-expression cells in the same experiment. Red lines indicate the mean.

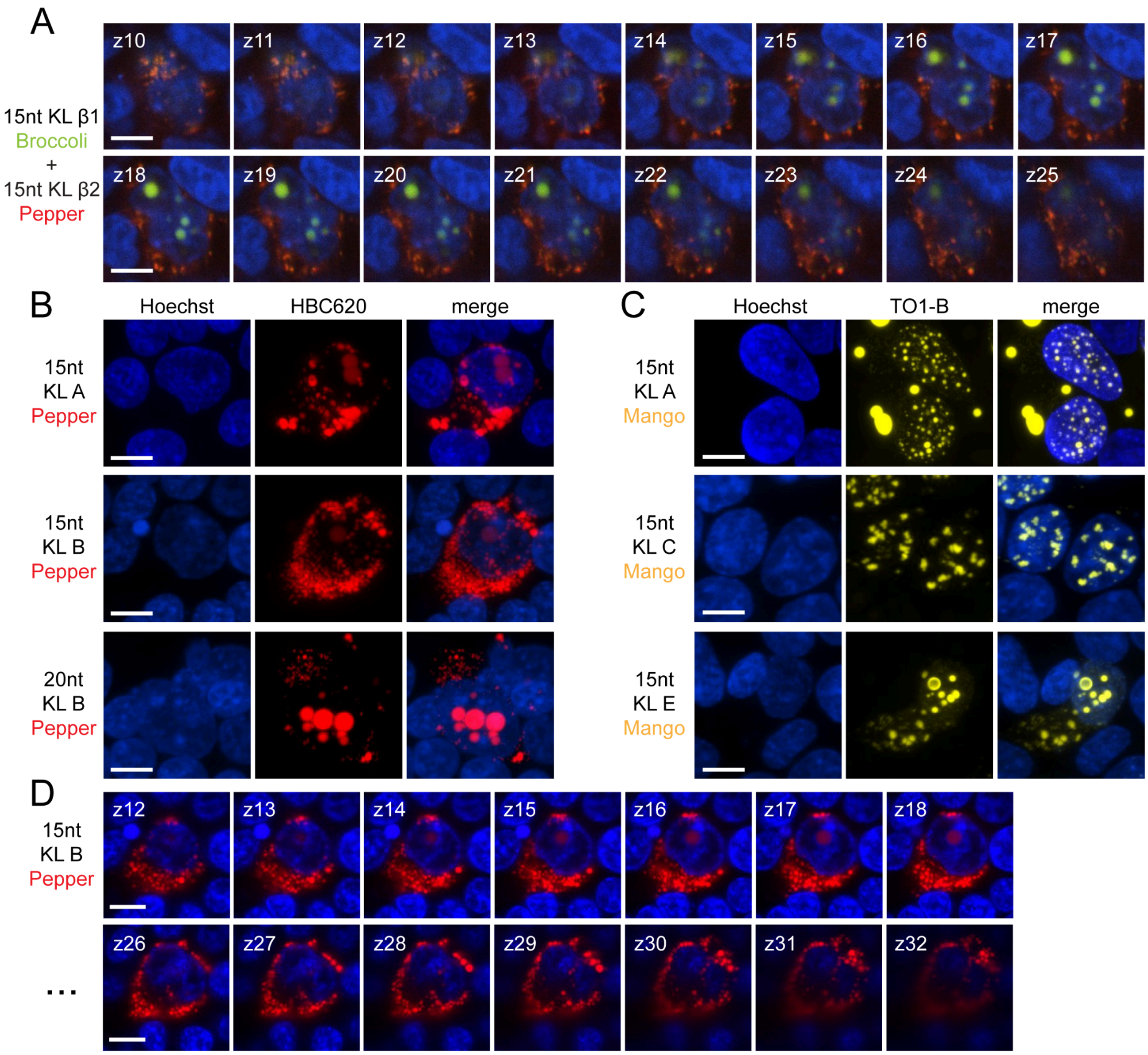


**Figure S38. Single midplane images and Split-channel images of the micrographs shown in Figure 4. A,** Single midplane images for the cell shown in Fig. 4A. **B,** Split channel images of nanostars tagged with Pepper. **C,** Split channel images of nanostars tagged with Mango. **D,** Single midplane images for the cell shown in Fig. 4C. 15nt KL B Pepper nanostars form primarily cytoplasmic condensates**.** Scale bar, 5 µm.

scale bar 10 um

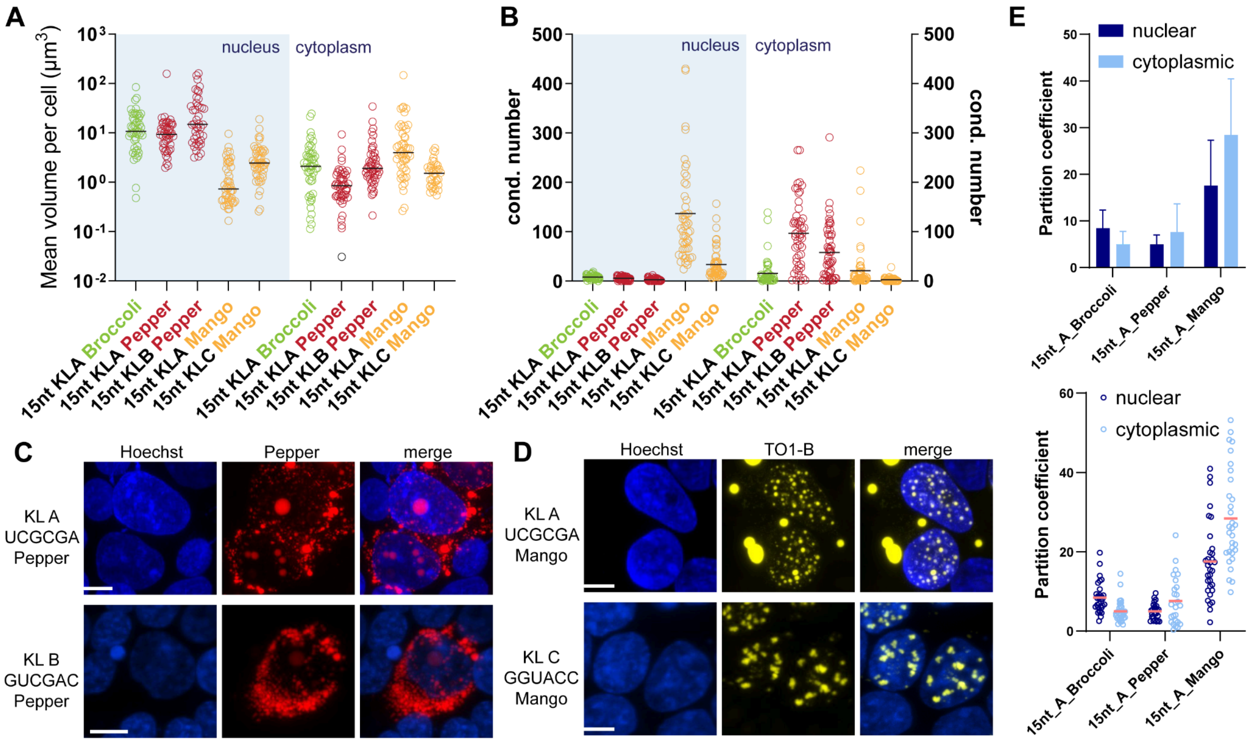


**Figure S39. The choice of fluorogenic aptamer influences nanostar condensation behavior. A,** Average condensate volume in the nucleus and cytoplasm of individual cells (log scale). **B,** Number of condensates in the nucleus or cytoplasm per cell. Each black circle represents one cell; red lines indicate the mean. **C, D,** Representative images of Pepper (C) and Mango (D) nanostars with different kissing loops. Although sharing the same stem (15 nt, KL A), nanostars display distinct localization and morphology depending on the aptamer used. Placement of UA base pairs alters kissing loop strength (KL A < KL B < KL C), leading to larger nuclear condensates for 15-B-Pp compared to 15-A-Pp, and larger, more nuclear-localized, and more irregular condensates for 15-C-Mango compared to 15-A-Mango. **E,** Bar plot (top) and dot plot (Bottom) showing partition coefficients of nanostars tagged with different aptamers. Scale bar: 10 μm.

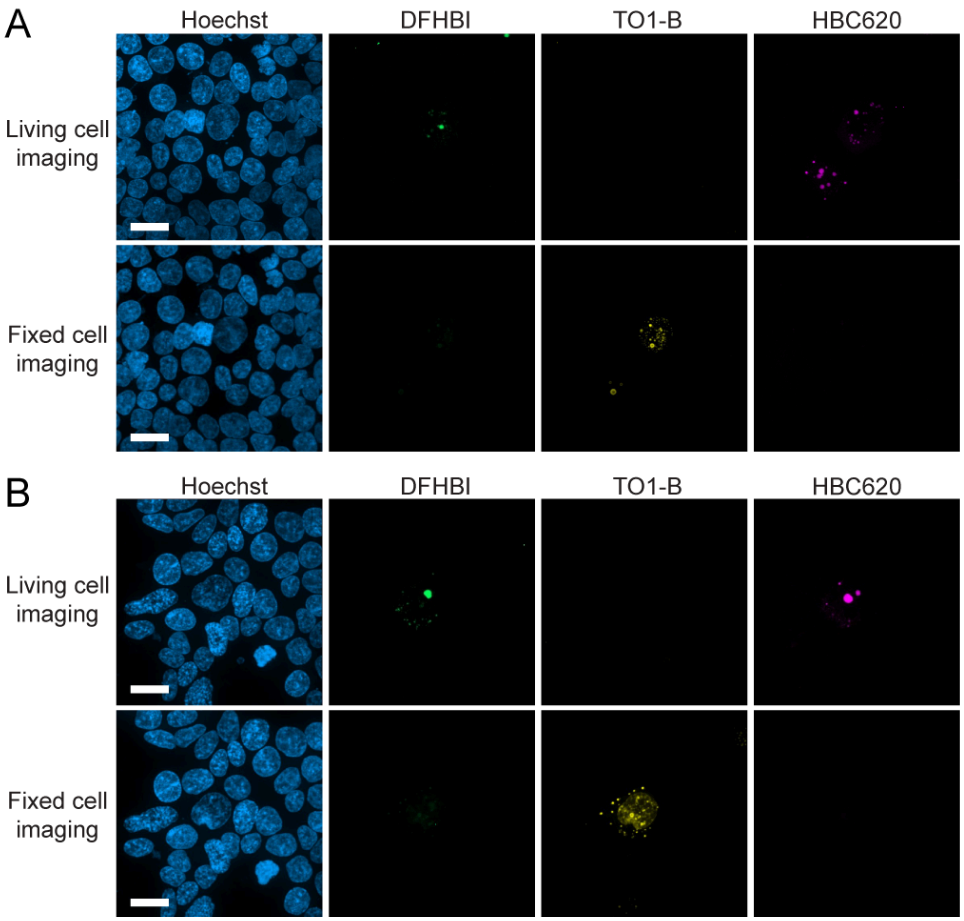


**Figure S40. Tri-color images were acquired by live cell imaging followed by on-stage fixation and fixed imaging at the same field of view. A**, Split channel images of condensates shown in Fig. 4D. **B**, Split channel images of condensates shown in Fig. 4K. For fixed cell imaging results, strong Mango signals in the yellow channel spillover into the green channel, resulting in weak green signals that colocalize with the yellow signal. All experiments are replicated three times. Scale bar, 20 μm.

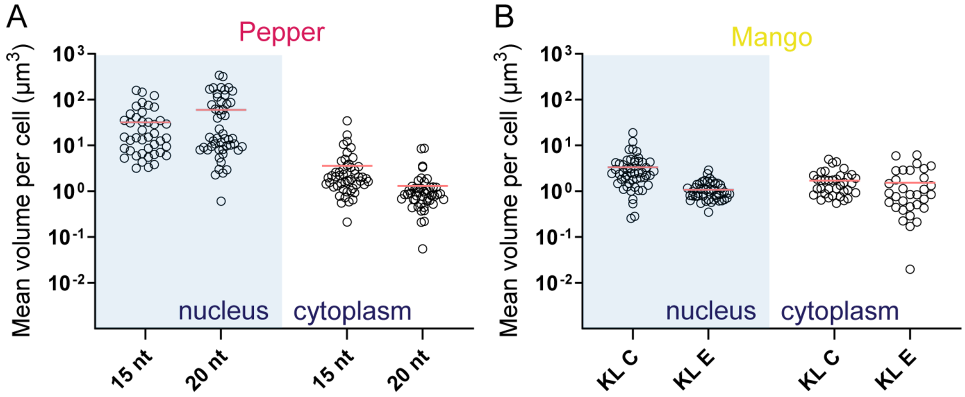


**Figure S41.** Average condensates' volume in the nucleus and cytoplasm of Pepper and Mango expressing cells on a log scale. Each black circle represents one cell. Red lines indicate the mean.

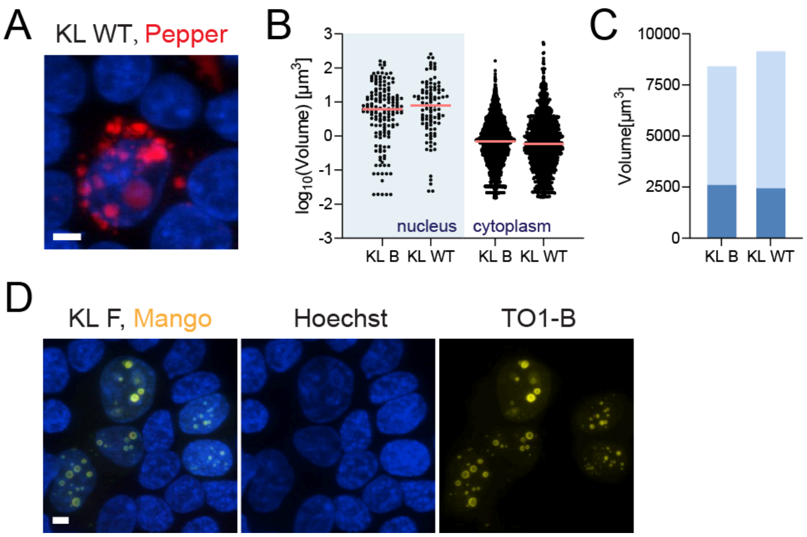


**Figure S42**. **A,** Z-projection of confocal microscopy images showing that Pepper-labeled nanostars (Fig. 4) with a WT kissing loop (GCGCGC) produce condensates that lose spherical shape, appearing as aggregates of smaller condensates. **B, C,** The cellular localization of the Pepper-WT nanostars does not change when compared to Pepper-B nanostars (Fig. 4). **D,** Z-projection of confocal microscopy images showing that Mango-tagged nanostars (Fig. 4) with kissing loop (AUAUAU) yields nuclear shells and diffusive Mango fluorescence in the cytoplasm, but not condensates were found. All experiments are conducted in triplicate. Scale bar, 5 µm.

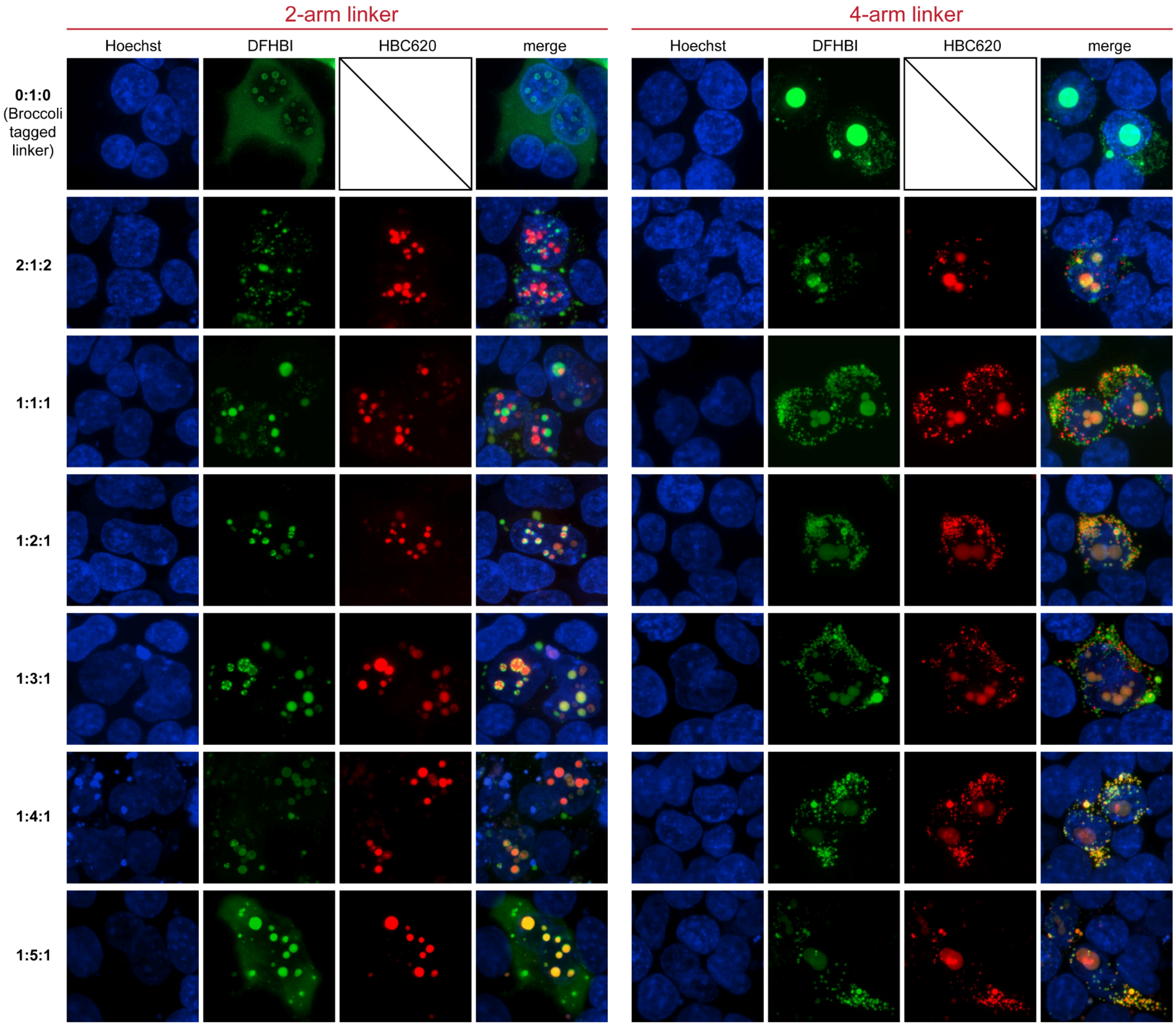

**Figure S43. Split-channel images of the micrographs shown in Figure 5.** Scale bar, 10 μm.

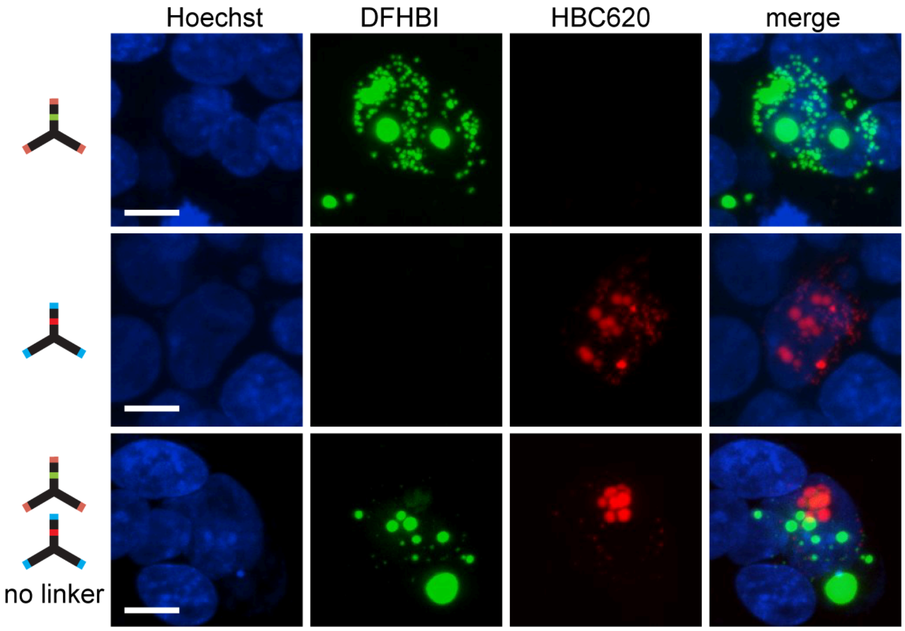


**Figure S44. Broccoli- and Pepper-labeled nanostars form orthogonal condensates in the absence of RNA linkers.** Nanostars with 20nt long arm and orthogonal kissing loops form non-mixing condensates. Images are representative of three replicates. Scale bar, 10 µm.

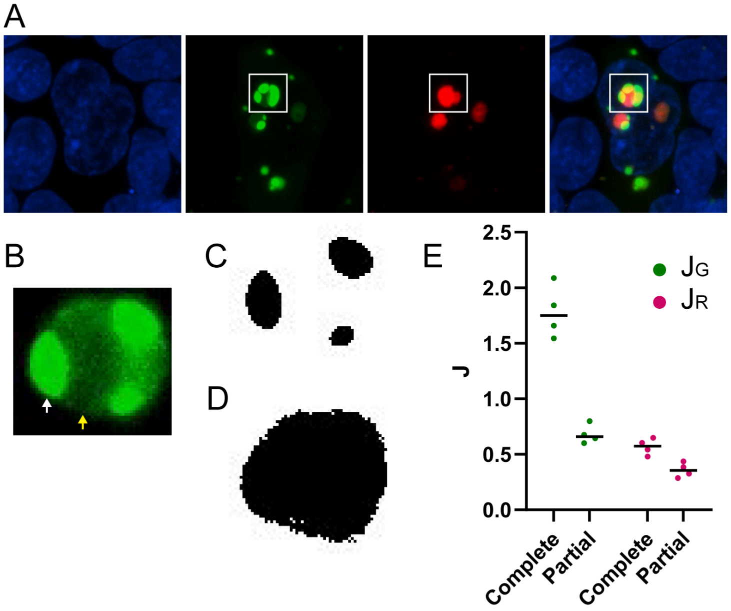


**Figure S45. Mixing index is influenced by non-uniform mixing. A,** Representative z-projection image showing a cell transfected with nanostars with kissing loops A and B, and RNA linker at 1:4:1 plasmid ratio. **B,** Image showing the Broccoli signal (nanostar A) at a single slice, indicating a non-uniform mixing in which we can identify a high signal region (white arrow) and low signal region (yellow arrow). Classifying this droplet as partial (with mask generated in **C**) or complete (with mask generated in **D**) mixing leading to significantly different mixing index, as shown in **E**. In the manuscript, all droplets from 1:4:1 ratio are defined as complete mixing.

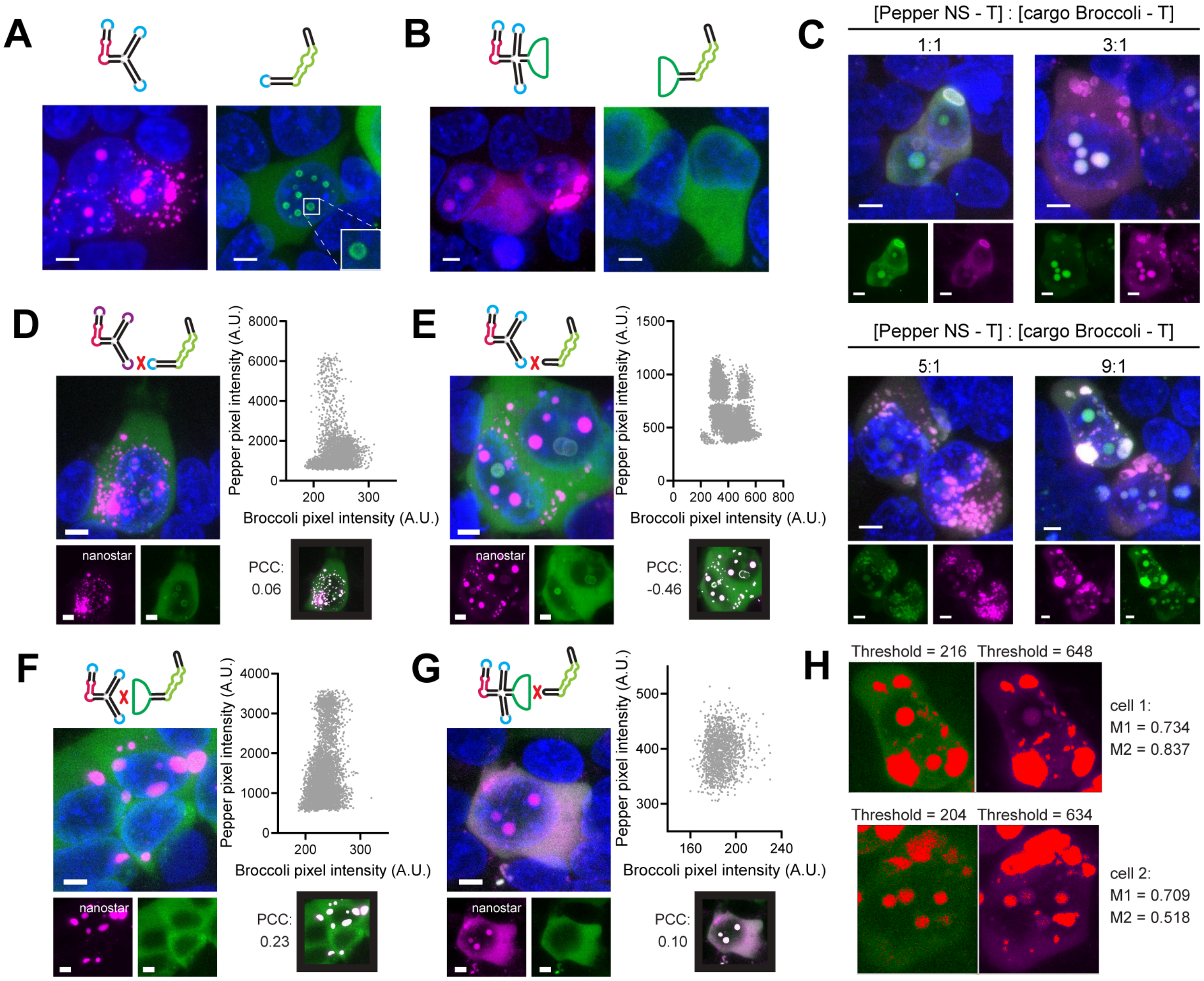


**Figure S46. A, B,** Confocal images of Pepper-labeled nanostars and Broccoli-labeled RNA molecules transfected individually. **C,** Confocal images of cells co-transfected with nanostar and RNA cargo plasmids at different transfection ratios. Steric hindrance from hybridization may impede condensate formation; therefore, a higher nanostar-to-cargo ratio is required for condensation. **D-G,** Micrographs and scatter plots showing control experiments in Fig. 6. Pearson correlation coefficients (PCC) show no colocalization between Pepper-labeled nanostars and Broccoli-labeled RNA molecules with absence of matched sequences. **H,** Thresholded images (highlighted in red) of cells 1 and 2 in Fig. 6F and their corresponding M1 and M2 coefficients confirmed colocalization between nanostars labeled with Pepper in magenta and the target RNA labeled with Broccoli in green. Although visual colocalization was evident, the low Broccoli signal intensity in cell 2 produced a low PCC value of 0.18 in Fig. 6F. Manders' colocalization coefficients, which quantify the fraction of overlapping signal, help resolve this issue. Scale bar, 5 μm.

#### Supplementary Tables

| Caption | Sequence |
|---|---|
| • Yellow is the U6+27 promoter.<br>• Teal is the 5’ ribozyme sequence.<br>• Cyan is the 3’ ribozyme sequence.<br>• The ^ symbols represent cleavage sites catalyzed by ribozymes and mark the circularized expression product sequence.<br>• Green is the RNA sequence of interest (15nt-KLA-Br, here). The [] symbols mark an arm. The <> symbols mark a kissing loop (KL A, here).<br>• Red is the U6 terminator.<br><br>Sites used for cloning (underlined):<br>• SalI (GTCTAC) and XbaI (TCTAGA), inserting using this pair generates linear RNA nanostar (As demonstrated in Supplementary Fig. 2)<br>• NotI (GCGGCCGC) and SacII (CCGCGG), inserting using this pair generates circularized RNA nanostar | RNA sequence<br>GAGGGCCUAUUUCCCAUGAUUCCUUCAUAUUUGCAU<br>AUACGAUACAAGGCUGUUAGAGAGAUAAUUAGAAUU<br>AAUUUGACUGUAAACACAAAGAUAUUAGUACAAAAUA<br>CGUGACGUAGAAAGUAAUAAUUUCUUGGGUAGUUU<br>GCAGUUUUAAAAUUAUGUUUUAAAAUGGACUAUCAU<br>AUGCUUACCGUAACUUGAAAGUAUUUCGAUUUCUUG<br>GCUUUAUAUAUCUUGUGGAAAGGACGAAACACC<br>GUGCUCGCUUCGGCAGCACAUAUACUAGUCGACGG<br>GCCGCACUCGCCGGUCCCAAGCCCGGAUAAAAUGG<br>GAGGGGGCGGGAAACCGCCU^AACCAUGCCGAGUG<br>CGGCCGC[GCGAGAGCGCUGCCC<AAUCGCGAA<br>>GGGCAGCGCUCUCGC]AA[GGAUGG**ACGGUCG**<br>**GGUCC**GAGGGC<AAUCGCGAA>GCCCUC**GUCG**<br>**AGUAGAGUGUGGG**CCAUCC]AA[GCGUUCACAC<br>UGACC<AAUCGCGAA>GGUCAGUGUGAACGC]C<br>CGCGGUCGGCGUGGACUGUAG^AACACUGCCAAUG<br>CCGGUCCCAAGCCCGGAUAAAAGUGGAGGGUACAG<br>UCCACGCUCUAGAGCGGACUUCGGUCCGCUUUUU<br><br>DNA sequence (Template strand)<br>GAGGGCCTATTTCCCATGATTCCTTCATATTTGCATATA<br>CGATACAAGGCTGTTAGAGAGATAATTAGAATTAATTTG<br>ACTGTAAACACAAAGATATTAGTACAAAATACGTGACGT<br>AGAAAGTAATAATTTCTTGGGTAGTTTGCAGTTTTAAAA<br>TTATGTTTTAAAATGGACTATCATATGCTTACCGTAACTT<br>GAAAGTATTTCGATTTCTTGGCTTTATATATCTTGTGGA<br>AAGGACGAAACACC<br>GTGCTCGCTTCGGCAGCACATATACTAGTCGACGGGC<br>CGCACTCGCCGGTCCCAAGCCCGGATAAAATGGGAG<br>GGGGCGGGAAACCGCCT^AACCATGCCGAGTGCGGC<br>CGC[GCGAGAGCGCTGCCC<AATCGCGAA>GGGCAG<br>CGCTCTCGC]AA[GGATGG**ACGGTCGGGTCC**GAGGGC<br><AATCGCGAA>GCCCTC**GTCGAGTAGAGTGTGGG**CC<br>ATCC]AA[GCGTTCACACTGACC<AATCGCGAA>GGTC<br>AGTGTGAACGC]CCGCGGTCGGCGTGGACTGTAG^AA<br>CACTGCCAATGCCGGTCCCAAGCCCGGATAAAAGTGG<br>AGGGTACAGTCCACGCTCTAGAGCGGACTTCGGTCC<br>GCTTTTT |

**Supplementary Table 1.** Elements in the expression cassette using the TORNADO expression system.

| Kissing loop nomenclature | Sequence |
|---|---|

| | |
|---|---|
| A | 5’ - AA UCGCGA A - 3’ |
| B | 5’ - AA GUCGAC A - 3’ |
| C | 5’ - AA GGUACC A - 3’ |
| E | 5’ - AA GUAUAC A - 3’ |
| F | 5’ - AA UAUAUA A - 3’ |
| Beta 1 (non-palindromic) | 5’ - AA GCUACG A - 3’ |
| Beta 2 (non-palindromic) | 5’ - AA CGUAGC A - 3’ |

**Supplementary Table 2.** Nomenclature of kissing loops used in the manuscript.

| Panel | Plot | Location | Group | p-value | notation |
|---|---|---|---|---|---|
| B | Mean volume per cell | nucleus | 10nt vs 15nt | p<0.0001 | **** |
| B | Mean volume per cell | nucleus | 10nt vs 20nt | p<0.0001 | **** |
| B | Mean volume per cell | nucleus | 10nt vs 25nt | p<0.0001 | **** |
| B | Mean volume per cell | nucleus | 15nt vs 20nt | 0.0401 | * |
| B | Mean volume per cell | cytoplasm | 10nt vs 15nt | p<0.0001 | **** |
| B | Mean volume per cell | cytoplasm | 15nt vs 20nt | 0.0006 | *** |
| B | Mean volume per cell | cytoplasm | 15nt vs 25nt | 0.0018 | ** |
| B | Condensate number per cell | nucleus | 10nt vs 15nt | p<0.0001 | **** |
| B | Condensate number per cell | nucleus | 10nt vs 20nt | p<0.0001 | **** |
| B | Condensate number per cell | nucleus | 10nt vs 25nt | p<0.0001 | **** |
| B | Condensate number per cell | nucleus | 15nt vs 25nt | 0.0162 | * |
| B | Condensate number per cell | cytoplasm | 10nt vs 15nt | p<0.0001 | **** |
| B | Condensate number per cell | cytoplasm | 10nt vs 20nt | p<0.0001 | **** |

| | | | | | |
|---|---|---|---|---|---|
| B | Condensate number per cell | cytoplasm | 10nt vs 25nt | p<0.0001 | **** |
| B | Condensate number per cell | cytoplasm | 15nt vs 20nt | p<0.0001 | **** |
| B | Condensate number per cell | cytoplasm | 15nt vs 25nt | p<0.0001 | **** |
| D | Mean volume per cell | nucleus | 3 arms vs 4 arms | 0.0052 | ** |
| D | Mean volume per cell | nucleus | no arms vs 2 arms | 0.0364 | * |
| D | Mean volume per cell | nucleus | no arms vs 4 arms | p<0.0001 | **** |
| D | Mean volume per cell | nucleus | no arms vs 3 arms | p<0.0001 | **** |
| D | Mean volume per cell | nucleus | 2 arms vs 3 arms | p<0.0001 | **** |
| D | Mean volume per cell | nucleus | 2 arms vs 4 arms | p<0.0001 | **** |
| D | Mean volume per cell | cytoplasm | 3 arms vs 4 arms | p<0.0001 | **** |
| D | Mean volume per cell | cytoplasm | 2 arms vs 3 arms | p<0.0001 | **** |
| D | Mean volume per cell | cytoplasm | 2 arms vs 4 arms | p<0.0001 | **** |
| D | Mean volume per cell | cytoplasm | no arms vs 2 arms | p<0.0001 | **** |
| D | Mean volume per cell | cytoplasm | no arms vs 3 arms | p<0.0001 | **** |
| D | Mean volume per cell | cytoplasm | no arms vs 4 arms | p<0.0001 | **** |
| D | Condensate number per cell | nucleus | 2 arms vs 3 arms | 0.0027 | ** |
| D | Condensate number per cell | nucleus | 2 arms vs 4 arms | p<0.0001 | **** |
| D | Condensate number per cell | nucleus | 3 arms vs 4 arms | 0.0125 | * |
| D | Condensate number per cell | cytoplasm | 3 arms vs 4 arms | p<0.0001 | **** |
| D | Condensate number | cytoplasm | no arms vs 4 | p<0.0001 | **** |

| | | | | | |
|---|---|---|---|---|---|
| | per cell | | arms | | |
| D | Condensate number per cell | cytoplasm | 2 arms vs 4 arms | p<0.0001 | **** |
| D | Condensate number per cell | cytoplasm | no arms vs 2 arms | p<0.0001 | **** |
| D | Condensate number per cell | cytoplasm | 2 arms vs 3 arms | p<0.0001 | **** |
| D | Condensate number per cell | cytoplasm | no arms vs 3 arms | p<0.0001 | **** |
| F | Mean volume per cell | nucleus | KL E vs KL WT | 0.0007 | *** |
| F | Mean volume per cell | nucleus | KL E vs KL F | 0.0192 | * |
| F | Mean volume per cell | nucleus | KL E vs KL A | p<0.0001 | **** |
| F | Mean volume per cell | nucleus | KL F vs KL A | p<0.0001 | **** |
| F | Mean volume per cell | cytoplasm | KL F vs KL WT | p<0.0001 | **** |
| F | Mean volume per cell | cytoplasm | KL E vs KL WT | p<0.0001 | **** |
| F | Mean volume per cell | cytoplasm | KL F vs KL WT | p<0.0001 | **** |
| F | Mean volume per cell | cytoplasm | KL E vs KL A | p<0.0001 | **** |
| F | Condensate number per cell | nucleus | KL E vs KL A | 0.0288 | * |
| F | Condensate number per cell | nucleus | KL F vs KL A | 0.0247 | * |
| F | Condensate number per cell | cytoplasm | KL F vs KL WT | p<0.0001 | **** |
| F | Condensate number per cell | cytoplasm | KL E vs KL WT | p<0.0001 | **** |
| F | Condensate number per cell | cytoplasm | KL F vs KL A | p<0.0001 | **** |
| F | Condensate number per cell | nucleus | KL E vs KL A | p<0.0001 | **** |
| K | Mean volume per cell | nucleus | 15nt, KL WT vs 10nt, KL WT | p<0.0001 | **** |
| K | Mean volume per cell | nucleus | 20nt, KL WT vs | p<0.0001 | **** |

| | | | | | |
|---|---|---|---|---|---|
| | | | 10nt, KL WT | | |
| K | Mean volume per cell | nucleus | 25nt, KL WT vs 10nt, KL WT | p<0.0001 | **** |
| K | Mean volume per cell | nucleus | 25nt, KL WT vs 20nt, KL WT | 0.0348 | * |
| K | Mean volume per cell | cytoplasm | 15nt, KL WT vs 10nt, KL WT | p<0.0001 | **** |
| K | Mean volume per cell | cytoplasm | 15nt, KL WT vs 20nt, KL WT | p<0.0001 | **** |
| K | Mean volume per cell | cytoplasm | 15nt, KL WT vs 25nt, KL WT | p<0.0001 | **** |
| K | Condensate number per cell | nucleus | 15nt, KL WT vs 10nt, KL WT | p<0.0001 | **** |
| K | Condensate number per cell | nucleus | 20nt, KL WT vs 10nt, KL WT | p<0.0001 | **** |
| K | Condensate number per cell | nucleus | 25nt, KL WT vs 10nt, KL WT | p<0.0001 | **** |
| K | Condensate number per cell | nucleus | 20nt, KL WT vs 15nt, KL WT | p<0.0001 | **** |
| K | Condensate number per cell | nucleus | 25nt, KL WT vs 15nt, KL WT | p<0.0001 | **** |
| K | Condensate number per cell | nucleus | 25nt, KL WT vs 20nt, KL WT | 0.0284 | * |
| K | Condensate number per cell | cytoplasm | 15nt, KL WT vs 10nt, KL WT | p<0.0001 | **** |
| K | Condensate number per cell | cytoplasm | 20nt, KL WT vs 10nt, KL WT | p<0.0001 | **** |
| K | Condensate number per cell | cytoplasm | 25nt, KL WT vs 10nt, KL WT | p<0.0001 | **** |
| K | Condensate number per cell | cytoplasm | 20nt, KL WT vs 15nt, KL WT | 0.0034 | ** |
| L | Mean volume per cell | nucleus | 3 arms vs 5 arms | p<0.0001 | **** |
| L | Mean volume per cell | nucleus | 4 arms vs 5 arms | p<0.0001 | **** |
| L | Mean volume per cell | cytoplasm | 3 arms vs 5 arms | p<0.0001 | **** |

| | | | | | |
|---|---|---|---|---|---|
| L | Mean volume per cell | cytoplasm | 4 arms vs 5 arms | p<0.0001 | **** |
| L | Condensate number per cell | nucleus | 3 arms vs 4 arms | p<0.0001 | **** |
| L | Condensate number per cell | nucleus | 3 arms vs 5 arms | p<0.0001 | **** |
| L | Condensate number per cell | nucleus | 5 arms vs 4 arms | p<0.0001 | **** |
| L | Condensate number per cell | cytoplasm | 3 arms vs 5 arms | p<0.0001 | **** |
| L | Condensate number per cell | nucleus | 5 arms vs 4 arms | 0.0013 | ** |

**Supplementary Table 3. P-values in Figure 3.** Differences among the groups not mentioned in the table are not statistically significant.

| Target Gene | Orientation | Sequence (5'-3') |
|---|---|---|
| ISG15 | Forward | GGCTGGGAGCTGACGGTGAAG |
| | Reverse | GCTCCGCCCGCCAGGCTCTGT |
| IFIT1 | Forward | TTGATGACGATGAAATGCCTGA |
| | Reverse | CAGGTCACCAGACTCCTCAC |
| OASL | Forward | CCATTGTGCCTGCCTACAGAG |
| | Reverse | CTTCAGCTTAGTTGGCCGATG |
| RPS11 | Forward | GCCGAGACTATCTGCACTAC |
| | Reverse | ATGTCCAGCCTCAGAACTTC |
| IFN-β | Forward | GTC AGA GTG GAA ATC CTA AG |
| | Reverse | ACA GCA TCT GCT GGT TGA AG |

**Supplementary Table 4.** RT-qPCR primer sequences.
